# Spatial Coding Dysfunction and Network Instability in the Aging Medial Entorhinal Cortex

**DOI:** 10.1101/2024.04.12.588890

**Authors:** Charlotte S. Herber, Karishma J.B. Pratt, Jeremy M. Shea, Saul A. Villeda, Lisa M. Giocomo

**Author notes:** These authors contributed equally (Karishma J.B. Pratt, Jeremy M. Shea). Corresponding authors: Lisa M. Giocomo, and Charlotte S. Herber,.

## Abstract

Across species, spatial memory declines with age, possibly reflecting altered hippocampal and medial entorhinal cortex (MEC) function. However, the integrity of cellular and network-level spatial coding in aged MEC is unknown. Here, we leveraged *in vivo* electrophysiology to assess MEC function in young, middle-aged, and aged mice navigating virtual environments. In aged grid cells, we observed impaired stabilization of context-specific spatial firing, correlated with spatial memory deficits. Additionally, aged grid networks shifted firing patterns often but with poor alignment to context changes. Aged spatial firing was also unstable in an unchanging environment. In these same mice, we identified 458 genes differentially expressed with age in MEC, 61 of which had expression correlated with spatial firing stability. These genes were enriched among interneurons and related to synaptic transmission. Together, these findings identify coordinated transcriptomic, cellular, and network changes in MEC implicated in impaired spatial memory in aging.

## Introduction

Numerous cognitive domains decline over the human lifespan,^1^ posing a significant challenge to our aging societies.^2^ In particular, declining spatial cognition limits the functional independence of aged individuals, as learning new routes and returning efficiently to remembered locations becomes more difficult.^3–5^ Non-human primates^6^ and rodents^7,8^ also experience age-mediated spatial memory decline. To address this behavioral dysfunction, it is critical to better understand aging-dependent molecular, cellular, and circuit-level changes in the neural systems that support spatial cognition.

Across mammalian species, neural systems in the medial temporal lobe, including the medial entorhinal cortex (MEC) and hippocampus (HPC), are required for spatial memory.^9–11^ The MEC contains grid cells that fire periodically during environmental traversals and hexagonally tile physical space in rodents,^12^ non-human primates,^13^ and humans.^14^ This firing is proposed to provide a map of space that can support path integration.^15,16^ Head direction-,^17^ border-,^18,19^ speed-,^20,21^ and object vector-tuned^22^ cells have also been identified in MEC, providing information regarding an animal’s movement through the environment and sensory features likely relevant to navigation. Additionally, MEC neurons can change their firing rates or shift where their firing fields are active, phenomena collectively referred to as ‘remapping’.^19,23–30^ MEC remapping events often occur in response to changes in task demands and environmental features,^19,24–30^ potentially facilitating the differentiation of distinct contexts. Such remapping in MEC grid cells is likely complemented by place cells^31^ and goal-vector cells^32,33^ in the reciprocally connected HPC, which can also exhibit context-dependent remapping.^34–36^ Collectively, this network of functional cell types across MEC and HPC may provide the necessary neural substrates for an animal to navigate to goals in novel and familiar environments.

Several lines of evidence suggest that MEC-HPC circuit dysfunction contributes to aged spatial memory deficits. Aged humans navigating novel virtual environments take less accurate and more inefficient paths to learned destinations in a manner correlated with reduced activation of medial temporal lobe structures.^37,38^ Both aged humans^38–40^ and rodents^8^ avoid allocentric navigation strategies, suggesting a failure to map novel environments or to retain such a map. This idea is supported by recordings of aged rodent place cells, which are both less stable over days in the same environment^41,42^ and less flexible in novel environments, failing to remap with changing context cues.^43,44^ More indirectly, fMRI experiments uncovered compromised grid-like representations in the MEC that correlated with the extent of path integration errors in aged humans.^45^ Longitudinal structural imaging has also revealed that small EC volume changes are sufficient to predict human spatial memory decline over time.^46^ Notably, neural circuits in the MEC are affected early in forms of aging-mediated cognitive impairment, including preclinical Alzheimer’s disease (AD).^47,48^ Additionally, grid-like representations are impaired in young adults at risk of developing AD due to expression of the APOE-ε4 allele,^49^ and MEC grid coding is degraded in multiple transgenic rodent models of AD.^50–52^ However, changes in MEC spatial coding have yet to be directly investigated in normal aging.

In particular, it is unclear how aging impacts the quality or stability of tuning to navigational variables across MEC functional cell types. The integrity and flexibility of population-level spatial maps in the aged MEC also remains unknown. Since the HPC and MEC are reciprocally connected, one possibility is that spatial coding dysfunction in these regions might interdependently contribute to spatial memory decline in aging. Eventually rejuvenating aged spatial cognition dependent on MEC-HPC networks will also require a more precise understanding of the molecular mechanisms that drive cellular and circuit dysfunction. Towards this goal, mouse brain aging across regions has been comprehensively studied using bulk and single-cell transcriptomic approaches.^53–55^ However, characterizing aging-dependent changes in MEC across phenotypic levels in the same animals would provide unique insight into which genes might drive cellular and circuit dysfunction relevant to behavioral impairment.

Here, we combined *in vivo* silicon probe recordings^57^ with neuronal bulk sequencing in MEC in the same mice, complemented by single nucleus RNA sequencing (snRNA-seq), to identify neural and molecular substrates of aged spatial memory function. Advanced electrophysiologic tools permitted the simultaneous recording of hundreds of neurons per day from each mouse. As a result, we could robustly analyze age effects on MEC spatial coding at the animal level. Moreover, we interrogated how aging altered single neuron firing patterns and population-level spatial coding phenomena. Using a virtual-reality (VR) task with two dynamically interleaved contexts and another with invariant cues,^23^ we demonstrated how aging dually impacts the flexibility and stability of MEC spatial coding at both these levels. Finally, by correlating key spatial coding metrics with the expression of neuronal genes differentially expressed across age groups, we identified potential molecular drivers of aging-mediated spatial cognitive decline in MEC.

## Results

### Experimental strategy to assess spatial memory and entorhinal neural activity in virtual reality

To identify neural correlates of spatial memory impairment in the aging medial entorhinal cortex (MEC), we first recorded MEC neural activity from head-fixed young (mean age ± standard error of the mean [SEM] = 2.58 ± 0.07 months, n = 4 male, 5 female mice), middle-aged (MA, 12.63 ± 0.09 months, n = 5 male, 5 female mice), and aged (22.05 ± 0.021 months, n = 5 male, 5 female mice) C57Bl/6 mice navigating VR environments (Figure 1A). To record neural activity during behavior, we acutely inserted Neuropixels silicon probes^56^ into the MEC for up to six neural recordings per mouse (up to three in each hemisphere) (Methods). Using this approach, we recorded *in vivo* activity from thousands of cells in each age group (n = 15,152 young, 15,011 MA, and 13,225 aged cells).

**Figure 1.**
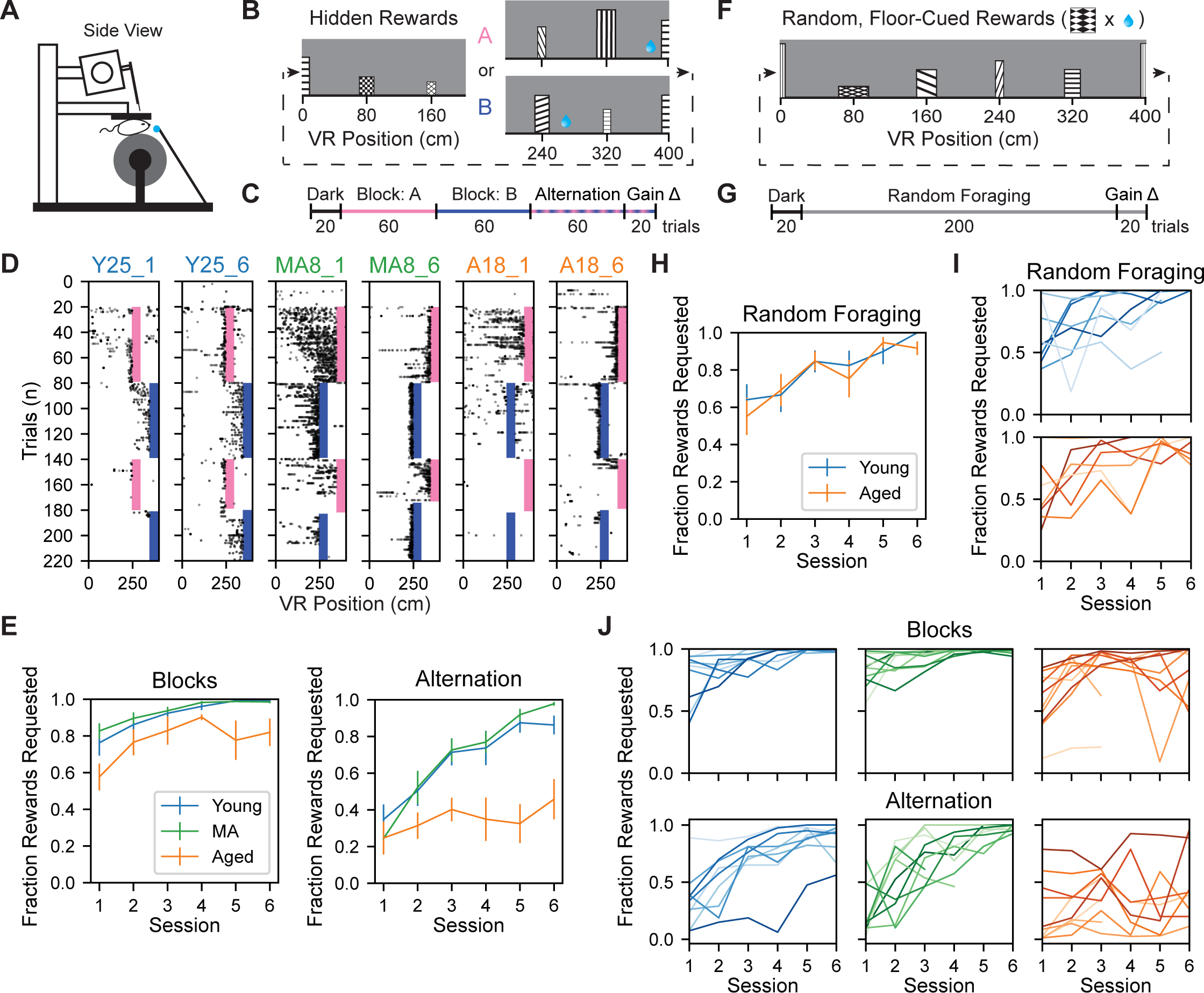
Aged mice exhibit impaired dynamic spatial memory in a VR context discrimination task. **(A)** Schematized acute recording setup. **(B)** Schematized Split Maze (SM) VR track (see also Figure S1A). **(C)** Schematized SM session trial structure. **(D)** Non-consummatory SM lick raster plots with alternation trials sorted by VR context, from a representative young, middle-aged [MA], and aged mouse (left to right). Shading indicates reward zones (pink: context A, dark blue: context B; colors maintained throughout). Subpanel titles indicate mouse and session number. **(E)** Average SM task performance over sessions during the block (top) vs. alternation (bottom) phases by age group (n = 9 young [blue], n = 10 MA [green], & n = 10 aged [orange] mice) (colors maintained throughout). Vertical bars indicate SEM of age group. **(F)** Schematized Random Foraging (RF) VR track (see also Figure S2A). **(G)** Schematized RF session trial structure. **(H)** As in (E), average RF performance over session by age group (n = 8 young, 7 aged). **(I)** Individual RF task learning curves for young (top) and aged (bottom) mice. Each line corresponds to an individual mouse. **(J)** Individual SM learning curves during blocks (top) vs. alternation (bottom) for young, MA, and aged mice (left to right). See also Figure S1 and S2.

To interrogate spatial memory during navigation, mice performed a VR task that contained two hidden reward locations (the Split Maze [SM] task). On each traversal of the 400 cm linear VR track, called a trial, mice could receive water rewards by licking within one of two reward zones, spanning 50 cm (Figure 1B). Each reward location was associated with distinct visual cues (e.g. floor patterns, landmark tower shape) in the second half of the track, termed context A or B (Figures 1B and S1A). To control for the differences in proximity to landmarks, reward-context associations were counterbalanced within each age group. In each session, contexts A and B were presented in successive groups of 60 trials each (“blocks”) and then pseudo-randomly alternated for 80 trials (“alternation”) (Figure 1C). Ten automatic rewards were provided at the beginning of each block to indicate the reward location. After this, accurate context discrimination and licking were required to receive rewards.

By the sixth session, young and MA exhibited licking at the reward zone on block and alternation trials, while aged mice failed to lick consistently in either context during alternation (Figures 1D and S1). To quantify behavioral performance improvements over days while accounting for variance among animals, we fit linear mixed effects models (LMMs) to the fraction of rewards requested during blocks and alternation over sessions, with animal identity as a random effect (see Methods). Block performance improved equivalently across age groups over sessions (Session, β = 0.045, p = 0.004; Session x Aged, β = -0.008, p = 0.709; Aged vs. Young, β = -0.182, p = 0.079). By contrast, alternation performance improved significantly less for aged versus (vs.) young mice (Session, β = 0.106, p = 3.76 x 10^-12^; Session x Aged, β = -0.085, p = 2.70 x 10^-7^; Aged vs. Young, β = -0.292, p = 0.059) (Figure 1E). This indicates an aging-mediated deficit in context discrimination during rapid context alternation.

To control for possible age-mediated differences in the ability to run or motivation to lick, we next implemented a simpler task in a separate group of mice, in which they could lick for rewards at randomly appearing, visually marked zones (Random Foraging [RF]) (Figure 1F).^23^ Other track visual cues were invariant on all trials (see Methods) (Figures 1G and S2A). Using the same electrophysiological approach as in the SM task, we recorded 10,590 cells from young mice (mean age ± SEM = 4.23 ± 0.56 months, n = 5 male, 3 female mice) and 10,228 cells from aged mice (mean age ± SEM = 22.99 ± 0.51 months, n = 2 male, 5 female mice). Across age groups, mice exhibited equivalent behavioral performance in this task (Session, β = 0.064, p = 0.00013; Session x Aged, β = 0.008, p = 0.737; Aged vs. Young, β = -0.018, p = 0.886) (Figure 1H) and had similar learning curves (Figure 1I). This indicated that the motivation and ability to consume rewards in VR remained grossly intact in aged mice. Notably, we observed similar running speed differences across age groups in both the SM and RF tasks (see Figures S1B and S2B). Reward-triggered slowing and licking were also equivalent across age groups in the SM block phase and RF task (see Figures S1E and S2C-G). Therefore, it is unlikely that aged spatial memory deficits in the SM are attributable to impairments in running or licking behavior. Additionally, to exclude the possibility that vision differences impacted context discrimination, all SM mice completed a quantitative visual acuity assessment prior to recording (see Methods) (Figure S1C). The estimated visual acuity thresholds of SM mice did not differ across age groups (Figure S1D). Together, these results indicated spatial memory deficits in aged mice on the SM task, consistent with prior work in aged rodents.^7–8, 41–44^

Lastly, to determine the degree of variability in the aged spatial memory impairment on the SM task, we examined individual behavioral trajectories over SM task experience. While mice showed stereotyped improvement over sessions in the SM blocks within and across age groups, aged mice had more heterogeneous learning curves and failed to improve consistently over days during SM alternation (Figure 1J). We considered to what extent variables other than age might explain these differences. Neither the location of the context A reward (“reward order,” 270 vs. 370 cm context A reward location, β = -0.046, p = 0.673), nor male sex (Male vs. Female, β = - 0.101, p = 0.340) predicted differences in alternation performance (Figure S1G). However, male sex predicted greater aged alternation performance (Aged x Male vs. Aged x Female, β = 0.371, p = 0.015). This raised the question of what neural circuit and molecular differences might underlie the variability in aged spatial cognition, within and across sexes, among these genetically identical mice.

### Single cell spatial coding correlates of heterogeneous spatial memory in aging

We next considered how different functionally-defined MEC cell types in the SM task are impacted by aging.^12,15–22^ We differentiated putative grid from non-grid spatial (NGS) cells using established methods to identify distance tuning during dark running (see Methods).^57–59^ Putative grid cells had peaks in spatial firing autocorrelation on dark trials, which putative NGS cells lacked despite exhibiting spatial tuning on VR trials (Figures 2A-D and S3A-F) (see Methods). Notably, we observed that the density of recorded grid and NGS cells in VR corresponded to that observed in freely moving experiments^58,60–62^ and was unaffected by aging (Figures 2D and S3D). Next, we interrogated the quality and context specificity of spatial tuning across age groups in each cell type.

**Figure 2:**
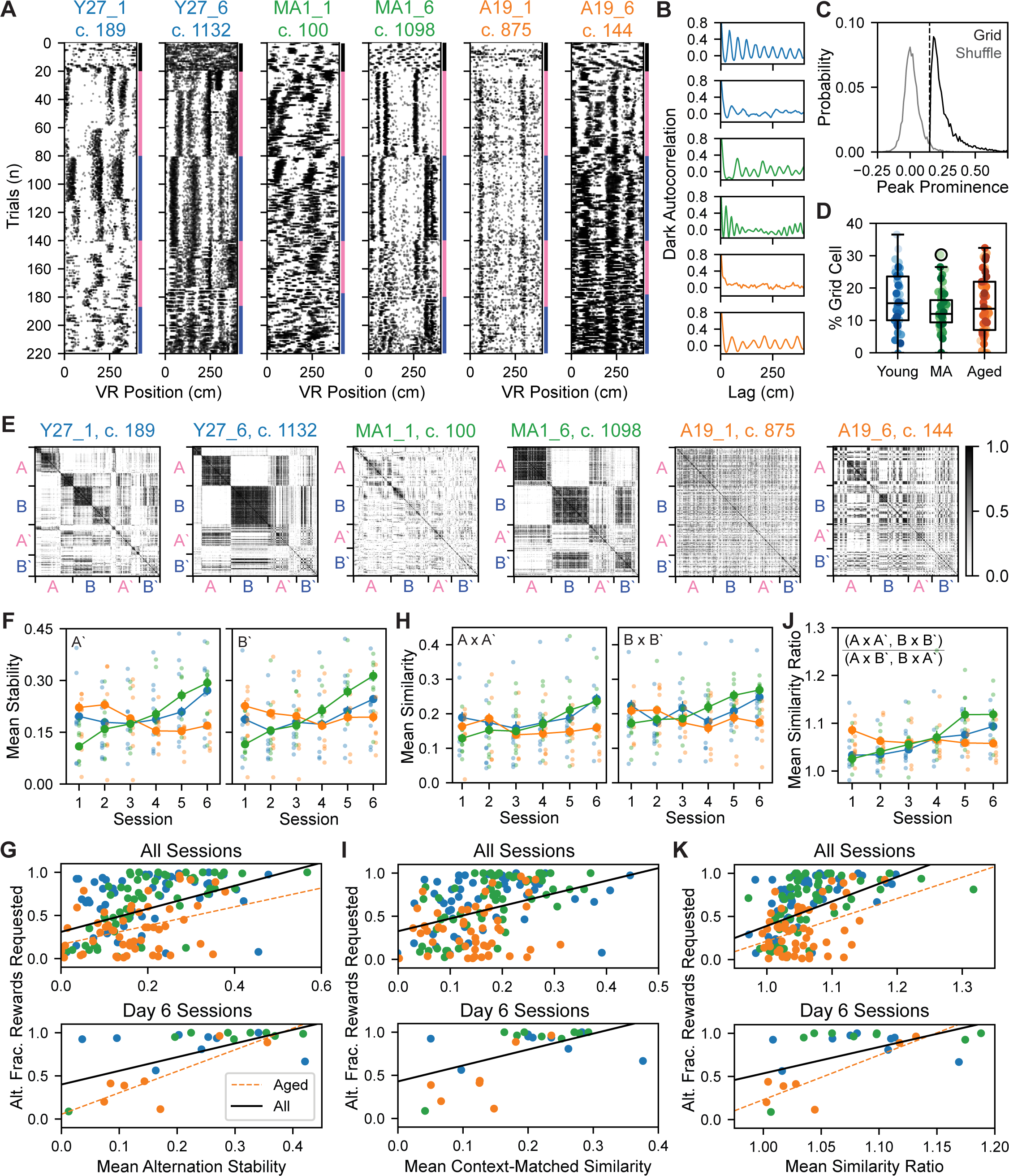
Formation of stable and orthogonal spatial activity patterns across VR contexts relates to dynamic spatial memory. **(A)** Two example SM grid cell raster plots from sessions 1 (left subpanels) and 6 (right subpanels), each from the young, MA, and aged mouse (left to right) closest to the 80th percentile of SM alternation performance in their age group. Dots are individual spikes. Colored right axis indicates trial type, after sorting alternation trials by context (dark [black], context A [pink], and context B [dark blue]). Subpanel titles indicate mouse, session, and cell number (c.). **(B)** Dark trial firing rate (FR) autocorrelations, corresponding to cells in (A) from top to bottom, revealing distance-tuning. This approach was used to identify putative grid cells. **(C)** Probability densities of the prominence of the largest peak in the grid cell dark activity autocorrelation vs. the spike-shuffled activity of those cells at that same lag. Shuffle and real activity measures differed significantly (n = 6,508 model pairs, peak prominence, grid vs. shuffle activity, 0.25 ± 0.0010 vs. 0.01 ± 0.0008, Wilcoxon signed-rank, p < 0.0001). The vertical dashed line indicates the prominence threshold for autocorrelation peaks; 17,498 cells (40.2% SM cells) had a peak in their dark autocorrelation. Among these cells, the 99th percentile of shuffle peak prominences was 0.2017 ± 0.00095. Bin size was 0.01. **(D)** Box and whisker plot of grid cell density by age group (n = 54 young, 58 MA, 55 aged sessions) with outliers indicated by circles. Dots indicate session density values, colored by mouse identity (see Figure S1H for color legend). The box and whiskers indicate the interquartile range (IQR) and 1.5 x IQR, respectively for each age group. The density of grid cells among recorded units for each session did not differ across age groups (% grid cells, young vs. MA vs. aged, 16.17 ± 1.14 vs. 13.38 ± 0.79 vs. 15.05 ± 1.24, Kruskal Wallis test, H = 2.58, p = 0.27). **(E)** Cross-trial correlation matrices corresponding to individual neurons in (A) from left to right, omitting dark and gain change trials and sorting alternation trials by context. Color bar indicates trial-by-trial spatial correlation value. **(F)** Effect of age and session interaction on the mean spatial stability of grid cells during alternation epochs A’ (left) and B’ (right), fitted by separate LMMs. Large dots and vertical bars indicate age group mean and SEM, respectively. Small, pale dots represent LMM-fitted session averages, jittered by age group. **(G)** Alternation performance (Alt. = alternation, Frac. = fraction) vs. mean grid cell stability in each age group (top: n = 53 young, 57 MA, and 53 aged total sessions; bottom: n = 9 young, 9 MA, and 7 aged Day 6 sessions). The black, solid vs. dashed orange lines represent linear regression fits for significant correlations across age groups vs. among aged mice (colors maintained throughout). **(H)** As in (F), for the effect of age and session interaction on the mean similarity of grid cells firing across context-matched task epochs (A x A’ [left], B x B’ [right]), fitted by LMMs separately. **(I)** As in (G) for mean grid cell similarity across context-matched task epochs (top: n = 53 young, 57 MA, & 51 aged sessions; bottom: n = 9 young, 9 MA, & 7 aged sessions). **(J)** As in (F) for the effects of age and session interaction on the mean grid cell similarity ratio (see Methods). **(K)** As in (G) for mean grid cell similarity ratio (top: n = 53 young, 57 MA, & 51 aged sessions; bottom: n = 9 young, 9 MA, & 7 aged sessions). See also Figure S3.

To compare grid and NGS cell spatial firing patterns across SM task epochs (termed in A, B, A’, and B’ in reference to Block A, Block B, Alternation A, and Alternation B trials), we sorted spatially binned firing activity by context on alternation trials. We observed that grid cells exhibited greater context-dependent changes in spatial firing patterns than NGS cells in the SM task (Figures 2A and S3A). To reveal patterns in the similarity of each cell’s spatial firing activity across all trial pairs, we next generated cross-trial correlation matrices for each grid and NGS cell (Methods). By the sixth session, many young and MA grid cell cross-trial correlation matrices displayed a checkerboard structure, reflecting similar spatial firing in context-matched epochs (e.g. A x A’ and B x B’) and dissimilar spatial firing across context-mismatched epochs (e.g. A x B, A x B’, B x A’, A’ x B’) (Figures 2E and S3H). Qualitatively, this correspondence was less common among aged grid cells and among NGS cells from all age groups, which displayed higher spatial firing similarity among epochs. This led us to postulate that context-specific grid cell firing might constitute a neural correlate of the heterogeneity of aged mouse context discrimination.

To address this hypothesis, we interrogated the relationship between alternation performance and three features of spatial firing: (1) stability during each alternation epoch, (2) similarity across context-matched task epochs (e.g. A x A’), and (3) dissimilarity between context-mismatched epochs in grid and NGS cells (e.g. A x B’) (Figures 2F-K and S3I-N). In each epoch for each cell, we computed the moving average pairwise correlation of spatial firing on neighboring trials to assess spatial firing stability (see Methods). Using LMMs to capture variance across cells and animals, we modeled the effect of the interaction of age group and session on grid and NGS cell stability during alternation epochs (A’, B’) (Figures 2F and S3I). In both cell types, we observed that session predicted increasing stability in A’ and B’ in young and MA mice (Session x Young, Session x MA: Grid A’: β = 0.013, β = 0.032; Grid B’: β = 0.014, β = 0.037; NGS A’: β = 0.015, β = 0.033; NGS B’: β = 0.016, β = 0.028) (all p < 0.001). In aged mice, alternation epoch stability decreased or did not change over sessions (Session x Aged: Grid A’: β = -0.016, p < 0.001; Grid B’: β = -0.009, p < 0.001, NGS A’: β = -0.008, p = 0.004; NGS B’: β = 0.004, p = 0.137). Whether we considered all sessions or only the sixth session, the mean alternation (across A’, B’) stability of co-recorded grid cells related to alternation performance across age groups (all sessions: r = 0.45, p = 2.3 x 10^-9^; day 6 sessions: r = 0.62, p = 0.0012) (Figure 2G). Within the aged group, alternation performance and grid cell stability were also correlated (all sessions: r = 0.38, p = 0.0046; day 6 sessions: r = 0.81, p = 0.0275). Similar or weaker relationships were observed between NGS cell stability and performance during alternation (see Figure S3J). Collectively, these findings suggest that aged mice that fail to stabilize spatial firing also tend to fail to improve context discrimination over alternation task experience.

Similar spatial firing on context-matched alternation trials might facilitate the recall of context-associated reward locations. Therefore, we next computed the similarity, or mean pairwise correlation, of grid and NGS cell spatial firing across trials in context-matched epochs (A x A’, B x B’). In particular, we used LMMs to capture age effects on the change in spatial firing similarity over sessions for grid and NGS cells (Figure 2H, S3K). In both cell types, we observed that session predicted increasing similarity between A x A’ and B x B’ in young and MA mice (Session x Young, Session x MA: Grid A x A’: β = 0.019, β = 0.017; Grid B x B’: β = 0.013, β = 0.023; NGS A x A’: β = 0.013, β = 0.019; NGS B x B’: β = 0.009, β = 0.016) (all p < 0.001). By contrast, in aged mice, context-matched epoch similarity decreased or did not change over sessions across cell types (Session x Aged: Grid A x A’: β = -0.002, p = 0.440; Grid B x B’: β = - 0.005, p = 0.060; NGS A x A’: β = 0.006, p = 0.048; NGS B x B’: β = 0.006, p = 0.069). Grid cell context-matched epoch similarity (across both A x A’, B x B’) also related to alternation performance across age groups (all sessions: r = 0.39, p = 3.1 x 10^-7^; day 6 sessions: r = 0.54, p = 0.0060) but not among aged animals alone (all sessions: r = 0.17, p = 0.22; day 6 sessions: r = 0.74, p = 0.060) (Figure 2I). Similar relationships between NGS cell context-matched epoch similarity and alternation performance were also observed (Figure S3L). This suggests that MEC spatial firing similarity across block and alternation trials in the same VR context also relates to recall of learned reward locations during alternation.

To interrogate the role of context-specific grid and NGS firing in context discrimination across groups, we computed a similarity ratio between matched and mismatched task epochs (Methods). Specifically, this ratio compared the average similarity of A x A’ and B x B’ trials to that of A x B’ and B x A’ trials, effectively quantifying the checkerboard structure in the bottom left (or top right corner) of each cross-trial correlation matrix (see Figure 2E). A similarity ratio > 1 implies orthogonal spatial firing across epochs. LMMs revealed that session predicted increasing similarity ratio for grid cells in young and MA but not aged mice (β = 0.013, p < 0.001 vs. β = 0.016, p < 0.001 vs. β = -0.001, p = 0.578) (Figure 2J). By contrast, NGS cell similarity ratio increased for all age groups over sessions (β = 0.009, p < 0.001 vs. β = 0.010, p < 0.001 vs. β = 0.004, p = 0.005) (Figure S3M). This suggests that aged mice fail to orthogonalize grid cell spatial firing patterns over time compared to younger counterparts. Moreover, grid cell similarity ratio related to alternation performance across age groups (all sessions: r = 0.47, p = 2.7 x 10^-10^; day 6 sessions: r = 0.55, p = 0.0043) and among aged mice, especially on the final recording day (all sessions: r = 0.39, p = 0.0036; day 6 sessions: r = 0.86, p = 0.014) (Figure 2K). By contrast, aged alternation performance by the final session was not significantly related to NGS cell similarity ratio (r = 0.64, p = 0.087) (see Figure S3N). Taken together, these results indicate that spatial alternation performance in aging relates to the extent to which grid cells stabilize context-specific firing patterns.

### Network-wide context-aligned remapping dysfunction in aging

We next examined neural correlates of aging spatial memory performance at the population level in MEC. Given the more pronounced context-dependence of grid vs. NGS cell spatial firing activity in the SM task, we focused on the grid cell population. To identify remapping events and their alignment to VR context switches in the SM, we implemented a factorized k-means algorithm to clusters trials with similar network-wide spatial firing activity, termed spatial maps (Methods).^23^ To address the considerable heterogeneity in the structure of trial-by-trial network-wide similarity matrices (Figure 3A), we optimized the k hyperparameter for each session (Methods) (Figures S4A-D). We validated the k optimization procedure on RF spatial cell networks, which have been previously studied in young mice (Figures S4E-F).^23^ After optimization, k-means labeled maps captured the structure in SM and RF network similarity matrices (Figures S4F-G).

**Figure 3:**
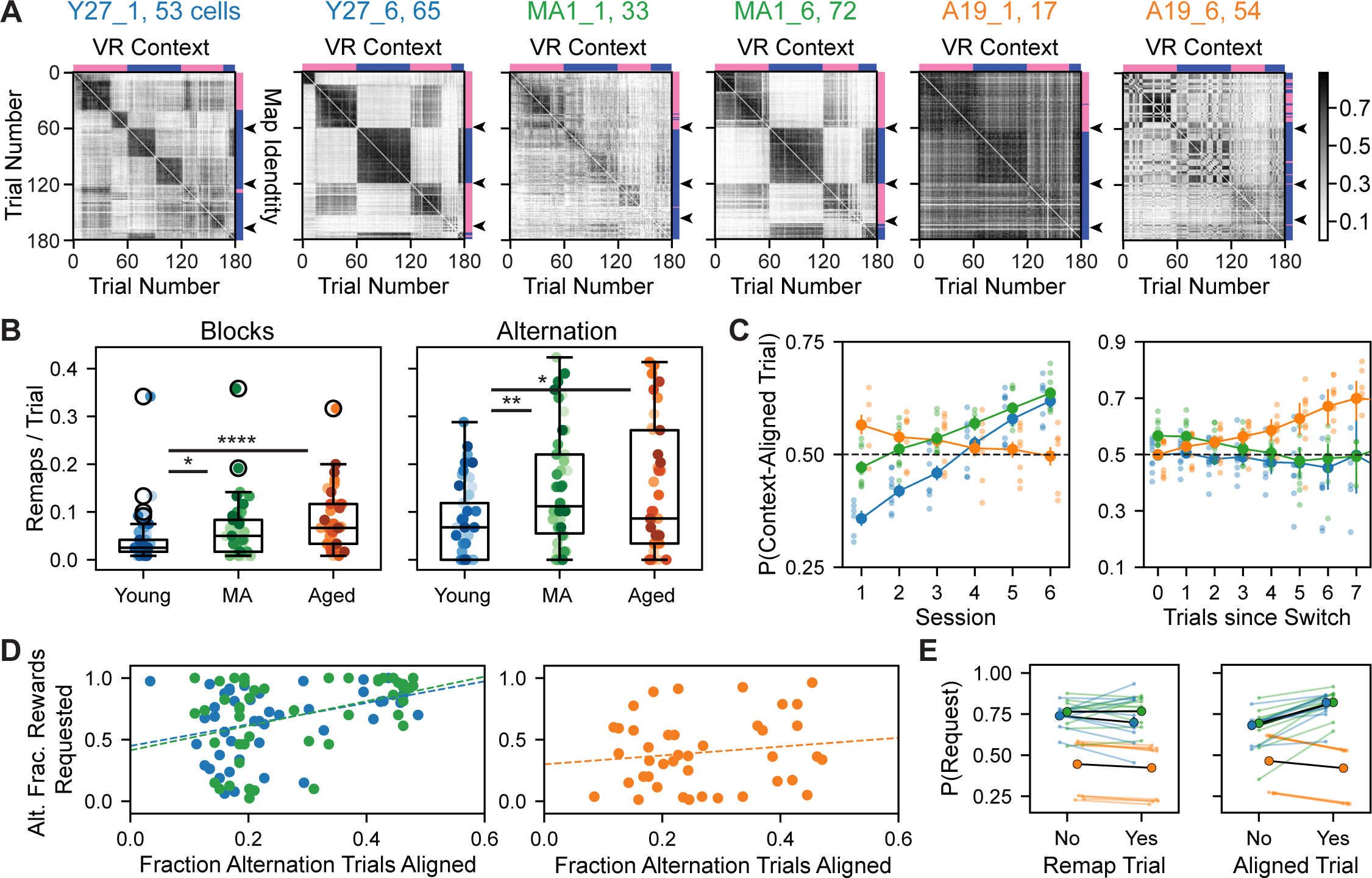
Grid network remapping is more frequent but less aligned to context in aged mice. **(A)** Grid network trial-by-by similarity matrices corresponding to the same example young, MA, and aged sessions (left to right) in Figures 2A-B and E. Dark and gain trials have been omitted, and alternation trials are sorted by VR context (top axis: context A [pink] vs. B [dark blue]). These compare with k-means map identities by trial (right axis: same color schema) (see Methods). Arrowheads (right axis) point out map identity transitions expected at VR context switches. Subpanel titles indicate mouse, session, and network grid cell count. Color bar indicates trial-by-trial spatial correlation. **(B)** Box and whisker plot, plotted as in Figure 2D, of remapping frequency (number of remaps per trial), which differed across age group (n = 47 young, 46 MA, and 41 aged sessions) in the block phase (left: remaps / trial, young vs. MA vs. aged, 0.0409 ± 0.0075 vs. 0.063 ± 0.0099 vs. 0.0792 ± 0.0096; Kruskal-Wallis test, H = 16.5, p = 0.00025, post-hoc Conover test, young vs. MA, p = 0.033; young vs. aged, p = 0.00010; MA vs. aged, p = 0.056) and during alternation (right: 0.0758 ± 0.0116 vs. 0.144 ± 0.0174 vs. 0.1529 ± 0.0218; H = 10.2, p = 0.0060, young vs. MA, p = 0.010; young vs. aged, p = 0.016; MA vs. aged, p = 0.84). **(C)** The effect of the interaction of age and session (left) and age and trials since a pseudorandom context switch (right) on the probability (P) of an aligned k-means map identity and VR context on alternation trials (context-aligned trial), modeled by logistic regression (n = 8079 trials, pseudo R^2^ = 0.0853, LLR p = 1.466 x 10^-194^). Large dots and vertical bars indicate age group mean and SEM fitted probability, respectively. Small, pale dots represent fitted session averages, jittered by age group. At right, we required data from at least three mice per group to plot a given x-value. Dotted horizontal lines indicate chance level alignment. **(D)** Alternation (alt.) performance (fraction [frac.] rewards requested) vs. fraction (frac.) of alternation trials for which k-means map identity and VR context are aligned for young and MA (left, n = 46 young, 47 MA sessions) vs. aged sessions (right, 41 aged sessions). **(E)** The effect of the interaction of age and remapping (left) and age and spatial map-context alignment (right) on the probability of an alternation reward request, as modeled by logistic regression (n = 8079 trials, pseudo R^2^ = 0.2098, LLR p < 0.0001). Bold dots and black lines indicate mean fitted age group probability. Light dots and lines indicate mean fitted animal probability, colored and jittered by age group. See also Figure S4 and S5.

To assess the flexibility of grid networks in aging, we first computed the frequency of remaps (the number of transitions between spatial maps/the number of trials in each task phase). In both SM task phases, MA and aged grid networks remapped more than young counterparts (young vs. MA vs. aged, block: 0.0409 ± 0.0075 vs. 0.063 ± 0.0099 vs. 0.0792 ± 0.0096, Kruskal-Wallis test, H = 16.5, p = 0.00025; alternation: 0.0758 ± 0.0116 vs. 0.144 ± 0.0174 vs. 0.1529 ± 0.0218; H = 10.2, p = 0.0060) (Figure 3B). As expected, remaps were more frequent in the SM vs. the RF task, in which no context changes occurred and networks in both age groups remapped similarly rarely (Figure S4H).

To examine the alignment of spatial maps and VR contexts transitions, we imposed context identities on labeled maps based on their occupancy of and similarity to network activity during A and B epochs (see Methods). By the sixth session, young and middle-aged grid networks more often exhibited transitions in map identity near VR context switches, but this was less common among aged grid networks (see right axes of Figure 3A). We next implemented a logistic regression model to quantify age and session effects on the probability that a given alternation trial had aligned spatial map and VR context identity (Methods) (Figure 3C). For young and MA grid networks, each session predicted a 28% and 15% greater chance of an aligned trial (Young, Odds Ratio (OR) = 1.28, p = 2.63 x 10^-22^; MA, Or = 1.15, p = 7.67 x 10^-10^). As such, the typical probability of map-context alignment exceeded random chance levels (50% for a given alternation trial) starting on day 4. By contrast, the probability of map-context alignment on aged alternation trials did not increase over sessions, remaining near chance level (Aged, Or = 0.95, p = 0.043). The single predictor of above chance map-context alignment for aged trials was the number of consecutive trials since a pseudorandom context switch. With each additional trial in the same context, the probability of aged, but not young or MA, map-context alignment increased by 9% (Aged, Or = 1.09, p = 0.0043; Young, Or = 0.98, p = 0.47; MA, Or = 0.94, p = 0.0388). This raises the possibility that aged spatial map context uniquely reflects how recently an animal experienced that context.

Importantly, we determined that the discreteness and coordination of remapping events was comparable across age groups (Methods, see Figure S5A-D). We also confirmed that positional information remains distinct across spatial maps in aging (Methods, see Figures S5E - H). Additionally, we accounted for age differences in the timing of context recognition on each trial by performing population analyses using only grid network activity in the back of the track (Methods, see Figures S5I-J). Collectively, these control measures ensured that age differences in map-context alignment were not attributable to differences in the nature of remapping.

Lastly, we assessed whether grid network remapping or map-context alignment related to spatial memory. While remapping frequency during alternation did not correlate with alternation performance in any age group (Figure S4I), we found that the fraction of aligned alternation trials correlated with alternation performance among young and MA, but not aged, sessions (young: r = 0.38, p = 0.0082; MA: MA, r = 0.41, p = 0.0043; aged: r = 0.055, p = 0.73) (Figure 3D). We used logistic regression to then assess whether the occurrence of a remap or map-context alignment predicted performance, measured by the probability of a reward request, on individual alternation trials (Figure 3E). We found that a remap did not predict increased performance in any age group (Young, Odds Ratio [OR] = 0.84, p = 0.31; MA, OR = 1.17, p = 0.25; Aged, OR = 0.90, p = 0.39). However, echoing results at the session level, trial alignment predicted a 52% and 95% greater chance of a reward request on young and MA, but not aged, alternation trials (Young, Odds Ratio (OR) = 1.52, p = 6.74 x 10^-5^; MA, OR = 1.95, p = 2.98 x 10^-10^; Aged, OR = 0.69, p = 5.92 x 10^-5^). Taken together, these findings suggest that aged grid networks exhibit more frequent but, ultimately, dysfunctional remapping that does not yield better spatial map alignment to changing contexts over sessions. Moreover, the relationship between map-context alignment in MEC and rapid context discrimination is lost at the session and trial levels in aging, raising the possibility that the dysfunctional MEC remapping might contribute to impaired spatial memory.

### Reduced LFP power during running in middle aged and aged mice

We next sought to uncover the impact of aging on MEC network-level oscillatory activity as measured by the local field potential (LFP). Incorporating new information into MEC spatial maps at fast time scales may require temporal coordination of neuronal activity, facilitated by theta frequency (6 - 12 Hz) oscillations in the LFP.^63,64^ Additionally, gamma frequency (slow: 20 - 50 Hz; fast: 50 - 110 Hz) oscillations in MEC LFP are critical for communication with the hippocampus and support spatial learning.^65^

We observed qualitative decreases in LFP power in theta and gamma frequency bands in MA and aged sessions compared to young ones (Figure 4A). This was also apparent in the mean power spectral density, pooled across sessions in each age group (Figure 4B). To quantitatively compare the power of theta and gamma rhythms across age groups, we first controlled for observed differences in running speed distributions across age groups (see Figures S1B and S2B), as theta dynamics in MEC are influenced by running speed and acceleration.^66,67^ In particular, we sampled MEC LFP power in each session from both tasks by in a speed range (20 - 40 cm/s) that collapsed running speed differences across age groups (n = 98 young, 58 MA, 97 aged sessions, mean running speed, young vs. MA vs. aged sessions, 29.93 ± 0.12 vs. 29.6 ± 0.13 vs. 29.39 ± 0.22, Kruskal-Wallis test, H = 2.85, p = 0.24; peak running speed, 40.0 ± 0.0 vs. 40.0 ± 0.0, 39.86 ± 0.07, H = 3.05, p = 0.22). Indeed, starting in middle age, mean theta (young vs. MA vs. aged power [dB], 43.66 ± 3.56 vs. 30.41 ± 3.94 vs. 27.29 ± 2.40, Kruskal-Wallis test, H = 14.29, p = 0.00079, post-hoc Conover test, young vs. MA, p = 0.0086, young vs. aged, p = 0.0012), slow gamma (1.46 ± 0.09 ; 0.94 ± 0.08 ; 0.92 ± 0.06, H = 21.63, p = 2.01 x 10^-5,^ young vs. MA, p = 0.00044, young vs. aged, p = 0.000052), and fast gamma power (0.57 ± 0.04 vs. 0.39 ± 0.05 vs. 0.35 ± 0.02, H = 23.44, p = 8.13 x 10^-6^ ; young vs. MA, p = 0.00024, young vs. aged, p = 0.000021) decreased (Figure 4C). These results raise the possibility that the rapid incorporation of information into spatial maps supported by theta rhythm, as well as the coordination of spatial coding across the MEC-HPC circuit by gamma rhythms, may be impaired over healthy aging.

**Figure 4.**
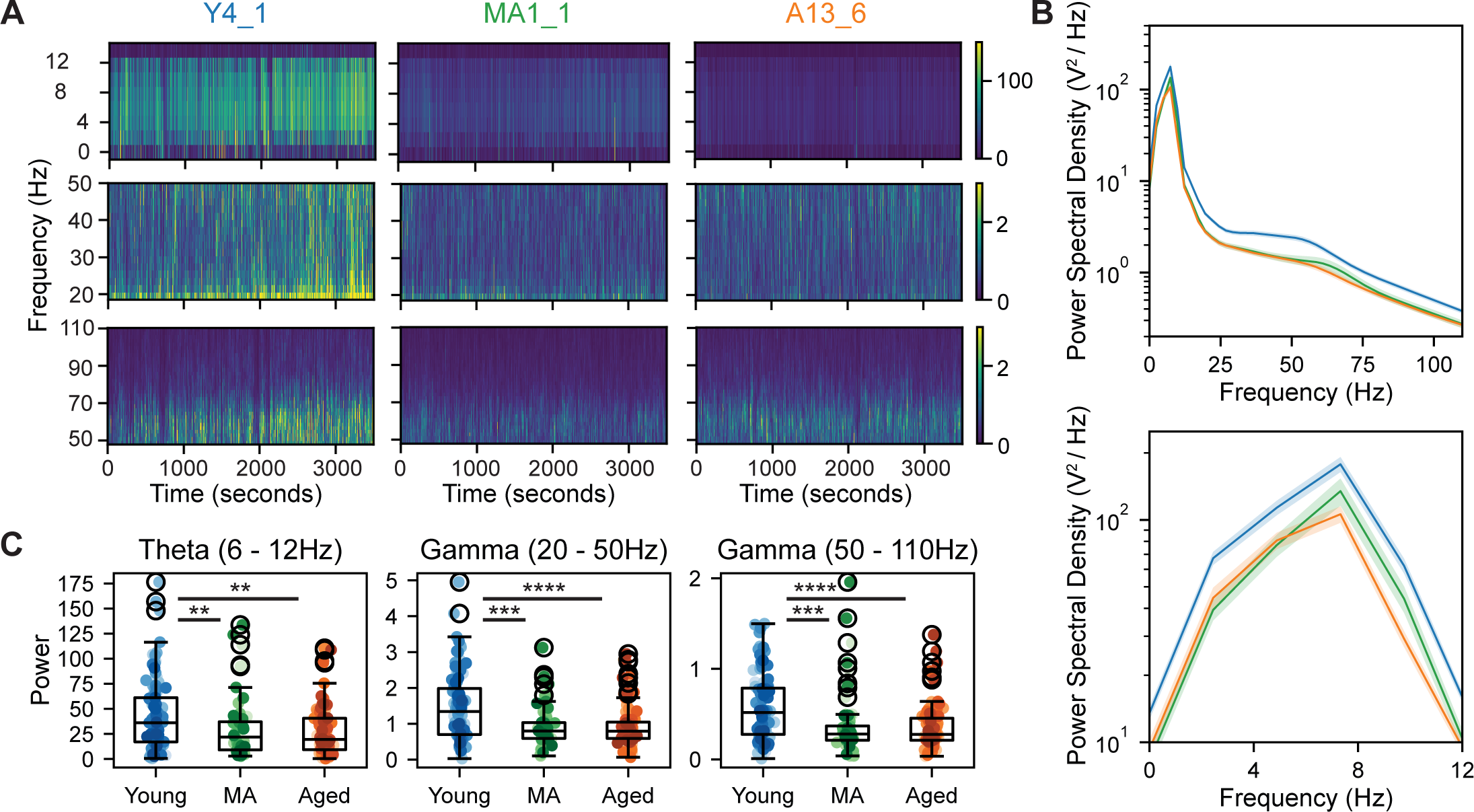
Theta, slow & fast gamma power during running is diminished in aged mice. **(A)** Example spectrograms in the theta (top row; 6-12 Hz), slow gamma (middle; 20-50 Hz), and fast gamma (bottom; 50-110 Hz) frequency bands for representative young, MA, and aged mice (left to right columns) for 3500 seconds of recording. Color coded for minimum (blue) and maximum (yellow) values. **(B)** Mean power spectral density across age groups (top: all frequency bands in (A); bottom: theta frequency range) with SEM shaded. **(C)** Box and whisker plot of mean LFP power in the theta, slow gamma, and fast gamma (left to right) frequency bands during running across age groups in both tasks (n = 98 young, 58 MA, 97 aged sessions). Dots indicate session power values, colored by mouse identity.

### Aged spatial coding instability in an invariant VR environment

Given the dysfunction we observed in aged single cell and network-level spatial coding in the dynamic SM task, we next examined the stability of aged spatial coding in the invariant RF task. As some RF sessions lacked dark trials, we considered all spatial cells (grid and NGS) together (Methods) (Figure 5A-D). Consistent with prior works, spatial cells in young mice exhibited stable spatial firing patterns across neighboring RF trials (Figure 5A), producing smooth, periodic trial-averaged spatial tuning curves.^23,69^ By contrast, in aged mice, we frequently observed spatial cells with drift in firing field locations across neighboring trials and less smooth tuning curves. Consistent with the SM task, spatial coding by the aged MEC was quantifiably degraded during RF sessions. Averaging across co-recorded spatial cells, we observed decreased spatial firing coherence (young vs. aged sessions, 0.7425 ± 0.0108 vs. 0.6480 ± 0.0136, Wilcoxon rank sum test, p = 4.526 x 10^-7^) (Figure 5E) and spatial information score with age (0.0928 ± 0.0081 vs. 0.0636 ± 0.0056, p = 0.00323) (Figure 5F). Moreover, aged sessions exhibited reduced mean spatial cell within map stability compared to young sessions (young vs. aged, 0.1314 ± 0.0087 vs. 0.0895 ± 0.0075, Wilcoxon rank sum test, p = 0.00026) (Figure 5G). These findings reveal that the spatial firing stability of aged spatial cells is significantly impaired even in unchanging environments.

**Figure 5.**
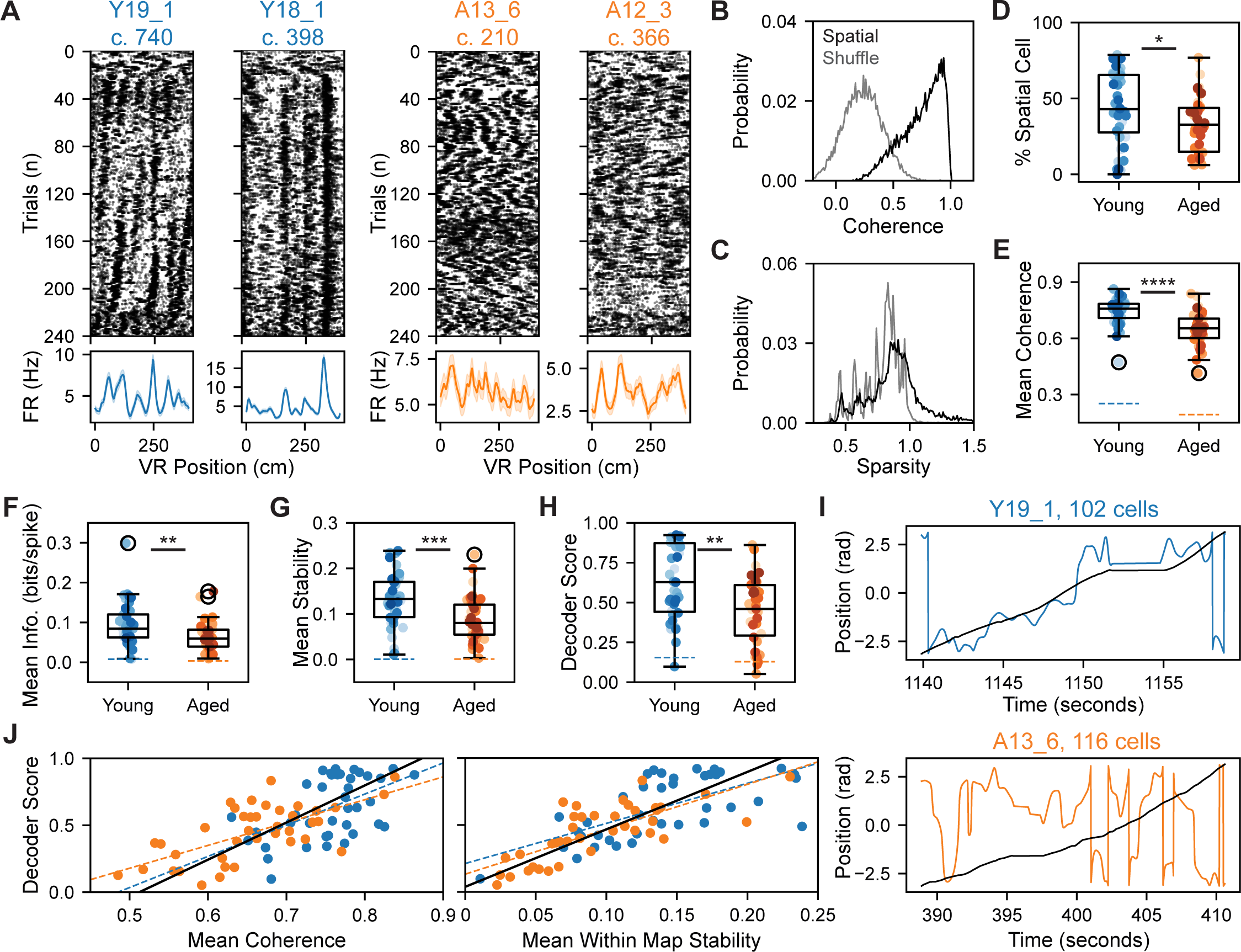
Aged mice in stable VR environments exhibit spatial cell instability that impairs position decoding. **(A)** Four example RF spatial cell raster plots from young and aged mice (top: left, right) and corresponding trial-averaged spatial tuning curves (bottom: shaded indicates SEM). The first 20 trials were run in the dark, and the last 20 trials were gain-manipulated. Plotted and labeled as in Figure 2A. **(B)** Probability density of spatial cell vs. shuffle activity spatial firing coherence. The spatial firing coherence of classified spatial cells’ activity significantly exceeds that of their shuffled activity, pooling cells across age groups (n = 8,476 model pairs, spatial cell vs. shuffle, 0.7391 ± 0.0019 vs. 0.2306 ± 0.0018, Wilcoxon signed-rank test, p < 0.0001). The 99th percentile of shuffle coherence was 0.4440 ± 0.0010 (n = 20,928 total RF cells). Bin size was 0.01. **(C)** As in (B) for the spatial firing sparsity of classified spatial cell activity vs. shuffled activity across age groups (0.9154 ± 0.0037 vs. 0.7716 ± 0.0016, p < 0.0001). Classified spatial cells have both significantly coherent and sparse spatial firing (see Methods). The mean 99th percentile of sparsity for 100 shuffles per spatial cell was 0.8021 ± 0.0022 (n = 20,928 total RF cells). Bin size was 0.01. **(D)** Box and whisker plot, plotted as in Figure 2D, for spatial cell density in RF sessions (n = 44 young, 42 aged sessions with spatial cells). Spatial cell density decreased with age (% spatial cells, young vs. aged, 44.29 ± 3.52 vs. 32.37 ± 2.79, Wilcoxon rank sum test, p = 0.0136). Dots indicate session values, colored by mouse identity (see Figure S1C for legend). **(E)** Mean spatial cell coherence decreased among aged sessions (n = 43 young, 42 aged sessions). Dashed lines indicate mean shuffle score for each age group. Plotted as in (D) except that dots indicate session mean values across cells. **(F)** As in (E), for spatial cell spatial information (info). **(G)** As in (E), for spatial cell within-map stability. **(H)** As in (D), decoder performance for models trained and tested on aged vs. young was lower (score 0 = chance, score 1 = perfect; n = 39 young, 39 aged sessions). **(I)** Example decoder performance on single trials from representative young (top) and aged (bottom) RF sessions, corresponding to the first raster from each age group in (A). Lines indicate VR (black) vs. decoded position (colored). **(J)** Session decoder score vs. mean spatial cell coherence (left) and mean spatial cell stability (right). Decoder score relates to mean spatial cell spatial coherence across (r = 0.98, p = 1.1 x 10^-54^) and within age groups (young, r = 0.53, p = 0.00052; aged, r = 0.65, p = 5.2 x 10^-6^). Decoder score relates to mean spatial cell within map stability across (r = 0.98, p = 4.2 x 10^-55^) and within age groups (young, r = 0.67, p = 3.5 x 10^-6^; aged, r = 0.76, p = 2.1 x 10^-8^).

Since MEC neurons project to multiple brain regions that process spatial information,^69^ we next considered to what extent network-level positional information output by MEC would be degraded by spatial cell instability in aging. We fit circular-linear regression models, termed decoders, to estimate animal position from spatial cell network activity (Methods) (Figure 5H-J). We found that decoder performance decreased when trained and tested on spatial cell activity from aged vs. young sessions (young vs. aged, 0.6307 ± 0.0369 vs. 0.448 ± 0.0333, Wilcoxon rank sum test, p = 0.00185) (Figure 5H; see also Figure S5E). Sample decodes on single trials revealed the greater number and severity of position estimation errors for aged compared to young sessions (Figure 5I). Moreover, session decoder score was highly correlated with mean spatial cell spatial coherence (r = 0.98, p = 1.1 x 10^-54^) and within map stability (r = 0.98, p = 4.2 x 10^-55^) (Figure 5J). Collectively, these results suggest that aged spatial cell instability is strongly associated with degraded positional information output from the MEC network.

### Network-wide increased speed gain and speed-tuning instability in aging

Given the degradation of position coding by the aged MEC, we next considered how the coding of other navigational variables, in particular speed, might also change. To address this, we identified positively and negatively modulated speed-tuned (+ vs. - speed cells) (Methods) (Figure 6A-C).^20,58^ The density of speed-tuned cells was unchanged by aging (see Figure S6). Next, we compared the gain, or sensitivity of, speed cells to changes in animal speed (Methods). In both + and - speed cells, we observed more gain to speed in aged vs. young sessions (+ speed cells: young vs. MA vs. aged, 0.0445 ± 0.0015 vs. 0.0508 ± 0.0027 vs. 0.0556 ± 0.0026, Kruskal-Wallis test, H = 15.15, p = 0.0005; post-hoc Conover test, young vs. MA, p = 0.1111, young vs. aged, p = 0.0003; - speed cells: -0.0366 ± 0.0024 vs. -0.0432 ± 0.004 vs. -0.05 ± 0.0026, H = 24.33, p = 5.221 x 10^-6^; young vs. MA, p = 0.0816, young vs. aged, p = 0.000002) (Figure 6D). Trending differences between young and MA sessions imply that this change, like decreased LFP power, begins earlier than 21 months of age. In addition to altered speed tuning gain, + speed cells from aged sessions demonstrated decreased speed tuning stability over trials (0.1905 ± 0.0063 vs. 0.2143 ± 0.0061 vs. 0.1693 ± 0.0047, Kruskal-Wallis test, H = 29.28, p = 3.96 x 10^-7^; post-hoc Conover test, young vs. MA, p = 0.00080, young vs. aged, p = 0.013) (Figure 6E) (Methods). We also recapitulated these differences in control groups of + and - “speed only cells,” from which we excluded any cells that also qualified as spatial cells (Figures S6F-G). Taken together, these results uncovered increased speed tuning sensitivity and instability in aging that may be implicated in spatial map instability.

**Figure 6.**
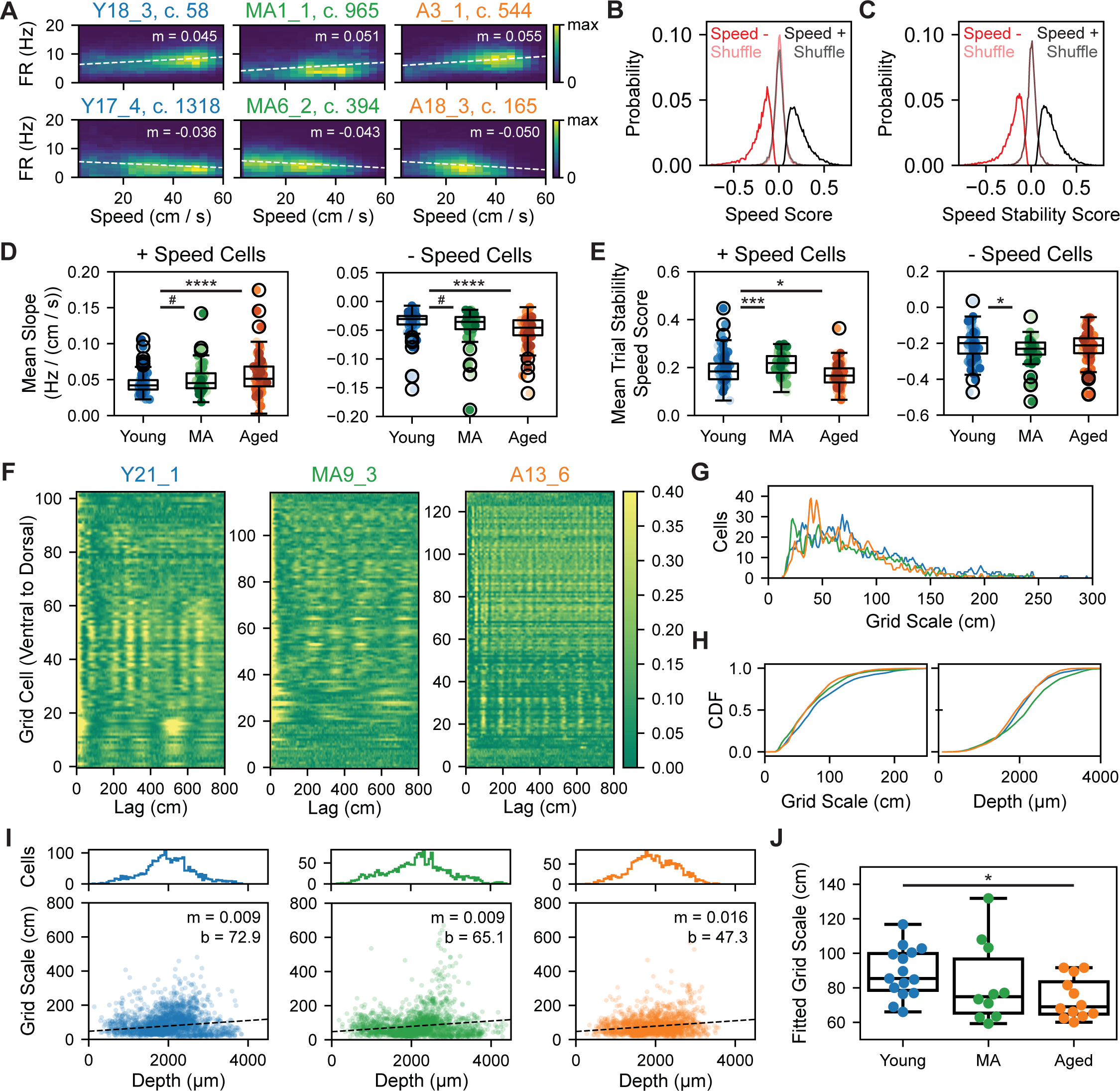
Increased speed gain and speed-tuning instability accompany grid scale compression in aging. (A) Double histograms of speed and firing rate (FR) of + (top) and - (bottom) speed cells from young, MA, and aged (left to right) mice. Dashed white line indicates regression fit (m = FR-speed slope). Color bar indicates occupancy in 0.02 second time bins. Subpanel title indicates mouse, session, and cell number. **(B)** Probability density of speed scores of + (black) and - (red) speed cells vs. shuffled activity. Classified + and - speed cells had speed scores whose absolute value exceeded that of speed scores generated from their shuffled activity, pooling cells across age groups (n = 17,091 model pairs, + speed cell vs. shuffle speed scores, 0.2204 ± 0.0009 vs. -0.005 ± 0.0005, Wilcoxon signed-rank test, p < 0.0001; n = 8,068 model pairs, - speed cell vs. shuffle, - 0.2035 ± 0.0013 vs. 0.0001 ± 0.0006, p < 0.0001) (see Methods). The 99th percentile of shuffle speed scores was 0.1194 ± 0.0002 (n = 64,316 cells across tasks). Bin size was 0.01. **(C)** As in (B) for speed stability scores. Classified + and - speed cells also had speed stability scores whose absolute value significantly exceeded those generated from their shuffled activity (+ speed cell vs. shuffle speed stability scores, 0.2181 ± 0.0008 vs. -0.001 ± 0.0005, p < 0.0001; - speed cell vs. shuffle, -0.2049 ± 0.0013 vs. -0.002 ± 0.0006, p < 0.0001). The 99th percentile of shuffle speed stability scores was 0.1239 ± 0.0002. **(D)** Box and whisker plot of mean FR-speed slope of + speed cells (left) and - speed cells (right) across age groups in both tasks (n = 96 young, 58 MA, 96 aged sessions with + speed cells; n = 96 young, 58 MA, 94 aged sessions with - speed cells). Dots indicate session mean values across speed cells, colored by mouse identity. **(E)** As in (D) for mean trial stability speed score. - speed cell trial stability did not change systematically with age (-0.2125 ± 0.0081 vs. -0.2393 ± 0.0098 vs. -0.2165 ± 0.0082, H = 6.41, p = 0.040; young vs. MA, p = 0.040, MA vs. aged, p = 0.09, young vs. aged, p = 0.59). **(F)** Heatmaps of dark trial FR autocorrelation for all grid cells in young, MA, and aged (left to right) example sessions, with cells sorted by distance from the probe tip (ventral [bottom] to dorsal [top]). Color bar indicates autocorrelation value. **(G)** Ηistogram of estimated grid cell scales by age group (n = 2,441 young, 2035 MA, and 2032 aged cells). Bin size was 1 cm. **(H)** Cumulative density of grid cell scales (left) and recorded depths (right) by age group. The distributions of grid scale differed by age (two- sided Kolmogorov-Smirnov test, aged vs. young, stat = 0.12, p = 6.0 x 10^-14^, sign = 1, loc = 100; MA vs. young, stat = 0.085, p = 1.7 x 10^-7^, sign = 1, loc = 54; aged vs. MA, stat = 0.076, p = 1.5 x 10^-5^, sign = -1, loc = 35), as did those for grid cell depth (aged vs. young, stat = 0.057, p = 0.0014, sign = 1, loc = 1795; MA vs. young, stat = 0.14, p = 1.5 x 10^-20^, sign = -1, loc = 2138; aged vs. MA, stat = 0.18, p = 3.5 x 10^-30^, sign = 1, loc = 2105). **(I)** Histograms of grid cell depths (top) and scatter of grid cell scale and depth (bottom) by age group (young, MA, aged, left to right; linear regression fit, dashed black line with slope and intercept labeled). **(J)** Box and whisker plot of LMM-fitted grid scale across age groups (see Methods), indicating decreased grid scale with age accounting for depth. Dots indicate LMM-fitted animal averages. See also Figure S6.

MEC speed coding is also supported by local interneurons (INs) and conjunctive speed-tuned grid cells.^17,70,71^ To interrogate aged speed coding in these populations, we separated recorded putative fast-spiking INs from excitatory cells (Methods, Figures S6A-C), noting an increase in the density of recorded INs and speed-tuned INs in MA and aged vs. young mice (Figures S6D-E). Consistent with our observations in + speed cells, we also observed increased speed gain and speed-tuning instability in positively modulated speed-tuned INs and conjunctive speed-tuned grid cells (Figures S6F-G), revealing that aging produces aligned changes in speed coding across cell types.

### Grid scale compression in aging

Speed gain and grid scale change proportionally when environmental dimensions are manipulated,^30^ consistent with a continuous attractor network model framework of entorhinal activity.^72,73^ This suggested that grid scale compression could co-occur with increased speed gain in aging. To estimate grid scale (i.e. the distance between grid firing nodes) in 1D VR, we identified the location of the first peak in each grid cells’ spatial firing autocorrelation on dark trials (Methods).^57^ In young and middle-aged sessions, we observed an increase in grid scale from dorsal to ventral MEC, consistent with prior work (Figure 6F).^57^ However, aged grid cell autocorrelation peaks were commonly closer together, indicating smaller grid spacing, with weaker gradients from dorsal to ventral recording sites.

Consistent with observations of grid scale in the autocorrelations (Figure 6F), comparing grid scale distributions across age groups revealed a peak corresponding to smaller aged grid scales (Figure 6G) and a left-ward shift of grid scale cumulative density for aged and MA vs. young cells (Figure 6H). To control for the relationship between grid scale and dorsal-ventral location in MEC,^12,17,74^ we also compared the cumulative density of estimated recording depths along the probe axis across age groups (Methods). There were significant differences in depth sampling across age groups (Figure 6H), so we fit the linear relationship between grid cell depth and scale within each age group via linear regression (Figure 6I). Indicative of grid scale compression, the slope of grid cell depth and scale increased 78% (m = 0.016 vs. m = 0.009), and the y-intercept decreased 35% among aged vs. young mice (b = 47.3 vs. b = 72.9). Moreover, we used LMMs to model grid scale as a function of age group and unit depth accounting for cell and animal variance (Methods). Accounting for the effect of unit depth (β = 0.009, p = 1.28 x 10^-20^), being aged predicted decreased grid scale (Aged vs. Young, β = -15.255, p = 0.030) (Figure 6J). These results raise the possibility that grid scale changes in the aged MEC may influence the scale or stability of place fields in the aged HPC,^41–44^ given their known interdependence in young animals.^62^ Ultimately, altered long-term place map stability might further contribute to impaired spatial learning and memory in aging via these changes in entorhinal inputs.

### Transcriptomic correlates of spatial coding dysfunction in aged MEC neurons

To further investigate the molecular and cellular correlates of the MEC spatial coding dysfunction uncovered in healthy aging, we performed bulk RNA sequencing of MEC neuronal nuclei isolated from the hemispheres of the same young and aged mice that completed the RF task (Figures 7A and S7A) (see Methods). By comparing neuronal gene expression across age groups, we identified 458 genes that were differentially expressed (DEGs) with age in MEC. The expression of 172 genes increased while the expression of 286 genes decreased with age in MEC (absolute fold change [FC] ≥ 0.5, mean expression value ≥ 25, and adjusted p-value [p-adjusted] ≤ 0.05) (Figure 7B). A heatmap of the top 100 DEGs with age revealed consistent differential gene expression across mice within each age group (Figure S7B). To classify the role of aging DEGs in biological processes, we performed gene ontology and gene set enrichment analysis (see Methods). We found that pathways regulating synaptic transmission, cell communication and cell signaling were significantly enriched among aging DEGs (Figure 7C). Furthermore, the two top annotated gene sets among aging DEGs were pathways related to reactive oxygen species and oxidative phosphorylation (Figure 7D).

**Figure 7.**
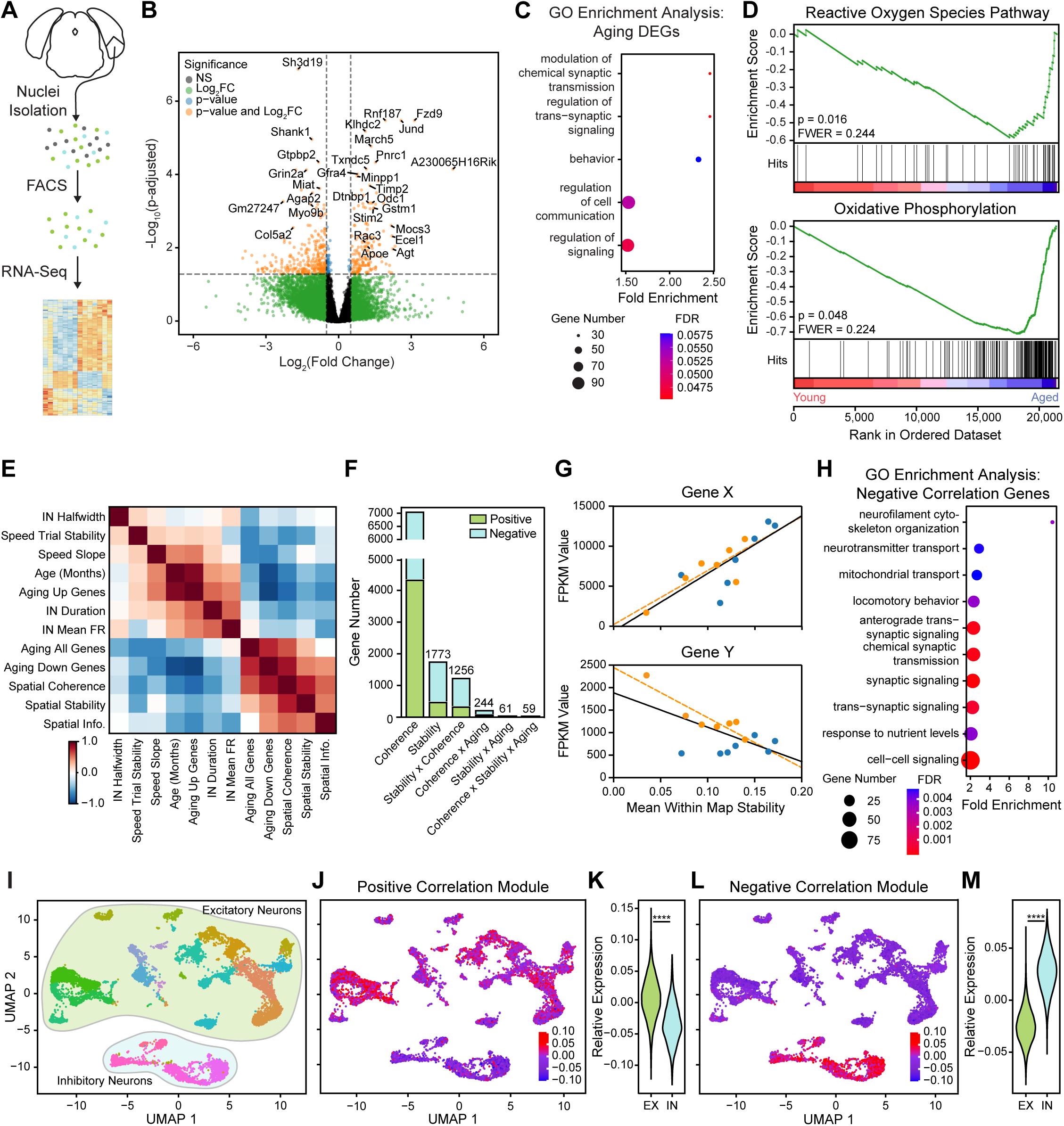
Transcriptomic correlates of spatial coding dysfunction in the aged MEC. **A)** Schematized neuronal nuclei isolation and RNA-seq library generation from MEC, dissected from hemispheres at ± 3.1 - 3.5mm lateral and 4.75 - 5.25mm posterior to bregma below 3.5mm from dorsal brain surface (see Methods). **(B)** Volcano plot of differential expression between young and aged MEC neuronal nuclei (n = 7 young, 7 aged mice; Wald test followed by Benjamini and Hochberg multiple hypothesis correction). 458 significant differentially expressed genes (DEGs) are indicated by orange dots. **(C)** Gene ontology (GO) enrichment analysis of biological processes on DEGs between young and aged MEC neuronal nuclei (Fisher’s Exact test followed by False Discovery Rate [FDR] calculation). **(D)** Top annotated gene sets correlated with age by family-wise error rate (FWER) using gene set enrichment analysis (permutation test followed by FWER correction; 6 gene sets were significant at p < 0.05; 2 gene sets were significant at FWER < 0.25). The running sum of the enrichment score is displayed in green, along with the gene set members placement (hits) among dataset wide expression changes as black vertical bars. Dataset genes are ranked in order of expression change from young to old (left to right). Gene sets are annotated with their p-values and FWERs. **(E)** Correlation among LMM-fitted animal-mean of all neural parameters computed for RF sessions across cell types (INs, speed cells, and spatial cells), animal age in months, and aging DEG expression (total, up-regulated with age [Up], and down-regulated with age [Down]). Color bar indicates correlation value. **(F)** Bar chart of number of dataset genes and age DEGs (denoted by the label “x Aging”) with expression correlated with spatial coherence and stability (Linear regression, p ≤ 0.05). Correlations with multiple spatial coding variables were calculated iteratively. Bar color indicates the correlation value’s sign. Number denotes total across positive and negative correlates. **(G)** Example genes that are positively (top, *Gene X*, across age groups: r = 0.79, p = 0.0007; among aged mice: r = 0.82, p = 0.0238) and negatively (bottom, *Gene Y*, across age groups: r = -0.60, p = 0.0204, among aged mice: r = - 0.88, p = 0.0059) correlated with LMM-fitted animal mean within map stability. Dots are colored by age group (young [blue] vs. aged [orange]) as in Figure 2G, 2I, and 2K. **(H)** Gene ontology enrichment analysis of the genes negatively correlated with LMM-fitted animal mean within map stability (Fisher’s Exact test followed by FDR calculation). **(I)** UMAP plot of neurons from snRNA-seq of the MEC. Neuronal subtypes are identified by dot color (see Figure S7E), and excitatory neurons (lime green) and interneurons (light blue) are grouped into clouds, colored as in Figure S6A. **(J)** UMAP plot with the relative expression of the positively correlated stability gene core module (n = 130 genes). Color bar indicates expression relative to UMAP background. **(K)** Violin plot of the relative expression of the positively correlated stability gene core module between INs and excitatory (EX) neurons (0.0073 ± 0.0003 vs. -0.0336 ± 0.0006, EX vs. IN, Wilcoxon rank sum test, p < 0.0001). **(L)** As in (J), for the negatively correlated stability gene core module (n = 541 genes). **(M)** As in (L), for the negatively correlated stability gene core module (-0.0244 ± 0.0002 vs. 0.0274 ± 0.0004, EX vs. IN, Wilcoxon rank sum test, p < 0.0001). See also Figure S7.

To examine connections between transcriptomic and functional changes in the aged MEC, we assessed the correlation of gene expression across this dataset and among DEGs with the LMM-fitted animal mean of each *in vivo* neural parameter measured during RF sessions (see Methods) (Figure 7E). Of these parameters, mean spatial cell firing coherence and within map stability were most strongly related to aging transcriptional changes. As such, we identified sets of 7,008 and 1,773 genes with expression related to spatial firing coherence and within map stability, respectively (Figure 7F). Since most genes (1,256/1,773) and nearly all (59/61) DEGs correlated with *in vivo* spatial stability were also coherence-correlated, we focused the analysis on stability-correlated genes (Figure 7G). Next, we applied gene ontology enrichment analysis to all strongly stability-correlated genes (absolute value r ≥ 0.6) (Figure S7C) and the subsets of positively and negatively stability-correlated genes (“positive” vs. “negative correlation” modules). While we found no pathways significantly enriched among positive correlation module genes, pathways related to cytoskeletal organization, neurotransmission, and synaptic signaling were highly enriched in the negative correlation gene module (Figure 7H). Taken together, these results are most consistent with the possibility that cytoskeletal and synaptic gene pathways are up-regulated in response to aged MEC spatial coding dysfunction.

Finally, to determine if stability-associated transcriptomic changes affect particular MEC layers or neuronal subtypes, we employed a snRNA-seq approach in MECs from a pair of unilaterally recorded young and aged mice. We performed UMAP dimensionality reduction on snRNA-seq expression data and annotated the resulting clusters by superimposing the expression of known MEC layer and cell type marker genes (see Methods) (Figures 7I and S7D-E). While DEG expression was relatively uniform across clusters (Figure S7F-I), we observed a striking enrichment of positive correlation module gene expression in excitatory neuron clusters (Figures 7J-K), particularly in Layer 5/6 neurons (Figure S7J). Conversely, expression of negative correlation module genes was specific to INs (Figures 7L-M) and enriched among parvalbumin-expressing (PV+) INs (Figure S7J). These findings converge to suggest that transcriptomic and spatial coding changes co-occur in aged MEC excitatory cells, including grid and speed cells, in a manner coordinated with distinct gene expression changes in MEC INs. Further supporting this, MEC excitatory, but not IN, marker gene expression decreased with age (Figure S7K), suggesting that aged MEC excitatory cells exhibit a loss of function at both the transcriptomic and spatial coding levels. Consistent with decreased spatial firing stability in aged mice (see Figure 5G), however, expression of positive and negative correlation gene modules in MEC decreased and increased with age, respectively (Figure S7K). Since aged MEC INs exhibit increased expression of these stability-correlated synaptic signaling and neurotransmission genes, these findings are ultimately consistent with a protective response to counteract excitatory cell dysfunction related to spatial coding and spatial memory deficits.

## Discussion

Here, we combined VR behavior tasks with *in vivo* electrophysiology to demonstrate that aging is associated with flexible spatial memory impairment and with spatial coding dysfunction at the single cell and network levels in the MEC. Among individual spatially tuned neurons, aging impaired the formation of stable, context-specific firing patterns in the SM task and the maintenance of stable firing patterns in the RF task. Across recording sessions, flexible spatial memory performance correlated with the stability and context-specificity of grid cell firing patterns. At the population level, aged grid networks exhibited impaired flexibility in the SM environment, where remapping occurred more frequently but with less alignment to context switches. Together, these results suggest that disrupted MEC spatial coding may contribute to impaired spatial cognitive flexibility in aging. Starting in middle age, we also found changes in the power of theta and gamma; increased gain and instability of velocity coding across speed-tuned cell types; and compressed grid scale gradients along the dorsal-ventral axis in MEC. These differences constitute potential circuit-level contributors to spatial cell and network instability. Finally, by sequencing the RNA of MEC neurons from these mice, we identified 61 putative transcriptomic mediators of MEC spatial coding dysfunction, which are differentially expressed with age, correlated with spatial cell instability, and enriched among INs.

To explore MEC substrates of aged spatial cognitive decline, we designed the Split Maze (SM) to elicit aging spatial navigation deficits. Rodent spatial memory is canonically measured with 2D mazes, such as the Morris Water Maze,^75^ Barnes Maze,^7^ and spontaneous or forced alternation T-Maze paradigms.^76^ In the SM task, we adapted the latter maze to a linear VR environment, permitting precise control of visual environmental features during the recording of MEC neural activity. In particular, we associated reward locations with VR contexts based on evidence that MEC participates in the recognition and relay of context identity information to the HPC.^77–79^ Rapidly alternating these contexts elicited heterogeneous spatial memory performance among aged mice (see Figure 1J), especially across sexes (see Figure S1G), consistent with previous work. In humans, healthy cognitive aging occurs at highly variable rates,^80–82^ likely driven by a complex interaction among genetic,^83^ epigenetic,^84^ environmental, and other health and lifestyle factors.^85^ Other groups have also reported increased vulnerability of female rodents,^86^ non-human primates,^87^ and humans^88^ to spatial memory decline. Though this makes the statistical discernment of age effects on cognition more challenging, it creates natural experiments to identify factors that influence the rate of cognitive aging. For example, previous studies have stratified rodents into behaviorally age-impaired and -unimpaired groups that exhibited corresponding levels of hippocampal place coding dysfunction.^89–92^ Similarly, studying aged individuals with remarkably preserved cognitive ability, so-called “super agers,” may offer insight into neuroprotective factors.^93^ We observed at least one such aged mouse (A24; see darkest orange line in Figure 1J and Figure S4G) in this study. This behavioral heterogeneity, across and within sexes, among genetically identical aged mice offered us a powerful opportunity to identify functional correlates of spatial cognition in MEC.

Since the HPC and MEC are reciprocally connected regions that cooperate to support spatial cognition, it is highly likely that aged spatial coding dysfunction in these two brain regions interact to impair spatial memory. Supporting this, we observed parallels between spatial coding impairments in aged grid cells in MEC and those previously reported in aged HPC place cells. In MEC, aged grid cells failed to stabilize context-specific firing patterns over sessions, which correlated with their SM alternation impairment (see Figures 2G-K). Aged grid networks also exhibited increased remapping frequency (see Figure 3B). Similarly, in HPC, aged place cells exhibit impaired experience-dependent spatial coding, such as persistent remapping over successive exposures to the same environment over days.^41–43^ Aged place field stability also correlates with spatial learning.^42^ Furthermore, aged rats showed HPC map mis-alignment to external cues correlated with an impairment in goal-directed navigation,^44^ which corresponds to the poor context alignment of aged mouse grid maps we observed (see Figure 3D). Finally, we observed decreased theta power in aged MEC, adding to previous findings that theta rhythm changes in aged rodent HPC^94^ and in aged humans.^95–98^ Theta rhythm across the MEC-HPC network is modulated by medial septal inputs^99–101^ and critically supports navigation and spatial memory.^102,103^ Collectively, these findings raises the following non-mutually exclusive possibilities explaining aged spatial cognitive deficits: (1) unreliable sensory information is conveyed to the MEC-HPC network; (2) the integration of sensory information in either MEC or HPC fails, propagating similar spatial coding errors throughout the network; or (3) similar impairments arise independently in MEC and HPC through convergent molecular and cellular mechanisms. Future work co-recording HPC and MEC in aged mice is needed to further distinguish these possible ways in which grid and place coding deficits interact to impair spatial navigation.

Supporting the first possibility that MEC-HPC sensory inputs degrade, we observed increased speed gain and speed-tuning instability in aged MEC. Since the speed signal is altered across MEC positively speed-modulated speed cells, interneurons, and grid cells (see Figure 6D-E and S6F-G), this raises the possibility that inputs to MEC carrying velocity information are impacted by aging. In addition to impacting the vestibular and visual systems of aged animals,^104^ aging may specifically alter the function of the brainstem circuit recently shown to drive MEC speed cell activity, originating in the pedunculopontine tegmental nucleus (PPN).^105^ In fact, PPN neuronal density decreases with age in rats^106^ and humans.^107^ Examining velocity signals in the aged PPN would help to identify the source of aged velocity coding dysfunction. This is an important, open question because continuous attractor models of grid network activity predict that proportional MEC velocity gain and grid scale changes are required for stable spatial coding and accurate path integration.^72,73^ Increased speed gain and grid scale compression co-occurred in aging (see Figure 6), but it is unclear how precisely coordinated these changes are at behavioral timescales. In addition to speed tuning instability, uncoordinated changes in speed gain and grid scale may contribute to MEC spatial coding instability in aging. Therefore, identifying circuit-level sources of degraded speed information may elucidate how spatial coding instability arises across MEC and HPC.

Providing evidence for the second possibility that MEC–HPC spatial coding deficits interact, we observed a pronounced impairment in fast gamma power (see Figure 4C). Optogenetic perturbation of MEC fast gamma oscillations impairs spatial learning, likely via disturbed synchrony of MEC and dentate gyrus (DG).^65^ Reduced synchrony of MEC and DG oscillations during aging may interact with reduced synaptic innervation of DG and CA3 subregions by entorhinal cortex in aging^108–110^ to impair the transmission of contextual and positional information. Moreover, our findings reveal the positional information output by aged MEC spatial cell networks is significantly degraded (see Figure 5H). These results suggest that MEC spatial information and its transmission to HPC both become less reliable in aging. Additionally, we observed a few aged sessions with high grid network spatial map alignment to context but paradoxically impaired task performance, compared to almost no young or MA sessions of this type (see Figure 3D). Therefore, poor performance in at least some aged mice may be explained by impaired transmission or utilization of MEC context information by downstream brain regions, including HPC. Finally, changes in MEC grid scale alone, as we observed (see Figure 6F-J), might plausibly alter HPC place scale and stability.^62^ Further exploring how aging changes the functional connectivity of MEC and HPC may elucidate how subregion-specific HPC place coding impairments arise.^91,111^

Towards the goal of finding molecular and cellular factors linked with aging in MEC, we identified 458 genes differentially expressed with aging (DEGs) in MEC neuronal nuclei using bulk RNA sequencing. We observed the up-regulation of genes related to oxidative phosphorylation and reactive oxygen species regulation in the aged MEC, consistent with a compensatory response to oxidative stress in normal aging.^112^ Furthermore, we observed increased expression of the AD-associated genes *Apoe* and *App* with age in MEC. This aligns with recent work showing significant overlap between transcriptomic reprogramming in aged and AD non-human primate entorhinal cortex^113^ and between AD-associated human brain gene modules and those in aged wild-type mice.^114^

By examining the relationship between neuronal gene expression and *in vivo* MEC spatial coding stability, we identified positively and negatively stability-correlated MEC gene modules. Consistent with age-mediated loss of MEC excitatory cell function, positively stability-correlated gene expression was enriched among excitatory neurons, especially in Layer 5/6, and decreased with age, along with excitatory cell markers. Aging MEC Layer 5/6 excitatory cell dysfunction might impact long-term memory formation and retrieval by impairing HPC output to neocortical networks, feedback projections from HPC to MEC Layer 2/3, and subfield-specific HPC spatial coding.^115,116^ Conversely, negatively stability-correlated gene module expression was specific to MEC INs, enriched among PV+ INs, and increased with age. Gene ontology analysis revealed that this module related strongly to synaptic transmission and signaling. In young mice, MEC PV+ INs are required for selective spatial coding^70^ and receive inhibitory medial septal inputs that modulate MEC theta rhythm^99–101^ and, in turn, support spatial learning and memory.^102–103^ Taken together, these findings suggest two possible transcriptomic mechanisms by which aging might impair MEC spatial coding: (1) expression of stability-supporting genes decreases in MEC excitatory cells while expression of genes deleterious to stability increases in MEC INs; or (2) altered synaptic and neurotransmission-related gene expression in aged MEC INs occurs as a protective response to excitatory cell dysfunction. As such, future investigations should aim to elucidate the roles of genes up-regulated with age and negatively correlated with spatial coding stability in MEC circuit function. Along with understanding the signaling pathways that modulate the expression of these genes, such studies might uncover tractable molecular targets to improve aged spatial coding and memory dysfunction. Ultimately, by identifying coordinated transcriptomic and spatial coding changes in aged MEC, this work motivates elucidating the role of *Gene Y* and other age-modulated, stability-correlated genes in MEC IN synaptic plasticity, which may be a critical determinant of MEC-HPC spatial coding and spatial memory decline in aging.

## STAR Methods

### Experimental model and subject details

#### Animals

All experimental approaches were approved by the Institutional Animal Care and Use Committee at Stanford University School of Medicine. A total of 18 (8 female, 10 male) young, 10 (5 female, 5 male) middle-aged (MA), and 17 (10 female, 7 male) aged C57Bl/6 mice (Charles River and The Jackson Laboratory) were used for bilateral neural recordings across two behavioral tasks. The young, MA, and aged groups were 3.35 ± 0.31, 12.63 ± 0.09, and 22.44 ± 0.26 months old (ranges: 2.30 - 6.64; 12.30 - 12.99; 21.12 - 24.41) at the time of the last recording. Young and MA mice were acquired from Charles River and the Jackson Laboratory, respectively, at 4 - 6 weeks of age. Aged mice were acquired from the Jackson Laboratory (Stock No. 000664) at 72 - 84 weeks old. All mice were housed in transparent cages (Innovive) with five same-sex littermates. Middle-aged and aged mice were aged in-house to 48 - 53 and 88 - 92 weeks old before surgical headbar implantation. Mice were given an in-cage running wheel 4 weeks prior to this surgery. After headbar implantation, mice were co-housed with 1-3 same-sex implanted littermates unless separation was necessary in response to signs of aggression. After craniotomy surgery and during recording, all mice were single-housed. All mice were kept on a 12-hour light-dark cycle with experiments conducted during the dark phase. An additional young animal underwent surgeries, behavioral training, and sham silicon probe insertion to generate neuronal gene expression data from MEC via single nucleus RNA sequencing (snRNA-seq) data from MEC, reflected in Figures 7I-M and S7D-J.

### Method Details

#### Virtual reality (VR) setup & tasks

The VR recording setup was similar to previously published designs.^23,58,68^ Head-fixed mice ran on a foam cylinder (15.2 cm diameter) fixed to rotate along one axis. Virtual environments were generated using commercial software (Unity 3D) and displayed on three 24-inch monitors around the mouse. A quadrature encoder (Yumo 1024P/R) measured cylinder rotation, which a microcontroller (Arduino Uno) processed into motion signals to advance the VR environment. VR track gain was calibrated such that each track was 400 cm long. Upon reaching 400 cm, mice were teleported seamlessly to the track start, making the track seem circular. Each VR track featured black and white visual cues, including a patterned floor for optic flow and five pairs of landmark towers spaced 80 cm apart, each with unique dimensions and patterns (all with neutral luminance) (see Figures 1B, 1F, S1A, and S2A). Water rewards were delivered via a custom-built lickport, consisting of Tygon tubing attached to a metal spout in front of the mouse. In reward zones, mouse licks (“requests’’) resulted in reward delivery. Licks broke an infrared beam positioned across the lickport, which triggered an audible solenoid valve (Cole Palmer) to pump water via a microcontroller (Arduino Uno). Failure to lick before the reward zone center resulted in an automatic reward delivery in some training and recording conditions. For each VR frame, the Unity engine recorded mouse track position and frame time. Reward zone locations, lick times, and reward delivery times were also recorded.

With this setup, we collected behavioral and neural data in two VR tasks: the Split Maze (SM) and the Random Foraging (RF) tasks. For the first 20 SM trials, mice ran in darkness with no visual cues. On each subsequent trial, mice encountered one of two sets of distinct visual cues (contexts A and B), each associated with one of two possible hidden reward zones (50 cm long, centered at 270 or 370 cm). Contexts A and B were introduced sequentially for ∼60 trials each (“blocks”) and were then pseudo-randomly alternated for another ∼80 trials (“alternation”) (see Figure 1C). For the first 10 trials in each block, rewards were automatically delivered if not requested. The association of reward locations with contexts was counterbalanced within age groups to control for the impact of the locations of hidden rewards. The first 200 cm (“front”) of the track was mostly shared across contexts, featuring a black and white diamond checkerboard floor and three pairs of rectangular prism towers, each with different sizes and patterns (see Figure S1A). In contexts A and B, respectively, a small cue tower was positioned immediately to the left or right of the floor at 40 cm from the start. Large black doors, which opened when the mouse was 10 cm away, obscured the second 200 cm (“back”) of the track and a uniform gray landscape after 400 cm. On context A trials, the back of the track featured a floor with horizontal and vertical lines and two different cylindrical tower pairs. On context B trials, the back featured a dotted floor and two bullet-shaped tower pairs. During the last 20 alternation trials, the gain of the VR track translation to motion signals was decreased to 0.7, requiring the mouse to run 568 cm to advance the VR environment 400 cm.^58^ We aimed for each SM mouse (n = 9 young [4 male, 5 female], 10 MA [5 male, 5 female], and 10 aged [5 male, 5 female]) to complete the task daily for six days. Mice A16 and MA7 showed signs of peri-craniotomy skin infection by the sixth session, so only five recordings were completed. Mouse MA4 exhibited signs of distress, including long periods of freezing, during its final recording. Behavioral data for the last two completed sessions of each of these mice were excluded. Finally, recording from A14 was terminated after the third session due to poor running and licking behavior post-craniotomy. These sessions are included in neural and behavioral datasets, since A14 completed all SM task trials. snRNA-seq MEC gene expression data from this animal are also reflected in Figures 7I-M and S7D-J.

In the RF task sessions, mice completed ∼200 trials of running on a track with unchanging visual cues, including a checkerboard floor and five cylindrical tower pairs.^23^ In addition, 50 cm reward zones appeared at a probability of p = 0.0025 or 0.005 per cm, titrated to mouse performance, within the middle 300 cm of the track and at least 50 cm apart. Reward zones were marked by a diamond checkerboard pattern hovering slightly above the floor. After a successful reward request or complete reward zone traversal, the current zone disappeared, and the next became visible. In a subset of RF sessions (male mice only), 20 trials of dark running and 20 trials of gain manipulation were added to the beginning and end of the task, respectively. We also aimed to collect six sessions from each RF mouse (n = 9 young [6 male, 3 female], 7 aged [2 male, 5 female]). Hardware failures resulted in the collection of only five sessions from Y2, Y3, and Y11 and three sessions from Y16. Finally, recording from Y9 was terminated early due to poor running and licking behavior post-craniotomy. Behavioral data from these three partially completed sessions were excluded.

Before recording, a third task was administered to SM mice to estimate their visual acuity. In this task, we repurposed the RF track but adjusted the reward zone opacity as mice ran 100 trials with a reward probability of p = 0.005 / cm (see schematic in Figure S2C). For the first five trials, reward zones appeared at full opacity (α = 1). For the next five trials, reward zones were invisible (α = 0). For remaining trials, reward zone opacity was pseudorandomized (0 < α < 1). We also changed the floor to a uniform gray color and dropped the reward zone to the floor to prevent mice from recognizing low opacity reward zones by subtle optic flow changes or the presence of a floating obstacle. To estimate visual acuity, we calculated the fraction of rewards requested for binned reward zone opacities (bin size = 0.1) and fit that with the following sigmoid function using scipy (optimize.curve_fit), initialized at center = 0.1 and scale = 0.002:

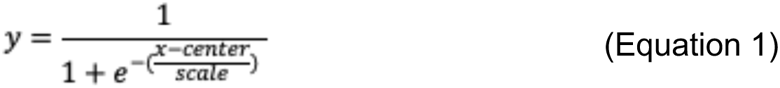

The center of the resulting psychometric curve was each animal’s estimated visual acuity threshold (see Figure S2D).

#### In vivo survival surgeries

A mixture of oxygen and isoflurane was used to induce (4%) and maintain (0.5 - 2%) anesthesia. Perioperative pain control was achieved with a subcutaneous buprenorphine (0.05-0.1 mg/kg) injection post-induction. Following surgery and for three additional days after, mice were monitored and administered Baytril (10 mg/kg) and Rimadyl (5 mg/kg) subcutaneously. In the first surgery, we attached a custom-built titanium headbar, implanted a screw soldered to a gold ground pin, and made fiducial marks on the skull. These marks were made at ±3.3 mm relative to midline and 3.7 mm posterior to bregma to guide Neuropixels probe insertion. The ground pin was implanted ∼2.4 mm anterior and ∼1.2 mm lateral of bregma. Metabond (Parkell S380) was used to cover any exposed skull surface and secure the ground pin and headbar. After behavioral training, we made bilateral craniotomies (∼200 μm in diameter) at 3.7 - 3.95 mm posterior and 3.2 - 3.4 mm lateral to bregma in a second surgery. These were posterior and centered to fiducial marks and exposed the transverse sinus. Bilateral durotomies were made central to the craniotomy sites using a bent 30-gauge needle. A plastic well was also implanted around each craniotomy and fixed with Metabond. Craniotomy sites were washed with sterile saline and sealed with a silicone elastomer (Kwik-Sil, World Precision Instruments) to conclude surgery and each recording day. After both surgical procedures, mice were administered 1mL of saline and placed on warming blankets to recover.

#### Training and handling

After headbar implantation, mice were handled for at least 5 minutes twice each day. Three days after implantation, they were restricted to 0.8 - 1.5 mL water per day with *ad libitum* access to food. Mice were monitored and weighed daily to maintain them at 80-85% of *ad libitum* postoperative weight. Water volume was increased within this range if signs of dehydration, such as skin tenting, were observed. Any water consumed during training or recording in VR was supplemented afterwards in the home cage to reach the total allocated amount.

After initiating water deprivation, mice were trained in phases (A - C) to run in VR and request rewards according to implicit rules. In Phase A, mice acclimated to head-fixation and moving in VR. Initially, the experimenter helped advance the cylinder forward and prevent backward motion in two daily sessions lasting 10 minutes each. When mice initiated forward movement independently, training time increased to 20 minutes, and the VR monitor was turned on, displaying the RF track. The next day, we introduced the lickport and dispensed water rewards (∼1.5 - 2 μL per reward) automatically at each randomly appearing reward zone (p = 0.10 / cm). With two daily 30-minute training sessions, mice learned to associate solenoid clicks with reward delivery and, eventually, to request rewards by licking within cued reward zones. Gradually, we increased goal trial number from 10 to 25 to 50 and 100, while decreasing reward probability from p = 0.10 to 0.05 to 0.01 to 0.005. When mice requested >80% of rewards before automatic delivery, we switched to operant dispensation.

When mice operantly requested >80% of rewards and ran 100 trials in 30 minutes, training protocols diverged for the SM and RF tasks. SM task mice were next administered the visual acuity assessment task described above. Then, adapting the tower and floor cues from the RF track, SM mice were trained to lick in a single hidden reward zone per session (Phase B) and, then, to abruptly switch between rewarded locations during the session (Phase C). In Phase B, mice ran 110 trials and received automatic rewards in a hidden zone (switching between centers at 220 or 320 cm each session) for periods of 60 and then 40, 20, and 10 trials. When mice acquired the hidden reward location within 10 trials, Phase C began. The trial number doubled to 220, and the reward location switched at 110 trials. Only 10 automatic rewards were delivered at the session start and after the switch to indicate reward location. SM mice were considered ready for recording when they completed >220 trials / hour and consumed >80% of rewards at both locations. After Phase A, RF task mice continued daily RF track training, lasting a maximum of one hour. The goal trial number increased to 150, 200, and then 250 with reward probability (p) = 0.0025. RF mice were considered ready for recording when they completed >250 trials/hour and operantly requested >80% of rewards. Therefore, the SM track cues were novel to the mice at the start of recording, whereas the RF track cues were highly familiar.

#### In vivo electrophysiological data collection

All neural recordings occurred at least 16 hours after the craniotomy surgery upon animal recovery. To record neural activity, mice were head-fixed on the VR rig and a single craniotomy site was exposed and rinsed with saline. Immediately prior to recording, Phase 3B1 Neuropixels 1.0 silicon probes^58^ with ground and reference shorted together were dipped in one of three fixable, lipophilic dyes (DiD, DiI, DiO, ThermoFisher V22889) 10-20 times at 10 second intervals. Using a motorized micromanipulator (MP-225, Sutter), the probe was positioned over the craniotomy site at 10.6° from vertical and targeted to ∼50-300 μm anterior of the transverse sinus. The probe was located behind the mouse to minimize visual disturbances to the VR environment (Figure 1A). To sample new cell populations on each day, three recordings were made per hemisphere, targeting at least 50 μm medial or lateral of previous recording sites on consecutive days. The 384 active recording channels on the probe were set to occupy the 4 mm closest to the tip. The reference electrode was connected to the ground pin, and then the probe was advanced slowly (1 - 5 μm/s) into the brain until mechanical resistance was met or activity near the probe tip quieted (∼3,000 - 4,000 μm from surface). The probe was retracted 50-100 μm and allowed to settle for at least 15 minutes. Finally, the craniotomy site was covered with sterile saline and silicone oil (Sigma-Aldrich 378429). Final probe depth and insertion position in medial-lateral and anterior-posterior dimensions were recorded daily.

Signals were sampled at 30 kHz with gain = 500 (2.34 µV/bit at 10 bit resolution) in the action potential band (high-pass 300 Hz to 10 kHz filter), digitized with a CMOS amplifier and multiplexer built into the probe, then written to disk with SpikeGLX software (https://billkarsh.github.io/SpikeGLX/). Local field potentials were similarly sampled at 2.5 kHz with gain = 250 (4.69 µV/bit at 10-bit resolution) and 0–300 Hz low-pass filter. Each Unity VR frame emitted a TTL pulse, which was then relayed by a microcontroller (Arduino Uno) to an auxiliary National Instruments data acquisition card (NI PXIe-6341 with NI BNC-2110). VR frame times were thus also recorded by SpikeGLX, permitting synchronization of VR data with electrophysiological traces.

#### Offline spike sorting

The processing of electrophysiological recordings was performed by adapting Jennifer Colonell’s fork of the Allen Institute for Brain Science ecephys library (open source from Github: https://github.com/jenniferColonell/ecephys_spike_sorting). Briefly, this pipeline employed CatGT (https://billkarsh.github.io/SpikeGLX/help/catgt_tshift/) to perform global common averaging referencing (-glbcar) and filter transient artifacts that appear on >25% of channels with peak amplitudes of at least 0.40 mV (-gfix=0.4,0.10,0.02). Spike sorting and drift correction were then performed using the open source algorithm Kilosort 2.5.^117^ Kilosort 2.5 improves on Kilosort 2, which identifies clusters in neural data but only tracks neural drift (https://github.com/MouseLand/Kilosort2).^118^ Next, ecephys modules extracted waveforms from the Kilosort output and calculated their properties (e.g. signal:noise ratio, duration, halfwidth, and peak:trough ratio) and the quality of corresponding spike clusters (e.g. autocorrelation contamination rate and interspike interval (ISI) violation rate and number). Clusters were curated in Phy (https://github.com/cortex-lab/phy). All clusters with fewer than 100 spikes were excluded. Clusters with signal:noise ratio < 1.0, firing rate (FR) < 0.10 Hz, repolarization slope < 0, depth > 3200 μM from the probe tip, or halfwidth > 0.30 ms were excluded as noise using an open-source filtering module created by Emily A. Aery Jones (https://github.com/emilyasterjones/ecephys_spike_sorting/tree/master/ecephys_spike_sorting/ <u>modules/prePhy_filters</u>). All remaining clusters were manually examined and labeled as “good” (i.e. likely corresponding to a well-isolated neural unit) or “MUA” (likely to represent multi-unit activity) based jointly on contamination rate and ISI violation rate and number. Only “good” units, or cells, with greater than 350 spikes were included for analysis in this paper. Sessions with fewer than 10 cells meeting these criteria were excluded from our neural datasets.

#### Behavioral data preprocessing and synchronization

Clustered, curated neural spiking data were then synchronized to VR behavior data using the TTL pulse times recorded by SpikeGLX and saved by Unity with custom MATLAB code. Appropriate syncing was confirmed by noting a high correlation between VR frame time differences and corresponding TTL time differences in SpikeGLX. As in previous work,^23,57^ we used interpolation to resample behavioral data at a constant 50 Hz framerate, since VR frame times are not constant. As the tracks were 400 cm long and effectively circular given teleports, recorded positions less than 0 or greater than 400 cm were converted to the appropriate position on the previous or subsequent trial. Trial transitions were identified as time points where the difference in position across time bins was less than -100 cm (i.e. from ∼ 400 cm to ∼0 cm). As such, each VR frame was assigned a trial number. Running speed was computed as the difference in VR position between consecutive frames, divided by the framerate. After removing and interpolating over frames with speeds < -5 cm/s or >150 cm/s, speed was smoothed with a Gaussian filter (standard deviation = 0.2 time bins). For all analyses, except lick and reward zone analyses (Figure S1E, I, & J and Figure S2C-G), stationary time bins (speed < 2 cm/s) were excluded. The stereotypy of reward-triggered licking and slowing for each session (Figures S1E, S2F) was examined by averaging position-binned lick counts and speed traces (bin size = 2.5cm) across rewarded trials from 25 cm before to the end of the reward zone (75 cm total). We next found the location of absolute maxima in the mean lick and inverted mean speed traces and used the scipy.signal package’s peak_prominences function to find the prominence of licking and slowing there, termed slowing and licking magnitude.

#### Tissue collection & storage

Immediately after probe explant after the final recording session, mice were sacrificed with an overdose of Euthasol (Virbac) and transcardially perfused with ice-cold phosphate-buffered saline (PBS). Brains were extracted and split into hemispheres. One hemisphere was selected at random to be flash frozen in a sterile RNase free 1 mL Eppendorf tube on dry ice and stored at - 80°C for subsequent sub-dissection of the entorhinal cortex, nuclear RNA isolation, and RNA sequencing. The total time from recording end to hemisphere freezing was under 30 minutes for all mice. In preparation for histology, the remaining hemisphere was externally fixed in 4% paraformaldehyde in 0.1M PB pH 7.4 for 48 hours followed by cryoprotection in 30% sucrose. Fixed hemispheres permit validation of observed gene expression changes via RNA sequencing. To account for the loss of histologic confirmation of probe targeting, we required that recording sessions included in these analyses had greater than 50 probe channels with coordinated theta-rhythmic neural activity during running, which constitutes an electrophysiological signature of MEC.^119^

#### RNA isolation and bulk RNA-seq

Neuronal nuclei were isolated based on the protocol previously described with minor modifications.^120^ Briefly, MECs, dissected from flash-frozen hemispheres at ± 3.1 - 3.5mm lateral and 4.75 - 5.25mm posterior to bregma below 3.5mm from dorsal brain surface, were dounce-homogenized (Wheaton, Cat. 357538) in 500 μL of EZ lysis buffer (Sigma Aldrich, NUC101) with 1 U/μL of RNase Inhibitor (Sigma Protector, 3335399001). Given this dissection approach, some nuclei from the lateral entorhinal cortex and parasubiculum may be included in this dataset. Samples were homogenized with 20 strokes each of the loose and tight pestles, and 500μL of lysis buffer was added to the samples. Samples were incubated on ice for 7 minutes before filtering through a 40 mm filter and centrifuging at 500 RCF for 5 min at 4°C. Conjugated mouse monoclonal anti-NeuN-AlexaFluor488 antibody (Millipore, Cat. MAB377X, RRID: AB_2149209) in a staining buffer (PBS with 1% BSA and 1U/μL RNase Inhibitor) was added to the tube at a final dilution of 1:250. Samples were incubated on a tube rotator for 30 min at 4°C and then spun for 5 min at 500g at 4°C. Samples were resuspended in 400μL of staining buffer with Hoechst 33342 at a final concentration of 0.01 mg/mL. Samples were then filtered through a 35mm Fluorescence-activated cell sorting (FACS) tube filter and sorted.

Nuclei were sorted by FACS on a BD FACSAria Fusion with a 100 mm nozzle and with a flow rate of 1–2.5. Nuclei were first gated by forward and side scatter, then gated for doublets with height and width. Nuclei that were both Hoechst+ and NeuN+ were sorted into the staining buffer with 2x concentration of RNase Inhibitor (see Figure S7A). Nuclei were spun down at 500 RCF for 5 min at 4°C and resuspended in 250 μL Tri Reagent (Sigma Aldrich, T9424) for RNA expression analysis.

#### Single nucleus RNA-seq

Nuclei for snRNA-seq were isolated similarly to bulk samples. MECs from a young and aged mouse were sub-dissected and dounce homogenized as above. snRNA-seq samples were also stained with Hoescht 33342 and filtered as above, followed by FACS-sorting with the same parameters. Nuclei that were Hoescht+ were sorted into staining buffer with 2x concentration of RNase Inhibitor, spun down at 500g for 5 min at 4°C, and resuspended in the staining buffer. Nuclei were counted, and 25,000 from each sample were loaded onto a 10x Chromium Controller. Libraries were generated following 10x Genomics protocols and sequenced to a depth of 30,000 reads per nuclei on an Illumina 10B X. Fastqs were converted to feature barcode matrices using CellRanger 7.0, and downstream filtering and analysis was performed using Seurat v5.^121^

### Quantification & Statistical Analysis

#### Statistical tools & reproducibility

Neural and behavioral data analysis was conducted in Python (3.8.16) using the NumPy (https://numpy.org/) and SciPy (https://scipy.org/) libraries to compute statistics and perform linear regressions. Linear mixed effects and logistic regression modeling were performed using the statsmodels package (https://www.statsmodels.org/). Non-parametric tests were used to avoid assumptions about the normality of data distributions. Specifically, we applied Wilcoxon signed-rank tests to assess significance for paired data; Wilcoxon rank sum tests for unpaired data; and Kruskal-Wallis H-tests for comparisons of 3 groups. For multiple comparisons within 3 groups, we performed Holm-corrected Conover’s test using rank sums as a post-hoc test with the scikit_posthocs package (https://scikit-posthocs.readthedocs.io/). To assess the significance of differential gene expression, we performed a Wald test followed by Benjamni and Hochberg corrections of multiple comparisons). Gene ontology biological process enrichment was confirmed by Fisher’s Exact test and false discovery rate calculation. Significant gene set enrichment was assessed by a permutation test followed by family-wise error rate correction. followed Unless otherwise noted, all tests were two-sided; averaged data are presented as mean ± standard error of the mean (SEM); and correlation values are Pearson correlation coefficients. Data collection and analysis were not performed blind to conditions of the experiments. Sample sizes were consistent with previous similar studies and were not predetermined. Levels of significance across all figures are indicated as follows: * p < 0.05, ** p < 0.01, *** p < 0.001, **** p < 0.0001. Results that approached significance (0.05 < p < 0.10) are indicated by a hashmark (#).

#### Linear mixed effects models

To quantify behavioral performance and neural activity changes across sessions, we used linear mixed effects models that account for variance across animals. All models treated animal identity as a random effect, converged using restricted maximum likelihood estimation unless otherwise noted, and were implemented using the mixedlm method of the statsmodels.api package. We allowed only random intercepts for mice after confirming that including random slopes did not affect results, unless otherwise noted here.

For behavioral models (Figures 1E, 1H, S1E-G, S1I-J, and S2F), fixed effects included age group as a categorical variable; session number as a continuous variable; the interaction of session and age group; sex as a categorical variable; and the interaction of sex and age group. In the models of SM block and alternation performance, we also included reward location order (context A reward centered at 270 or 370 cm) and the interaction of reward order and age group as categorical fixed effects. To model performance across SM task epochs (Block A [A], Block B [B], Alternation A Trials [A’], Alternation B [B’] Trials), we added the following other categorical fixed effects: reward order, VR context (A or B), the interaction of context and age group, task structure (block vs. alternation), and the interaction of task structure and age group. In the four models in Figures 1I and S1F-G, random slopes were also allowed for mice, since this improved model fit. When the behavioral model dependent variable was the fraction of reward requested, we confirmed that applying a logit transform of these fractions did not qualitatively change results or improve model fit. For interpretability, we chose to report untransformed fractions.

For models of neural activity features (Figures 2F, 2H, 2J, S3I, S3K, S3M, and 6J), we also accounted for variance across cells using a nested random effects design. Fixed effects in these models included age group, sex, recording cohort, and the interaction of session and age group. Cohort (A - D) refers to groups of mice trained proximally in time and recorded sequentially as follows: all female RF mice (A), all male RF mice (B), all female young and aged SM mice and MA mice (C), and all male young and aged SM mice (D). To correlate gene expression with *in vivo* cellular features, like spatial firing coherence and within map stability (Figures 7E-G), we fit LMMs as described here to each cellular parameter computed across RF sessions (see Figure 7E) and then calculated the animal mean fitted measure. This approach accounted for variability in these parameters across cells and animals and for differences in session number and spatial cell density across sessions (see Figure 5D). The models of spatial firing stability by VR context and mean similarity across context-matched VR epochs (Figures 2F 2H, S3I, S3K) include mean non-consummatory licks per trial in the context of interest as a continuous fixed effect. The model of grid scale included unit recording depth as a continuous fixed effect. When the dependent variable was a similarity ratio (Figures 2J, S3M), we also confirmed that a logit transform of ratios did not qualitatively change results or improve model fit.

#### Spatial firing rate & autocorrelation, shuffle procedure, and distance tuning

We generated spatial firing rate vectors empirically for recorded cells by binning track position into 2 cm bins and dividing spike counts by occupancy for each bin. We performed this for every trial to generate by-trial spatial tuning curves or across trials to generate an estimate of average firing rate as a function of position. When indicated in subsequent methods, we smoothed this vector with a Gaussian filter (standard deviation = 4 cm).

To assess distance tuning, reflected in periodic activity during dark running,^57–59^ we computed the autocorrelation of the smoothed by-trial spatial firing rate vector, linearized across dark trials, using lags up to 800 cm.^57^ Next, we used scipy.signal package’s find_peaks function to detect the prominence and lag of the maximum peak in the autocorrelation with a width > 10 cm, height > 0.1, and prominence > 0.15. Additionally, we computed a shuffled spatial autocorrelation after shifting spike times relative to elapsed distance by random offsets (n = 100 shuffles, offset drawn from a uniform random distribution with the interval [20 seconds, max_t] where max_t was the recording duration). For each shuffle, we computed the height of the autocorrelation at the same lag as the original maximal peak. A cell was determined to be distance-tuned if it had at least one qualifying peak in its dark autocorrelation and the maximal peak prominence of the autocorrelation exceeded the 99th percentile of the shuffle distribution heights at that same lag.

The same spike time shuffle procedure was repurposed for the classification of speed-tuned and spatial cell types across both tasks and as a control for the comparison of cell and network-wide spatial firing quality across age groups in the RF task (Figures 5E-H, S5E).

#### Calculation of spatial coherence, sparsity, and information

Spatial coherence was calculated as follows to assess the strength of spatial firing by measuring the local smoothness of the spatial firing rate vector.^122^ First, a non-smoothed, trial-averaged spatial firing rate vector was calculated for each cell as above. For each 2 cm bin, the mean firing rate of the nearest eight bins was calculated. Then, coherence was computed as the correlation of firing rate with the mean nearest bin firing rate across bins, such that coherence ∼ 1 implies perfectly spatial firing. Spatial sparsity was calculated as follows to assess the coverage of the VR track by each cell’s spatial firing,^123^ where P_i_ is the probability of occupancy of bin i, R_i_ is the mean firing rate in bin i, and R is the overall mean firing rate:

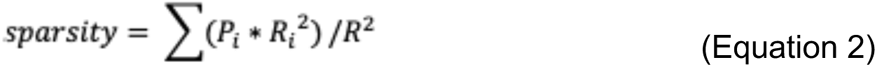

Finally, spatial information was calculated in bits per second over position bins according to the previous work,^124^ as follows, with variables defined as above:

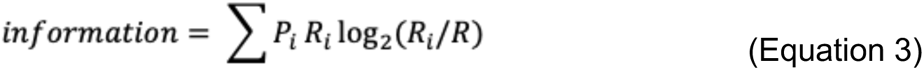

#### Speed tuning analyses

Speed tuning quality was determined by three scores: speed score, speed stability score, and trial stability speed score. Speed score was defined as the correlation between speed and instantaneous firing rate,^20^ calculated by smoothing the vector of spike counts at each time point with a Gaussian kernel (standard deviation = 40 cm). Speed slope and intercept were calculated using least-squares linear regression of instantaneous firing rate and speed to estimate speed tuning strength. To account for the correlation of running speed and position in VR, we calculated a speed stability score that takes the spike-weighted average of speed scores across track sections (5 x 80 cm):^58^

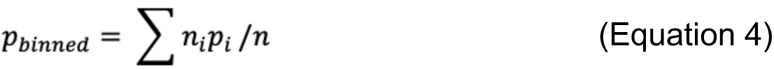

where n is the total number of spikes, i is the position bin index, and n_i_ and p_i_ are the number of spikes and speed score, respectively, calculated in the ith bin. Similarly, we calculated a trial speed stability score by taking the spike-weighted average of speed scores calculated on individual trials, such that i was trial instead of position bin. Trial stability or speed stability scores below the speed score for a given cell indicate instability in speed tuning over trials or position bins, respectively.

#### Identification of functional cell types

Putative excitatory cells were separated from putative fast-spiking INs using thresholds on waveform duration and mean firing rate. Putative INs had a waveform duration <0.35 ms or a mean firing rate (FR) >40 Hz. For each session, we confirmed that the union of these thresholds captured the clustering of co-recorded cells in peak:trough ratio and duration space and separated cells into low waveform halfwidth, high FR vs. high waveform halfwidth, low FR groups (Figure S6A).^125^ Among putative excitatory cells, we classified positively (+) and negatively (-) modulated speed cells.^20^ A cell was speed-tuned if the absolute value of its speed score and speed stability score each exceeded the 99th percentile of absolute value of speed and speed stability scores calculated on spike-shuffled data (n = 100 shuffles; see *Spatial firing rate & autocorrelation, shuffle procedure, and distance tuning*). Speed cells were split into + or - groups based on speed score sign. Additionally, we classified spatial cells among putative excitatory cells using spatial cell coherence and sparsity scores. In the RF task, cells were defined as significantly spatial if their spatial coherence and sparsity scores both exceeded the 99th percentiles of shuffle coherence and sparsity scores for that cell (n = 100 shuffles). In the SM task, cells were defined as spatial if their Block A spatial coherence and sparsity scores both exceeded the 99th percentile of shuffle distributions of the corresponding scores. Only Block A trials were used to minimize the effect of context-dependent changes in spatial firing activity on these scores. Putative grid cells were identified as excitatory cells that exhibited significant distance tuning in the dark (see *Spatial firing rate & autocorrelation, shuffle procedure, and distance tuning*).^57,59^ In SM sessions, putative non-grid spatial (NGS) cells were defined as the spatial cells that did not meet distance-tuning criteria (Figure S3A-F). Putative grid cells with low dark (≤0.05 Hz) or overall (<0.3 Hz) mean FR were excluded. For simplicity, we refer to putative grid and putative NGS cells as grid and NGS cells elsewhere. In some SM sessions, we observed a minority of classified grid and NGS cells with implausibly narrow (<4 cm) firing fields exclusively at reward delivery locations, consistent with unsuccessfully filtered lickport artifacts. To exclude these units, we applied a threshold on noise in trial-averaged spatial FR (mean Block A FR SEM / mean Block A FR > 0.45). Finally, we also identified the subsets of non-spatial + or - speed cells, termed + or - speed only cells, and + speed-tuned INs and conjunctive speed-tuned grid cells, termed + IN speed cells and + grid speed cells, respectively (Figures S6E-G).

#### Cross-trial correlation matrices and spatial stability and similarity

To compute the spatial stability in both tasks and spatial similarity across VR contexts in the SM task, we first computed a smoothed (Gaussian filter, standard deviation = 5 cm), normalized by-trial spatial firing rate. As previously described,^23^ we re-scaled each neuron’s firing rate separately to range between zero and one as such:

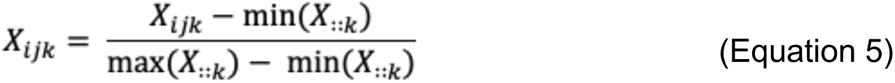

Here, X_ijk_ denotes an I x J x K array, or tensor of normalized spatial firing rates, where i, j, and k respectively represent trials, position bins, and cells. For SM sessions, we sorted alternation trials by VR context. As such max() and min() operators were applied on a neuron-by-neuron basis. This rescaling step prevented neurons with high firing rates from washing out low firing rate neurons in population analyses.

To generate a cell’s cross-trial correlation matrix, estimating the similarity of spatial firing across trial pairs (as in Figures 2E and S3H), we subtracted the mean FR from each trial in the matrix X_ij_ and computed the normalized cross-correlation of each pair of trials at a lag of 0 cm. Each cell’s spatial stability score for a given set of trials was calculated as moving average correlation across all pairs of neighboring trials (5 nearest trials, see Figure 2F, S3I, 5G). In Figures 5G and 5J, within map stability refers to stability calculated within each k-means labeled map and averaged across maps for each spatial cell (see *Population analysis and clustering algorithm with optimized k-values*). This approach controls for reduced local stability caused by any spontaneous remapping events, which occur equally rarely across age groups in the RF task (see Figure S4I). To calculate the similarity of each cell’s spatial firing across blocks of trials, we computed the mean pairwise cross-correlation of spatial firing vectors among those trials. In particular, we compared spatial firing similarity across context-matched and -mismatched SM task epochs (matched: A x A’, B x B’; mismatched: A x B’, B vs. A’) for each cell. To calculate context-matched similarity across task epochs, we required that a given cell’s mean pairwise similarity across trials in either baseline epoch (A or B) exceed 0.1. Finally, to estimate the relative similarity of context-matched vs. -mismatched task epochs, we computed a similarity ratio for SM grid and NGS cells as follows (see Figures 2J and S3M):

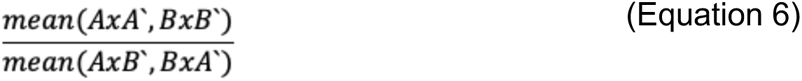

Here, AxA’, BxB’, AxB’, and BxA’ denote the pairwise similarity of all trials across those task epochs. This ratio equaled 1 if context-matched trial spatial firing was equally as mutually similar as context-mismatched trial spatial firing. A ratio > 1 implies context-dependent orthogonality of spatial firing.

To generate network-wide trial-by-trial similarity matrices (Figures 3A, S4A-C, F, G, and S5I), we computed the correlation for vectors vec(X_i::_) and vec(X_ì::_) for all pairs of trials (i, ì) for all spatial cells in an RF session or all grid cells in an SM session. RF sessions with ≤ 10 spatial cells and SM sessions with < 10 grid cells were excluded from population-level analyses, such spatial map clustering and position decoding.

#### Estimation of gain change responses and remapping coordination

Previous work has shown that dissociating visual flow and locomotion signals with reduced VR gain might identify putative grid cells, which are more sensitive than non-grid spatial (NGS) cells to gain reductions.^57,58^ We compared the relative magnitude of SM grid vs. NGS cell responses to VR gain manipulation during alternation to validate the classification of these two functional cell types (Figure S3F-G). To do this, we selected a set of A’ trials before the gain change equal in the number to A’ trials after the gain change. We then generated smoothed by-trial spatial firing rate vectors and cross-trial correlation matrices for each grid and NGS cell for those trial sets. Then, we computed the difference between the mean pairwise similarity within A’ trials before the gain change and the mean pairwise similarity of A’ trials before and after the gain change (within baseline - across gain change). The larger this difference, the greater the cell’s gain change response. We replicated these steps to find the magnitude of B’ trial gain change responses and confirmed that the grid cell responses exceeded NGS cell responses for both contexts in each age group (Figure S3F-G).

To assess whether network-wide matrices would similarly reflect the composite of co-recorded cell activity in each age group, we estimated and compared remapping coordination across groups according to established methods.^57^ In brief, remapping coordination was calculated as the correlation between the cross-trial correlation matrix of each unit and the network-wide similarity matrix, assembled using the remaining co-recorded units. No age group differences in remapping coordination were observed in either task (Figure S5C-D). Similarly, we computed the correlation between network-wide similarity matrices generated using spatial firing tensors from the front and back halves of the SM track (Figure S5I). We observed lower similarity between front and back matrices in MA and aged vs. young sessions (Figure S5J), indicating a stronger modulation of network activity by back of track visual cues. To control for this age effect on the timing of context recognition, spatial firing tensors used for population-level analyses in the SM task (Figures 3, S4, and S5) were assembled using back of track spatial firing activity only.

#### Population analysis and clustering algorithm with optimized k-values

To identify discrete spatial maps, consisting of clusters of trials with similar network-wide spatial firing patterns, in a data-driven manner, we adapted the factorized k-means clustering algorithm described in detail in Low et al. (2021) using Alex Williams’ open-source lvl package (https://github.com/ahwillia/lvl/tree/master/lvl).^23^ Briefly, the low-rank factorization model of the network’s normalized firing rate tensor (X_ijk_) is as follows:

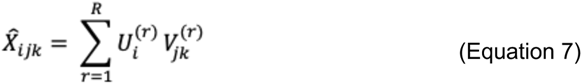

In general, R denotes the number of model components, or the model rank. As described in Low et al. (2021),^23^ k-means clustering arises as a special case of this model, in which R represents the number of clusters, traditionally termed k, or the number of spatial maps. In this case, U_i_^(r)^ gives the elements of an R x I matrix, where each row gives the cluster assignment, or map label, for every trial I, coded in R-dimensional space (“one-hot vectors”). V_jk_^(r)^ comprises the elements of an R x J x K array. For a given map label, r, the J x K slice of this matrix specifies a cluster centroid. This slice is interpretable as a spatial map in which the columns are J-dimensional vectors containing each neuron’s spatial tuning curve. Identifying synchronous, network-wide transitions between spatial maps permitted the comparison of remapping frequency across age groups in each task (Figures 3B and S4H). Moreover, this enabled us to compare the alignment between spatial map and SM context transitions in each age group and over sessions.

To capture the heterogeneity in trial-by-trial structure observed in network-wide similarity matrices, especially in response to the SM context switches, we first optimized the selection of R for each session individually with the range r = 1 - 4. For interpretability, we substituted the signifier k for R as the k-means model rank in all figures and figure legends. To select k, we used a silhouette score maximization approach deploying scikit-learn’s silhouette_score function to measure average cluster quality.^126^ For each k > 1, we fit a factorized k-means model with k components and 100 restarts and calculated the average silhouette score across maps on each of 10 repetitions. Restarts accounted for model variability and helped to select the model best fit to the data within each repetition. The k that maximized the mean silhouette score across repetitions was selected (see Figure S4A).

To set the maximum possible k, we swept the range k ≤ 8 for SM sessions using this optimization procedure. Fewer than 10% of eligible SM sessions (n = 13 / 134) had an optimal k > 4. We compared the train and test cross-validation performance (R^2^) of the k-means model at all possible k to assess if overfitting occurred at higher k. Indeed, we observed that an optimal k > 4 increased test R^2^ minimally compared to the next best smaller k, while increasing train R^2^ significantly (Figure S4B). In each session where k > 4, we also observed at least one low quality cluster with no silhouette scores exceeding the average across clusters. This indicated to us that k > 4 was rarely, if ever, appropriate to capture SM network-wide spatial firing patterns.

To exclude sessions that were not well-approximated by k > 1 models, termed “one-map” sessions, we compared the cross-validated performance (uncentered test R^2^ averaged over 10 replication sets) of k-means models fit on real vs. shuffled datasets. Specifically, we employed a randomized cross-validation procedure in which 10% of the data were censored in a speckled-holdout pattern.^127^ As in previous work,^23^ firing rate tensors (X_ijk_) were shuffled using a random rotation across trials. This preserved overall data norm and correlations between neurons and position bins but disrupted the sparsity pattern on U_i_^(r)^ imposed by k-means modeling. One-map sessions were those where the average model performance at the selected k on real data did not exceed that on shuffled data using a one-sided Wilcoxon signed-rank test (α < 0.05) (Figures S4C-D). As previously reported,^23^ spontaneous remapping in most RF sessions was best captured by k = 2 models (Figure S4E). By contrast, we observed more variability in optimal k across SM sessions in all age groups due to heterogeneous remapping responses to context switches (Figure S4E).

Finally, we confirmed that the discreteness of remapping was comparable across age groups in two ways. First, at the same rank as the optimal k, we compared the relative cross-validated test performance of the k-means model and truncated singular value decomposition (tSVD), which is a form of uncentered Principal Components Analysis (PCA) in this case. As described in Low et al. (2021),^23^ the interpretation that network activity belongs to discrete maps was supported by the comparable performance of the more restrictive k-means model to the tSVD model, which permits trials to constitute linear combinations of spatial maps instead of discrete spatial maps. Component-matched tSVD and k-means performance were strongly correlated in all sessions in each age group in both tasks (Figure S5A). Additionally, we confirmed that within map mean pairwise trial similarity was significantly greater than across map pairwise trial similarity for all age groups in both tasks (Figure S5B). These results indicated that discrete maps indeed capture network-wide similarity patterns in both tasks.

#### Map label & identity assignment to assess context alignment

To address the fact that k-means cluster labels are arbitrarily assigned, we systematically re-labeled spatial maps. For SM sessions with optimal k ≥ 2, spatial maps were re-labeled based on their order of appearance in the task, such that map 1 was the map with the most epoch A. For k = 3 - 4 sessions, map 2 was the remaining map with the most epoch B trials. For k = 4 sessions, map 3 was the remaining map with more trials before the midpoint of the context-sorted alternation phase. RF spatial maps were re-labeled in order of their mean running speeds, such that map 1 had the slowest running speed. Remaps were identified as changes in map label between consecutive trials. To calculate remapping frequency, we divided remap count by trial count for the corresponding session or task epoch.

To assess the alignment of SM context and spatial map transitions, it was necessary to assign context identities (A or B) to each labeled map (see Figure 3A). After map labeling, map 1 dominated epoch A and was thus assumed to represent context A. Similarly, map 2 was assigned to represent context B. For sessions with optimal k > 2, maps 3 and 4 were each assigned to represent context A or B depending on whether they had greater relative mean pairwise similarity to epoch A or B. As such, it was possible to assign up to three maps to represent one context for k = 4 sessions. We then computed the fraction of “aligned” trials, on which VR context and spatial map identity matched, overall and for individual epochs.

#### Logistic regression models of by-trial responses

To model binary outcomes for each SM alternation trial (i.e. map-context alignment or a reward request), we used the logit method of the statsmodel.api package to perform logistic regression. In the model of map-context alignment probability by trial, we specified the following fixed effects as categorical variables: age group, the interaction of age group and sex, the interaction of age group and VR context (A/B), the interaction of age group and session, and the interaction of age group and the number of trials since a random VR context switch (0-11). To model reward request probability by trial, fixed effects included those above and the interaction of age group and remap occurrence as a categorical variable (No/Yes) and the interaction of age group and an aligned trial occurrence as a categorical variable (No/Yes). Unlike the linear mixed effects models, these models do not account for animal variance with random effects.

#### Local field potentials and power calculation

Local field potentials (LFP) were analyzed on one channel per session across both tasks (Figure 4). We selected the channel with the highest theta power after subsetting on the 200 channels closest to the probe tip with at least one “good” recorded cell. Power spectral density was calculated using Welch’s method from the scipy.signal package. Power in each frequency band (6 - 12 Hz for theta, 20 - 50 Hz for slow gamma, and 50 - 110 Hz for fast gamma) was calculated using a multitaper spectrogram using the spectral_connectivity package with a window of 0.5 seconds.^18^ To control for the effects of differences in running speed across age groups (see Figures S1B and S2B) on theta power,^66,67^ we up-sampled frequency band power traces (2 Hz) to the VR framerate (50 Hz) using a second order spline for interpolation and averaged power across timepoints where mice were running at intermediate speeds (20 ≤ speed ≤ 40 cm/s). We confirmed that this range collapsed mean and peak running speed differences across age groups (see Results).

#### Position decoding analysis

To predict the animal’s position from the spiking activity of all co-recorded cells, we fit spherically projected multivariate linear models^129^ as implemented by Low et al. (2021).^23^ This approach was selected because of the effective circularity of VR tracks. Predicting a circular variable, such as track position at a given time (y_t_ ∈ [0, 2π)), from linear covariates, such as the number of spikes fired by neurons n at times t (s_nt_), requires a circular linear regression model. These models are termed decoders as common practice.^130^ In this model, two regression coefficients, β_n_^(1)^ and β_n_^(2)^, are optimized for each neuron with an expectation maximization procedure previously described.^129^ The model estimate given set of test inputs after fitting to training data is as follows:

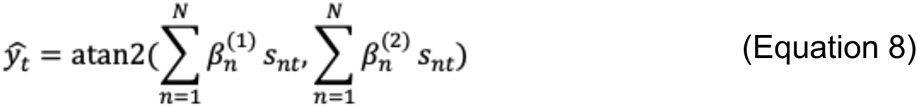

Here, atan2() corresponds to the two-argument arctangent function. Decoder score refers to the average of cos(y_t_ - ŷ_t_) over time bins in each session’s testing set, such that random position guesses over a circle would yield a score of 0 and perfect performance would have a score of 1.

For 10 repetitions per session, we fit decoding models to training sets comprising a randomly selected 90% of the network activity from RF sessions with >10 spatial cells and an optimal k = 2 and tested them on the held-out 10% of network activity. Synthetically ablating non-spatial cells from the network did not qualitatively alter decoder score differences across age groups, so we presented results from models trained and tested on spatial cell activity only (Figures 5H-J and S5E-F). Each datapoint in these figures represents the average decoder score across repetitions. For each session, training data were down-sampled to match spike number, position bins, running speed, and observations number across training sets drawn from the same session (e.g. map 1, map 2, both maps). We compared decoder scores to those for a spike time shuffle control for each session (see *Spatial firing rate & autocorrelation, shuffle procedure, and distance tuning*). Shuffle training sets were selected and down-sampled in the same manner. Since this shuffle disrupts spatial firing patterns, this control approximated chance decoder performance.

To further illustrate that distinct patterns of grid network activity occur across VR contexts in the SM task, we applied the same position decoding approach to training sets drawn from SM grid network activity across task epochs for all sessions with ≥ 10 grid cells (“Maps” A, B, A’ and B’ in Figure S5G-H). Decoders trained and tested on different epochs perform worse than those trained and tested on the same epochs (Figure S5G). Moreover, decoders trained and tested across context-mismatched epochs performed worse than those trained and tested across context-matched epochs (e.g. Train A / Test B’ vs. Train A / Test A’) (Figure S5H). Decoders struggled more to generalize across context-mismatched vs. -matched epochs, confirming that grid network activity was more conserved across matched vs. mismatched epochs.

#### Estimation of grid scale and unit recording depth

To compare the dorsal-ventral gradient of entorhinal grid cell scales,^12,17,74^ across age groups, we estimated grid cell scale and recording depth from brain surface along the probe using all sessions across tasks with dark trials (Figures 6F-J). Grid scale was calculated using the scipy.signal package’s find_peaks function to detect the location of the first peak in the autocorrelation on dark trials (width > 8 cm, height > 0.05, prominence > 0.05). We relaxed these peak detection criteria from those used to identify distance-tuning (see *Spatial firing rate & autocorrelation, shuffle procedure, and distance tuning*) to consistently select the first, instead of the largest, significant autocorrelation peak. To estimate each grid cell’s depth, we subtracted the median distance from the probe tip of the cell’s recorded spikes from the maximum inserted depth of the probe for that recording session.

#### Differential gene expression & correlational analysis

FASTQ files were mapped to the mouse transcriptome (mm39) that included nascent RNAs (introns included in the index) using Kallisto.^131^ The mapped reads were aggregated into a gene-based count matrix. Differential expression analysis was performed on the count matrix using DESeq2.^132^ Specifically, the likelihood ratio test was used for differential expression testing while controlling for sex-specific batch effects. A volcano plot of the differential expression results was generated with EnhancedVolcano (Figure 7B). A heatmap of the top 100 differentially expressed genes with row Z-score normalization between young and aged animals as determined by adjusted p-value was generated using pheatmap (Figure S7B).

To determine the relationship between aging gene expression and measures of neuronal activity, we performed correlational analysis between these measures. We determined the normalized expression of the gene sets that increase expression with aging (Aging Up, n = 170 genes expressed in every sample), decrease expression with aging (Aging Down, n = 260 genes expressed in every sample), and all aging genes (Aging All, n = 430 genes expressed in every sample) for each individual mouse. The Pearson correlation of these values with the LMM-fitted values of the neuronal activity measures was computed. The relationships between variables are represented in a correlation matrix (Figure 7E) generated using the corrplot package. To uncover individual genes whose expression is correlated with map stability, we used linear regression analysis to find significantly correlated genes. We present the number of correlated genes (both positive and negative), as well as their overlap with other variables in Figure 7F. Two examples of genes with significant correlations are presented in Figure 7G.

#### Gene ontology (GO) and gene set enrichment analysis (GSEA)

GO enrichment analysis was performed on several gene sets using Panther.^133^ Specifically, the biological process gene ontology sets were probed with genes differentially expressed during aging and those that significantly correlated with spatial firing stability. The top non-redundant biological process sets by false discovery rate (FDR) that had an enrichment of 1.25 above expected are shown in the plots (Figures 7C, 7H and S7C). GSEA^134,135^ was performed on the bulk neuronal RNA-seq. Briefly, the mouse-ortholog hallmark gene sets were probed with 21,388 expressed genes from the RNA-seq dataset for enrichment in the probed gene set of correlations between ages and expression. Gene set significance was determined using a permutation test and a threshold on family-wise error rate (FWER < 0.25) (see Figure 7C).

#### Single nuclei RNA-seq analysis

Cell Ranger was used to generate count matrices with include introns set to TRUE. Feature barcode count matrices were imported into Seurat v5.^121^ High-quality nuclei with a feature number greater than 200 and less than 7000, as well as mitochondrial content less than 0.25%, were kept for downstream processing. Both samples were integrated using the IntegrateData function after identifying integration anchors. The libraries were processed using the standard Seurat workflow to generate clusters. Subsequently, neurons were sub-clustered and dimensionality reduction was performed again, and new clusters were generated. These clusters were annotated based on marker expression (Figure S7D) found from FindAllMarkers in Seurat, and separated into excitatory and inhibitory neurons (Figure 7I), as well as layer specific excitatory neurons and inhibitory subclusters (Figure S7E).^136–139^

To determine the relationship between cell type and age related transcriptional changes, we generated module scores for genes that increase with age (n = 160 genes represented in snRNA-seq data) and decrease with age (n = 242 genes) using the AddModuleScore function in Seurat v5.^121^ These scores were overlaid onto the UMAPs of neurons (Figure S7F for age up and Figure S7H for age down), as well as used to determine the relative expression differences between excitatory and inhibitory neurons (Figures 7G,I). The Wilcoxon rank sum test with continuity correction was used to determine if there were significant differences in expression between the clusters. Similarly, we generated module scores for genes positively and negatively correlated with stability (n = 130 positive genes and n = 541 negative genes with an absolute correlation >0.6 that are represented in the snRNA-seq data) to investigate the relationship between stability and cell type. The correlation core module scores were overlaid onto UMAPs of neurons (Figures 7J, L) and used to determine expression differences between excitatory and inhibitory neurons (Figures 7K, M). The correlation core module scores were also determined for each neuronal cluster in Figure S7J. Additionally, the bulk neuronal RNA-seq age related changes of the stability correlation module genes are presented in Figure S7K.

To determine how markers of excitatory and inhibitory neurons change with age, we found marker genes for these two cell types in comparison to one another using the FindMarkers function in Seurat v5.^121^ The age-related changes of these gene sets in the bulk neuronal RNA-seq dataset are represented in Figure S7K.

## Supplemental Information

Figure S1, related to Figure 1.

Figure S2, related to Figure 1.

Figure S3, related to Figure 2.

Figure S4, related to Figure 3.

Figure S5, related to Figure 3.

Figure S6, related to Figure 6.

Figure S7, related to Figure 7.

## Acknowledgements

We thank A. Diaz for assistance with animal care; F.K. Masuda for providing training on surgeries, VR behavioral training, and neural recordings; C. Li for assistance with VR behavioral training; and I. Low and E. A. A. Jones for sharing code; and Giocomo Lab members for discussions and feedback. This work was supported by the Stanford University Medical Scientist Training Program (T32-GM007365 and T32-GM145402) and the National Institutes of Aging under a Ruth L. Kirschstein National Research Service Award (F30-AG079494) (to C.S.H); the Simons Foundation (SCPAB 811229) (to S.A.V. and L.M.G.); and the NIH Brain Initiatives (U19NS118284), NIMH (MH126904 and MH130452), the Simons Foundation (SCGB 542987SPI), NIDA (DA042012), the Vallee Foundation, and the James S. McDonnell Foundation (to L.M.G.).

## Author Contributions

C.S.H.: conceptualization, behavioral and neural recording data methodology, collection, and formal analysis, writing – original draft, writing – review & editing, and visualization; K.J.B.P.: transcriptomic data methodology, collection, and formal analysis, writing – original draft, visualization; J.M.S.: transcriptomic data methodology, collection, and formal analysis, writing – original draft, visualization; S.A.V.: conceptualization, writing – review & editing, supervision, and funding; L.M.G.: conceptualization, writing – review & editing, supervision, and funding.

