## Supplemental Legends & Figures S1 - S7 for "Spatial Coding Dysfunction and Network Instability in the Aging Medial Entorhinal Cortex"

#### Supplementary Figure Legends

*Figure S1. Aged mice employ divergent strategies from young and MA mice in the SM task.*

**(A)** Screenshots of the SM track from an aerial view on a context A (top) and context B (bottom) trial. Dotted lines outline the hidden reward locations in each context. A cue tower at 40 cm moved from left to right of the floor to indicate whether context A or B would appear. Black doors separated track halves at 200 and 400 cm but are obscured here. **(B)** Box and whisker plot of mean (left) and peak session running speed (right) across SM age groups (n = 54 young, 55 MA, 54 aged sessions). Running speed differed significantly across age groups (mean speed [cm/s]: young vs. MA vs. aged, 32.88 ± 0.73 vs. 28.99 ± 0.76 vs. 23.98 ± 0.86, Kruskal-Wallis test, H = 44.71, p = 1.95 x $10^{-10}$; peak speed [cm/s]: 76.58 ± 1.47 vs. 71.08 ± 1.59 vs. 59.52 ± 1.62, H = 43.53, p = 3.53 x $10^{-10}$). **(C)** Schematized visual acuity assessment trial structure where reward zone opacity is given by α (α = 0 if invisible vs. α = 1 if opaque) (left). Sample reward zones at four different alpha values (right). **(D)** Binned visual acuity assessment performance for SM mice (bin size = 0.1). Mouse visual acuity threshold, estimated as the center of the psychometric curve fitted to their task performance (see Methods), did not differ across age groups (center: young vs. MA vs. aged, 0.0981 ± 0.0007 vs. 0.0973 ± 0.0005 vs. 0.0975 ± 0.0004, Kruskal-Wallis test, H = 1.508, p = 0.4704). The highest estimated individual acuity threshold was 0.102, indicating all mice included in neural and behavioral datasets could see visual cues. Individual animals are plotted as pale dots jittered by age order. **(E)** Peak reward-triggered slowing (top left subpanels) and licking (top right subpanels) location, relative to the start of each reward zone, over sessions for each SM task phase. Age predicted later maximal licking but not slowing in the block phase only (LMM: Block Licking Location: Aged vs. Young, β = 13.964, p = 0.003; Slowing Location: Aged vs. Young, β = -3.190, p = 0.391). The LMM of peak alternation licking location did not converge. In the SM blocks, the prominence of mean reward-triggered slowing (bottom left) and licking (bottom right) did not differ by age group and increased over days (Slowing Magnitude: Aged vs. Young, β = -2.871, p = 0.451; Session, β = 2.737, p = 1.16 x $10^{-10}$; Licking Magnitude: Aged vs. Young, β = -0.545, p = 0.375; Session, β = 0.232, p = 0.003). Aged mice improved slowing but not licking magnitude less than young mice over days (Slowing Magnitude: Session x Aged, β = -1.205, p = 0.049; Licking Magnitude: Session x Aged, β = 0.050, p = 0.652). During alternation, slowing magnitude (bottom left) also improved less over sessions for aged mice (Session, β = 2.844, p = 1.383 x $10^{-13}$; Aged vs. Young, β = -3.536, p = 0.274; Session x Aged, β = -2.050, p = 3.09 x $10^{-7}$). Alternation licking magnitude (bottom right) did not change over days but was lower in aged mice overall (Aged vs. Young, β = -1.484, p = 0.019; Session, β = -0.052, p = 0.557; Session x Aged, β = 0.056, p = 0.656). Individual session data are plotted as dots, colored and jittered by age group. Bold lines represent age group means with age group SEM indicated by vertical bars. **(F)** Interaction of context identity (left, A vs. B) and trial structure (right, blocks vs. alternation) with age effects on SM task performance (fraction [frac.] of reward requested) by epoch, as fitted by LMM. Context B negatively predicted performance (B vs. A, β = -0.127, p = 6.333 x $10^{-5}$) but did not interact with age (Aged x B vs. Aged x A, β = -0.051, p = 0.150). The alternation trial structure negatively predicted performance in a manner that interacted significantly with age (Alternation [Alt.] vs. Block, β = -0.517, p = 5.188 x $10^{-24}$; Aged x Alt. vs. Aged x Block, β = -0.121, p = 0.001). Bold lines and dots indicate age group means with age group SEM indicated by vertical bars. Dots represent LMM-fitted performance for sessions epochs, colored and jittered by age group. **(G)** Box and whisker plot of the interaction of reward order (left; 270 vs. 370 cm context A reward location) and sex (right; female [F] vs. male [M]) with age effects on SM alternation performance (fraction [frac.] of reward requested), as fitted by LMM. Reward order did not predict performance overall (β = -0.046, p = 0.673) or among aged mice (270 x Aged vs. 370 x Aged: β = 0.118, p = 0.455; interaction effect omitted from subsequent models). This was also true for the effect of reward order on block phase performance (not shown; β = 0.012, p = 0.699; interaction effect omitted from final model). Male sex did not predict alternation performance (Male vs. Female, β = -0.101, p = 0.340), but male sex positively predicted aged performance (Aged x Male vs. Aged x Female, β = 0.371, p = 0.015). Age-sex effects on block performance were less apparent (not shown; Male vs. Female, β = -0.038, p = 0.489; Aged Male vs. Aged Female, β = 0.136, p = 0.111). Dots represent LMM-fitted performance for individual sessions, colored by age group. **(H)** Legend for dot colors in Figures S1B, 2D, S3D, 3B, S5D, and S5J. **(I)** Mean non-consummatory licks per trial by SM task phase over sessions across age groups, plotted as in S1E. Session negatively predicted block lick rate but not alternation lick rate via LMM (Blocks: Session, β = -0.812, p = 0.002; Alternation: Session, β = 0.042, p = 0.832). Being aged negatively predicted alternation but not block lick rate (Blocks: Aged vs. Young, β = -2.996, p = 0.154; Alternation: Aged vs. Young, β = -3.974, p = 0.006), confirming that aged mice fail during this task phase by missing rewards. Dots indicate individual session data, colored and jittered by age group. **(J)** Mean fraction (frac.) licks in the opposite (wrong) reward zone per trial by SM task phase over sessions as a measure of Type 1 lick error rate, plotted as in S1I. Session negatively predicted block but not alternation lick error rate (Blocks: Session, β = -0.009, p = 1.72 x $10^{-6}$; Alternation: Session, β = -0.003, p = 0.182). Being aged predicted neither alternation nor block lick error rate (Block: Aged vs. Young, β = -0.014, p =

0.429; Alternation: Aged vs. Young, β = -0.035, p = 0.068). This indicates that Type 1 errors are energetically penalized by the running wheel similarly across groups. Related to Figure 1.

*Figure S2. Random foraging behavior is spared in aged mice.*

**(A)** Screenshot of the RF track from an aerial view. Diamond checkerboard indicates the randomly appearing cued reward zone. **(B)** Mean (left) and peak (right) session running speed across age groups in the RF task, plotted as in Figure S1B (n = 43 young vs. 42 aged sessions). Mean speed was unchanged across age groups in the RF task (young vs. aged, 27.19 ± 0.8 vs. 29.96 ± 1.06, Wilcoxon rank sum test, p = 0.087), but peak running speed (right) was greater among young sessions (68.14 ± 1.58 vs. 61.72 ± 1.19, p = 0.0036). Individual dots represent session averages colored per the legend (far right) by mouse. **(C)** Mean running speed decreased in vs. outside of reward zones (left) for both RF age groups (n = 43 young, 42 aged model pairs; speed decrease (cm/s): young vs. aged, 10.89 ± 1.31 vs. 13.41 ± 1.61; Wilcoxon signed-rank test, $p = 8.43 \times 10^{-11}$ & $p = 9.41 \times 10^{-11}$). Fraction of trial licks was greater in vs. outside of reward zones for young and aged sessions (trial lick fraction increase: 0.5 ± 0.06 vs. 0.47 ± 0.06; $p = 2.3 \times 10^{-11}$ & $p = 5.18 \times 10^{-9}$). Pale dots indicate session averages, colored by age group, and large, bold dots indicate animal averages per the legend. **(D)** Mean reward zone-triggered slowing (left) and fractional (frac.) licking increase (right), plotted as in Figure S2B. Equivalent reward-triggered slowing (Wilcoxon rank sum test, p = 0.34) and licking (Wilcoxon rank sum test, p = 0.54) across RF age groups indicates intact foraging behavior in aging. **(E)** Speed (left subpanels) and lick count (right subpanels) near reward zones on individual reward trials (light gray) for the first (left) and last (right) sessions from representative young (top row) and aged (bottom row) mice. Colored lines indicate the session average reward-triggered behavior. Stereotypy of reward-triggered slowing and licking increases over days, reflected by the larger troughs in mean speed and peaks in mean licking. **(F)** Reward-triggered maximal slowing (top left) and licking (top right) location over sessions is equivalent across age groups per LMMs (Slowing Location: Aged vs. Young, β = 8.235, p = 0.235; Session, β = -0.862, p = 0.375; Licking Location: Aged vs. Young, β = 7.252, p = 0.332; Session, β = -1.301, p = 0.264). The prominence of mean reward-triggered slowing (bottom left) and licking (bottom right) is also equivalent across age groups over days as fitted by LMMs (Slowing Magnitude: Aged vs. Young, β = -0.918, p = 0.755; Session, β = 3.049, $p = 2.0 \times 10^{-5}$, Session x Aged, β = -0.855, p = 0.388; Licking Magnitude: Aged vs. Young, β = -0.453, p = 0.645; Session, β = 0.513, $p = 1.95 \times 10^{-5}$; Session x Aged, β = -0.160, p = 0.3330). This indicates RF behavior becomes more stereotyped in both age groups over sessions, contrasting SM alternation behavior (see Figure S1E). **(G)** The mean fraction (frac.) of trial licks outside the reward zone, plotted as in Figure S2B, did not differ across age groups (young vs. aged, 0.14 ± 0.02 vs. 0.16 ± 0.02, Wilcoxon rank sum test, $p = 0.1759$). This confirms that Type 1 error rates are similar across age groups in both tasks (see Figure S1J). Related to Figure 1.

*Figure S3: Non-grid spatial cell activity is less dynamic across VR contexts and over days.*

**(A)** As in Figure 2A, raster plots for NGS cell activity. **(B)** As in Figure 2B, for NGS cells in (A) revealing a lack of distance tuning. This is indicated by the lack of large peaks in the dark autocorrelation of each cell. **(C)** As in Figure 2C, for NGS cells. Dark autocorrelation peak prominence from NGS cell vs. shuffle activity differed significantly (n = 6,559 model pairs, mean ± SEM prominence, NGS vs. shuffle activity, 0.047 ± 0.001 vs. 0.011 ± 0.0007, Wilcoxon signed-rank, $p < 0.0001$) (see Methods). However, NGS cell dark autocorrelation peak prominence was significantly lower than that of grid cells (NGS vs. grid activity, 0.047 ± 0.001 vs. 0.25 ± 0.001, Wilcoxon rank sum, $p < 0.0001$). **(D)** As in Figure 2D, for NGS cells. NGS cell density by session does not differ among SM age groups (% NGS cells, young vs. MA vs. aged, 14.59 ± 0.87 vs. 15.73 ± 0.95 vs. 15.14 ± 1.08, H = 0.48, $p = 0.79$). **(E)** As in Figure 2E, for NGS cells in (A) from left to right. Qualitatively, NGS cells display more similar spatial firing across trials and task epochs than grid cells. **(F)** Probability density of the spatial firing coherence (left) and sparsity (right) of NGS cell activity in epoch A vs. shuffle activity. The spatial firing coherence and sparsity of classified NGS cells' activity significantly exceeds that of their shuffled activity, pooling cells across age groups (n = 6,559 model pairs; coherence, NGS vs. shuffle, 0.74 ± 0.002 vs. 0.22 ± 0.002, Wilcoxon signed-rank, $p < 0.0001$; sparsity, NGS vs. shuffle, 1.1 ± 0.009 vs. 0.72 ± 0.002, $p < 0.0001$). The average 99th percentile of shuffle coherence and sparsity scores were 0.4705 ± 0.0009 and 1.3739 ± 0.0118 (n = 43,388 total SM cells). Bin sizes were 0.01. **(G)** Schematized SM gain manipulation, indicating 200 trials with a fixed relationship between visual and motor feedback (top, ‘‘normal trials’’) and 20 trials that required the animal to run 1.42x as far to travel the same VR distance (bottom, ‘‘gain change trials’’). **(H)** Cumulative density function (CDF) of SM gain change responses, computed as the mean pairwise similarity change across gain change and non-gain change trials (see Methods), in young, MA, and aged (left to right) grid vs. NGS cells on A` trials (top) and B` trials (bottom). In each age group, grid cells responded stronger than NGS cells to A` gain change trials (gain change response, grid vs. NGS cells, Young: 0.19 ± 0.003 vs. 0.14 ± 0.002, Wilcoxon rank sum test, $p = 5.2 \times 10^{-25}$; MA: 0.20 ± 0.003 vs. 0.16 ± 0.002, $p = 1.2 \times 10^{-14}$; Aged: 0.22 ± 0.004 vs. 0.15 ± 0.002, $p = 3.8 \times 10^{-39}$). The same was true for B` gain responses (Young: 0.19 ± 0.003 vs. 0.14 ± 0.002, $p = 1.9 \times 10^{-25}$; MA: 0.21 ± 0.003 vs. 0.16 ± 0.002, $p = 4.4 \times 10^{-26}$; Aged: 0.20 ± 0.003 vs. 0.14 ± 0.002, $p = 4.4 \times 10^{-31}$). This validates grid vs. NGS cell classification by distance-tuning during dark running. Aged animals' grid cells exhibited stronger gain responses in both contexts (A`: Aged vs. Young, β = 0.065, p = 0.017; B`: Aged vs. Young, β = 0.061, p = 0.034). **(I)** As in Figure 2F, mean NGS cell stability during epochs A` (left) and B` (right) over sessions in each age group. **(J)** As in Figure 2G, for alternation (alt.) performance (fraction [frac.] reward requested) vs. mean NGS cell alteration stability (n = 54 young, 58 MA, and 55 aged sessions overall; n = 9 young, 9 MA, and 8 aged final sessions). Across days, alternation performance related to NGS stability across ages (r = 0.43, p = 4.1 x $10^{-9}$) and among aged sessions alone (r = 0.43, p = 0.0010). Final session alternation performance and mean NGS stability related overall (r = 0.65, p = 0.00034) but not among aged sessions only (r = 0.61, p = 0.10). **(K)** As in Figure 2H and (I), for mean NGS cell context-matched epoch similarity. The model of B x B` NGS similarity converged only using a maximum-likelihood estimator. **(L)** As in Figure 2I and (J), for mean NGS cell similarity across context-matched task epochs. This related to alternation performance across all ages (r = 0.38, p = 3.1 x $10^{-7}$) and among aged sessions (r = 0.35, p = 0.0093). Final session alternation performance and mean context-matched epoch similarity related across all ages (r = 0.54, p = 0.0054) but not among aged sessions (r = 0.64, p = 0.0906). **(M)** As in Figure 2J and (I), for mean NGS cell similarity ratio. **(N)** As in Figure 2K and (J), for mean NGS cell similarity ratio. This related to alternation performance across all ages (r = 0.47, p = 8.9 x $10^{-11}$) and among aged sessions (r = 0.49, p = 0.0002). Final session alternation performance and similarity ratio related for all ages (r = 0.60, p = 0.0012) but not among aged sessions (r = 0.64, p = 0.087). Related to Figure 2.

*Figure S4: K-means clustering with k optimization identifies grid network spatial maps.*

**(A)** Illustrated k hyperparameter optimization procedure for example young SM sessions fit by k-means models with optimal k = 2, 3, and 4 (from left to right) (see Methods). Subpanel titles indicate mouse and session number, if optimal k was greater than 1, and optimal k. The VR context (top x-axis) and k-means-labeled (right y-axis) trial-by-trial grid network similarity matrix using optimal k (top left) and 10-fold cross-validation test $R^2$ for each possible k for real vs. shuffled firing rate tensors (top right), and average (avg.) silhouette (sil.) score for possible k ≥ 2 on those 10 repetitions (bottom left; SEM shaded) are shown. The optimal k-means model captured similarity matrix structure (top left), yielded significantly greater test $R^2$ than when applied to shuffle data (top right), and maximized the average silhouette score (bottom left). Furthermore, it produced high quality clusters with distributions of trial silhouette scores that intersected the average silhouette score (black dashed vertical line; bottom right). **(B)** As in (A) but for example sessions with optimal k > 4 (young, MA, and aged from left to right) selected during the sweep to set the maximum possible k (see Methods). In brief, as before, we selected the k that maximized average silhouette score across clusters among possible 2 ≤ k ≤ 8 over 10 repetitions of a cross-validation procedure (bottom left; SEM shaded). For each possible k, we compared 10-fold cross-validation train vs. test $R^2$ (top right) and the distribution of trial silhouette scores for each map vs. the average silhouette score (bottom right). In each example, at least one cluster had no trials with silhouette scores greater than or equal the average, indicating a low-quality cluster. Since these subpanels do not reflect selected models, the grid similarity matrix (top left) is unlabeled. **(C)** As in (A), for an example SM session fitted poorly by k > 2 ("One Map = True"). At the optimal k, the model's mean test $R^2$ for the real vs. shuffle firing rate tensors did not differ (one-sided Wilcoxon signed-rank test, $p > 0.05$). Since this session was not fit by k > 2, the grid network similarity matrix (top left) is unlabeled. **(D)** As in (C), for an example RF session not fit by k > 2. Here, a spatial cell network similarity matrix (top left) is shown unlabeled. **(E)** Relative density of optimal k values for RF sessions with > 10 spatial cells (left; n = 37 young & 39 aged) and for SM sessions with ≥ 10 grid cells (right; n = 46 young, 47 MA & 41 aged). The optimal k distribution did not differ significantly across age groups for either task (RF: 2-sample Kolmogorov-Smirnov test, A vs. Y: D = 0.023, p = 0.25, sign = -1, loc = 2.0; SM: A vs. Y: D = 0.13, p = 0.79, sign = -1, loc = 2.0; MA vs. Y: D = 0.077, p = 0.99, sign = 1, loc = 1.0; A vs. MA: D = 0.18, p = 0.37, sign = -1, loc = 2.0). Consistent with previous work,[23] most RF sessions were fit by k = 2 models. **(F)** RF spatial cell network trial-by-trial similarity matrices with spatial maps labeled by k = 2 - 4 models (left to right) from example sessions in each age group. The matrices omit dark and gain trials, and the right axis gives k-means map labels for each trial with arbitrary color order. Labeled matrices reveal similarity structure well captured by optimized k-means models. Subpanel titles indicate mouse, session number, and network spatial cell count. Color bar indicates trial-by-trial spatial correlation value. **(G)** As in Figure 3A and (F), for SM grid network trial-by-trial similarity matrices, for which similarity structure is also well captured by optimized k-means models. VR context (top axis: context A [pink], context B [dark blue]) contrasts with k-means spatial map labels for each trial (right axis: map dominating context A trials [pink]; map dominating context B trials [dark blue], color of remaining maps, green or black arbitrarily). Each labeled map was also assigned a context identity (see Methods). Y27_6, MA1_6, and A19_6 map labels contrast with map identities in Figure 3A. Subpanel titles indicate mouse, session number, and network grid cell count. **(H)** As in Figure 3B, remapping frequency of RF spatial cell networks (n = 37 young, 39 aged sessions). Remapping frequency did not differ between age groups in the RF task (young vs. aged, 0.0078 ± 0.0006 vs. 0.0122 ± 0.0018, Wilcoxon rank sum test, p = 0.40). **(I)** Alternation (alt.) performance (fraction [frac.] reward requested) vs. remapping frequency (left) or optimal k value (right) for all SM sessions across age groups (n = 47 young, 46 MA, and 41 aged sessions). Remapping frequency did not relate to alternation performance across all sessions (r = -0.14, p = 0.11) or among aged sessions (r = -0.15, p = 0.35). K value also did not relate to alternation performance across all sessions (r = -0.11, p = 0.22) or among aged sessions (r = -0.045, p = 0.78). Plotted as in the top of Figure 2G, 2I, 2K, and 3D. Related to Figure 3.

*Figure S5: Remapping discreteness and coordination is preserved in aging, resulting in the conservation of positional information across maps and contexts.*

**(A)** Rank-matched k-means vs. truncated singular value decomposition (tSVD) model performance (test $R^2$) for RF sessions (left; n = 39 young, 39 aged) and for SM sessions (n = 48 young, 51 MA, & 45 aged) (right). Dot size scales with optimized k (k = 2 - 4). tSVD and k-means model performance related strongly for each age group across tasks (RF: Young, r = 0.98, p = $2.45 \times 10^{-28}$, Aged, r = 0.99, p = $1.16 \times 10^{-31}$; SM: Young, r = 0.98, p = $2.56 \times 10^{-34}$; MA, r = 0.98, p = $1.54 \times 10^{-37}$; Aged, r = 0.98, p = $7.15 \times 10^{-31}$). This indicates that k-means model assumptions of spatial maps comprising discrete trials are appropriate (see Methods). **(B)** Mean pairwise spatial similarity for trials within vs. across spatial maps for each RF session by age group where optimal k = 2 (left) and SM session where k ≥ 2 (right). Both RF age groups exhibited a significant mean similarity decrease (n = 27 young, 34 aged pairs; % mean similarity decrease, young, 42.85% ± 3.81%, Wilcoxon signed-rank test, young, p = $1.16 \times 10^{-10}$; aged, 37.35 ± 3.51%, p = $1.49 \times 10^{-8}$). There was no difference in the change in mean similarity across RF age groups (Wilcoxon rank sum test, p = 0.40). All SM groups also exhibited significant change in mean similarity (n = 47 young, 46 MA, and 41 aged pairs; % similarity decrease, young vs. MA vs. aged, 53.95 ± 2.38% vs. 48.23 ± 2.51% vs. 49.53 ± 2.14%). There was also no significant difference in the change in similarity across SM groups (Kruskal Wallis test, H = 4.17, p = 0.12). Dots represent individual session data, and black bars indicate the mean across sets of plotted sessions. **(C)** Remapping coordination in RF sessions with at least three spatial cells (n = 42 young, 42 aged), computed as mean correlation between cell and network trial-by-trial similarity matrices (see Methods), plotted as in Figure S2B. Remapping coordination did not differ across age groups (young vs. aged, 0.318 ± 0.006 vs. 0.317 ± 0.005, Wilcoxon rank sum test, p = 0.26). **(D)** As in (C) for SM sessions with at least three grid cells, plotted as in Figure 2D. Remapping coordination did not differ across age groups (n = 52 young, n = 57 MA, and n = 51 aged, young vs. MA vs. aged, 0.415 ± 0.014 vs. 0.4 ± 0.011 vs. 0.403 ± 0.013, Kruskal-Wallis test, H = 0.54, p = 0.76). **(E)** Decoder performance on spatial cell activity from all RF sessions with k = 2 across all possible training/testing map combinations (left to right) (n = 39 young, n = 39 aged for all), plotted as in

Figure 5H. Aged session decoder scores were lower than young ones for all train/test combinations (score, Map 1/1 young vs. aged, 0.6541 ± 0.036 vs. 0.466 ± 0.0361, Wilcoxon rank sum test, $p = 0.00096$; Map 2/2, 0.633 ± 0.039 vs. 0.4734 ± 0.036, $p = 0.0052$; Map ½, 0.4208 ± 0.0436 vs. 0.2533 ± 0.0295, $p = 0.0095$; Map 2/1, 0.4469 ± 0.0447 vs. 0.2401 ± 0.0281, $p = 0.00064$). Dashed line indicates mean shuffle score for each map combination and age group. **(F)** Decoder performance trained and tested within maps in the RF track was greater than when trained and tested across maps (n = 78 model pairs each; young train/test same vs. different, Wilcoxon signed-rank test, $p = 1.10 \times 10^{-14}$; aged train/test same vs. different, $p = 4.37 \times 10^{-14}$). This indicates that spatial network positional information is discretized across RF spatial maps in both age groups. Dots indicate session values colored by age group. **(G)** As in (F), decoder score when trained and tested within vs. across contexts for the SM block phase. Decoder performance decreased when trained and tested across vs. within contexts for each age group (n = 96 young model pairs, 0.7076 ± 0.0197 vs. 0.164 ± 0.0243, Wilcoxon signed-rank test, $p = 9.8 \times 10^{-18}$; n = 102 MA model pairs, 0.6623 ± 0.0168 vs. 0.1928 ± 0.0188, $p = 1.7 \times 10^{-18}$; n = 90 aged model pairs, 0.6225 ± 0.0226 vs. 0.128 ± 0.0156, $p = 1.6 \times 10^{-16}$). Within each age group, decoder score did not differ across contexts if train and test contexts matched (mean ± SEM train/test A vs. B score, n = 48 young, 0.7333 ± 0.0267 vs. 0.6818 ± 0.0287, Wilcoxon rank sum test, $p = 0.15$; n = 51 MA, 0.6429 ± 0.0255 vs. 0.6817 ± 0.0218, $p = 0.27$; n = 45 aged sessions, 0.6214 ± 0.0316 vs. 0.6238 ± 0.0325, $p = 0.74$). This indicates that grid network positional information is discretized across SM contexts for all age groups. **(H)** As in (G), decoder score when trained and tested across context-matched vs. -mismatched SM task epochs. Decoder performance was greater when trained and tested on context-matched (A x A`, B x B`) vs. -mismatched (A x B`, B x A`) epochs for each age group (n = 96 young model pairs, 0.3842 ± 0.0256 vs. 0.2206 ± 0.0233, Wilcoxon signed-rank test, $p = 2.7 \times 10^{-16}$'; n = 102 MA model pairs, 0.4093 ± 0.0196 vs. 0.2234 ± 0.0158, $p = 1.9 \times 10^{-16}$, and n = 90 aged model pairs, 0.3529 ± 0.0230 vs. 0.175 ± 0.0160, $p = 2.0 \times 10^{-15}$). Within each age group, decoder score was greater if trained and tested on B x B` than on A x A` contexts (mean ± SEM train/test A x A` vs. B x B` score, n = 48 young, 0.3054 ± 0.0369 vs. 0.4630 ± 0.0319, Wilcoxon rank sum test, $p = 0.0021$; n = 51 MA, 0.3269 ± 0.0266 vs. 0.4916 ± 0.0239, $p = 3.2 \times 10^{-5}$ ; n = 45 aged sessions, 0.2834 ± 0.0284 vs. 0.4225 ± 0.0332, $p = 0.0016$). These results underscore that grid network positional information is discretized across SM contexts in different task phases for all age groups. **(I)** Example SM grid network similarity matrices generated from activity in the front (0 - 200 cm) vs. back (200 - 400 cm) of the VR track, omitting dark trials, including gain trials, and sorting alternation trials by context. In each case, the back track matrix exhibited greater context-specific structure compared to the front track matrix.

**(J)** Correlation of front vs. back track SM grid similarity matrices (n = 48 young, 51 MA, and 45 aged sessions), plotted as in Figure 2D. This correlation differed across age groups (young vs. MA vs. aged, 0.66 ± 0.02 vs. 0.57 ± 0.01 vs. 0.58 ± 0.02, Kruskal-Wallis test, H = 17.0, p = 0.00021). Young sessions exhibited greater correlation compared to MA or aged sessions (Conover post-hoc correction, young vs. MA, p = 0.00014, young vs. aged, p = 0.0046, MA vs. aged, p = 0.33). To account for this difference in the timing of context recognition during trials, all analyses in Figure 3 were performed on back of track matrices. Related to Figure 3.

*Figure S6. Increased speed gain & speed tuning instability also occur in aged mouse fast-spiking interneurons, conjunctive grid-speed cells, & speed-only cells.*

**(A)** Classification of putative inhibitory interneurons (INs) (light blue) vs. excitatory (EX) cells (lime green) from an example aged session (see Methods). Putative INs vs. excitatory cells clustered in waveform duration and peak:trough ratio space and were well-separated by a waveform duration threshold (left) (dashed black line). Histograms of session IN vs. excitatory cells by waveform width (ms) (middle) and mean firing rate (FR [Hz]) (right) confirmed that INs exhibited narrower waveforms and higher FR. Colors for INs vs EX cells maintained throughout. **(B)** Probability density of the waveform duration of classified EX cells vs. INs (light blue) (n = 49,193 EX cells and 15,123 INs). Waveform duration EX cells exceeded that of INs (EX vs. INs, 0.5807 ± 0.0005 vs. 0.2432 ± 0.0005, Wilcoxon rank sum test, $p < 0.0001$). Dashed black line indicates the IN waveform duration threshold. A small minority of INs exceeded the duration threshold, since they were classified by the secondary FR threshold (> 40 Hz) (see Methods). Bin size was 0.025 ms. **(C)** As in (B) for waveform peak:trough ratio (PTR), which also differed significantly across cell types (EX vs. INs, 0.4445 ± 0.0010 vs. 0.6245 ± 0.0034, $p < 0.0001$). Bin size was 0.025. **(D)** Differences in IN density (first; n = 98 young, 58 MA, & 97 aged sessions) and waveform and FR properties (n = 97 young, 58 MA, 97 aged sessions) across age groups, including mean session waveform duration (second), PTR (middle), mean waveform halfwidth (fourth), and mean FR (last). IN density differed across age groups (% INs, young vs. MA vs. aged, 21.22 ± 0.92 vs. 27.35 ± 0.81 vs. 26.24 ± 0.87, Kruskal Wallis test, H = 39.21, $p = 3.05 \times 10^{-9}$; post-hoc Conover test, young vs. MA, $p = 1.88 \times 10^{-8}$; young vs. aged, $p = 8.11 \times 10^{-7}$). As did IN waveform duration (0.2481 ± 0.0018 vs. 0.2406 ± 0.0019 vs. 0.2402 ± 0.0016, H = 22.59, $p = 1.24 \times 10^{-5}$; young vs. MA, p = 0.00014; young vs. aged, p = 0.000063). IN waveform PTR did not differ with age (0.6281 ± 0.0121 vs. 0.5905 ± 0.0083 vs. 0.6166 ± 0.0079, H = 5.83, p = 0.054; young vs. MA, p = 0.059; young vs. aged, p = 0.66). IN waveform halfwidth differed significantly with aging (0.1503 ± 0.0015 vs. 0.1431 ± 0.0013 vs. 0.1433 ± 0.0012, H = 21.5, $p = 2.16 \times 10^{-5}$; young vs. MA, p = 0.00015, young vs. aged, p = 0.00014). Finally, IN mean FR differed significantly across age groups (9.0683 ± 0.4352 vs. 11.4235 ± 0.424 vs. 10.7849 ± 0.4427, H = 18.58, p = 9.22 x $10^{-5}$; young vs. MA, p = 0.000062, young vs. aged, p = 0.0107). These results collectively indicate possible differences in the relative density of MEC IN subtypes or changes in MEC IN function with age. Plotted as in Figures 4C and 6D-E. **(E)** Density of various speed-tuned cell populations: + and - speed cells (first, second), + and - non-spatial speed cells (third, fourth), speed-tuned interneurons (fifth); and speed-tuned grid cells (last) by age group (all except for last: n = 98 young, 58 MA, & 97 aged sessions; last: n = 82 young, 58 MA, & 67 aged sessions). + and - speed cell density did not differ with age (% cells + speed cells, young vs. MA vs. aged, 16.67 ± 0.77 vs. 16.13 ± 0.64 vs. 18.41 ± 0.95, Kruskal Wallis test, H = 1.71, p = 0.42; post-hoc Conover test, young vs. MA, p = 0.82; young vs. aged, p = 0.76; % cells - speed cells, 10.11 ± 0.6 vs. 11.02 ± 0.7 vs. 10.52 ± 0.74, H = 2.21, p = 0.33; young vs. MA, p = 0.54; young vs. aged, p = 0.99). Neither did the density of + and - non-spatial speed cells (% + speed only cells, 4.63 ± 0.41 vs. 3.96 ± 0.35 vs. 6.88 ± 0.76, H = 4.00, p = 0.14; young vs. MA, p = 0.84; young vs. aged, p = 0.26; % - speed only cells, 3.24 ± 0.3 vs. 3.59 ± 0.34 vs. 4.18 ± 0.39, H = 2.99, p = 0.37; young vs. MA, p = 0.82; young vs. aged, p = 0.35). + speed-tuned IN density increased starting in middle age (% + speed INs, 6.92 ± 0.35 vs. 10.97 ± 0.63 vs. 9.39 ± 0.48, H = 29.1, p = 4.90 x $10^{-7}$; young vs. MA, p = 3.99 x $10^{-7}$; young vs. aged, p = 2.27 x $10^{-4}$). Finally, + speed-tuned grid cell density was uniform across age groups (% + speed grid cells, 5.50 ± 0.38 vs. 4.64 ± 0.3 vs. 5.72 ± 0.55, H = 1.34, p = 0.51; young vs. MA, p = 0.75; young vs. aged, p = 0.95). These results suggest that differences in speed cell density are unlikely to account for altered speed tuning with age (see Figure 6D-E). Plotted as in Figures 4C and 6D-E. **(F)** As in Figure 6D, mean FR-speed slope, or speed gain, for + non-spatial speed cells (left, n = 88 young, 56 MA, 94 aged sessions), - non-spatial speed cells (second, n = 89 young, 51 MA, 92 aged sessions), + speed-tuned INs (third, n = 94 young, 58 MA, 95 aged sessions), and + speed-tuned grid cells (right, n = 77 young, 57 MA, 61 aged sessions). Speed gain became more extreme in + and - non-spatial speed cells in aged vs. middle aged sessions (+ speed only: slope, young vs. MA vs. aged, 0.0412 ± 0.0037 vs. 0.033 ± 0.0034 vs. 0.0466 ± 0.0035, Kruskal-Wallis test, H = 9.68, p = 0.0079; post-hoc Conover test, young vs. MA, p = 0.0868, young vs. aged, p = 0.2100, MA vs. aged, p = 0.0053; - speed only: -0.0309 ± 0.0042 vs. -0.0198 ± 0.0018 vs. -0.0344 ± 0.0033, H = 12.27, p = 0.0022; young vs. MA, p = 0.0899, young vs. aged, p = 0.0713, MA vs. aged, p = 0.0016). Speed gain was also increased among + speed INs and + speed grid cells in aged vs. young sessions (+ speed INs: 0.1458 ± 0.0057 vs. 0.1588 ± 0.0083 vs. 0.1857 ± 0.0082, H = 13.24, p = 0.0013; young vs. MA, p = 0.2307, young vs. aged, p = 0.0008; + speed grid cells: 0.0443 ± 0.0015 vs. 0.054 ± 0.0024 vs. 0.0515 ± 0.003, H = 13.43, p = 0.0012; young vs. MA, p = 0.0011, young vs. aged, p = 0.0241). **(G)** As in Figure 6E and (F) for mean trial stability speed score across speed-tuned cell types. Speed tuning stability over trials decreased after middle age for + speed only cells (score, young vs. MA vs. aged, 0.1944 ± 0.0085 vs. 0.2269 ± 0.0116 vs. 0.1739 ± 0.0067, Kruskal-Wallis test, H = 14.35, p = 0.0008; post-hoc Conover test, young vs. MA, p = 0.0520, young vs. aged, p = 0.0667, MA vs. aged, p = 0.0004). This was not true for - speed only cells (-0.2124 ± 0.0121 vs. -0.2104 ± 0.0115 vs. -0.207 ± 0.0114, H = 1.1, p = 0.57; young vs. MA, p = 1.0, young vs. aged, p = 1.0). + speed INs speed tuning stability decreased in aged vs. young sessions (0.2813 ± 0.0072 vs. 0.3105 ± 0.0076 vs. 0.2623 ± 0.0075, H = 19.48, p = $5.89 \times 10^{-5}$; young vs. MA, p = 0.0156, young vs. aged, p = 0.0316). Similarly, + speed grid cells tuning stability decreased in aged vs. young sessions (0.1804 ± 0.0069 vs. 0.2086 ± 0.0075 vs. 0.1514 ± 0.0054, H = 28.68, p = $5.92 \times 10^{-7}$; young vs. MA, p = 0.0014, young vs. aged, p = 0.0081). Related to Figure 6.

*Figure S7. Cell type- and layer-specificity of transcriptomic changes in the aging MEC*

**(A)** Fluorescence-activated cell sorting (FACS) gating strategy to isolate neuronal nuclei dissociated from MEC (see Methods). Briefly, nuclei were gated by forward (FSC-A) and side (SSC-A) scatter to remove debris (left top) and by height (FSC-H) and size (FSC-A) to remove doublets (left bottom). Hoechst+ and NeuN+ nuclei were then isolated iteratively (middle, right). Subplots are labeled with the type and percent of nuclei isolated at each sorting step. **(B)** Heatmap of the top 100 most differentially expressed genes as determined by adjusted p-value with an absolute fold change of Log2 > 1 (n = 7 young, 7 aged mice; Wald test followed by Benjamini and Hochberg multiple hypothesis correction). Color bar indicates Z-score of expression level with age (red = increased with age; blue = decreased with age). **(C)** Top ten biological processes from Gene Ontology (GO) enrichment analysis on the combination of positive & negative stability-correlated module genes (Fisher's Exact test followed by FDR calculation). **(D)** Dot plot of top representative cell marker genes used to distinguish clusters in snRNA-seq data. **(E)** UMAP plot of neurons from MEC snRNA-seq with neuronal subtype annotations layered on or above their respective clusters. If known, excitatory cell (Exc.) clusters are labeled with their gross cortical layer (e.g. '23' for 2/3 or '56' for 5/6) and numbered arbitrarily within the layer group. Excitatory cell clusters of unknown layer (Exc Uns) are numbered arbitrarily. Inhibitory IN clusters (Inh) are labeled with their primary cellular marker gene (e.g. *Pvalb*, *Sst*, *Vip*). A cluster corresponding to subiculum excitatory cells was also identified (Exc Sub). **(F)** UMAP plot with the relative expression of the gene module that increases expression during aging ("Age Up") (n = 160 genes represented in snRNA-seq data). Color bar indicates expression relative to control feature sets.

Y-axis is shared with (E). **(G)** Violin plot of the relative expression of the age increased gene module between INs and excitatory (EX) neurons. Age-increased genes are enriched among INs (-0.0233 ± 0.0003 vs. -0.0179 ± 0.0006, EX vs. IN, Wilcoxon rank sum test, $p < 0.0001$). **(H)** As in (F), for the gene module that decreases expression during aging (“Age Down”) (n = 242 genes). **(I)** As in (G), for the age decreased gene module. Age-decreased genes are enriched among excitatory cells (EX) (0.1682 ± 0.0003 vs. 0.1215 ± 0.0006, EX vs. IN, Wilcoxon rank sum test, $p < 0.0001$). (Wilcoxon rank sum test, $p < 0.0001$). **(J)** Violin plots of positive (top) & negative (bottom) stability-correlated core gene module relative expression across identified neuronal clusters. Expression of the positive correlation module is enriched in some Layer 2/3 and 5/6 excitatory cell clusters while expression of the negative correlation module is enriched among IN clusters. Dashed line indicates equivalent relative expression to the UMAP dataset. Colors correspond to labeled clusters in (E). **(K)** Violin plots of excitatory neuron (EX) (n = 323 genes) and IN (n = 199 genes) marker gene expression change with age (top) (Wilcoxon signed-rank test, **** $p<0.0001$) and positive and negative stability core gene modules expression change with age (bottom) in bulk expression data. EX cell marker expression changed with age (-0.2808 ± 0.02134, Wilcoxon signed-rank test, **** $p < 0.0001$), but IN marker expression did not (0.0008763 ± 0.02899, Wilcoxon signed-rank Test, $p = 0.7953$). Positive stability-correlated gene module expression decreased with age (-0.3395 ± 0.04431, Wilcoxon signed-rank test, **** $p < 0.0001$), and negative stability-correlated gene module expression increased with age (0.4383 ± 0.02449, Wilcoxon signed-rank test, **** $p < 0.0001$). Cross-variable comparisons within or across subpanels should not be made given different gene set sizes and expression levels. Dashed lines indicate equivalent expression across ages. Dotted lines indicate medians. Data are represented as the log2 fold change (FC) of aged over young expression. Lower opacity emphasizes that these data reflect bulk RNA-seq gene expression. Related to Figure 7.

#### Figure S1

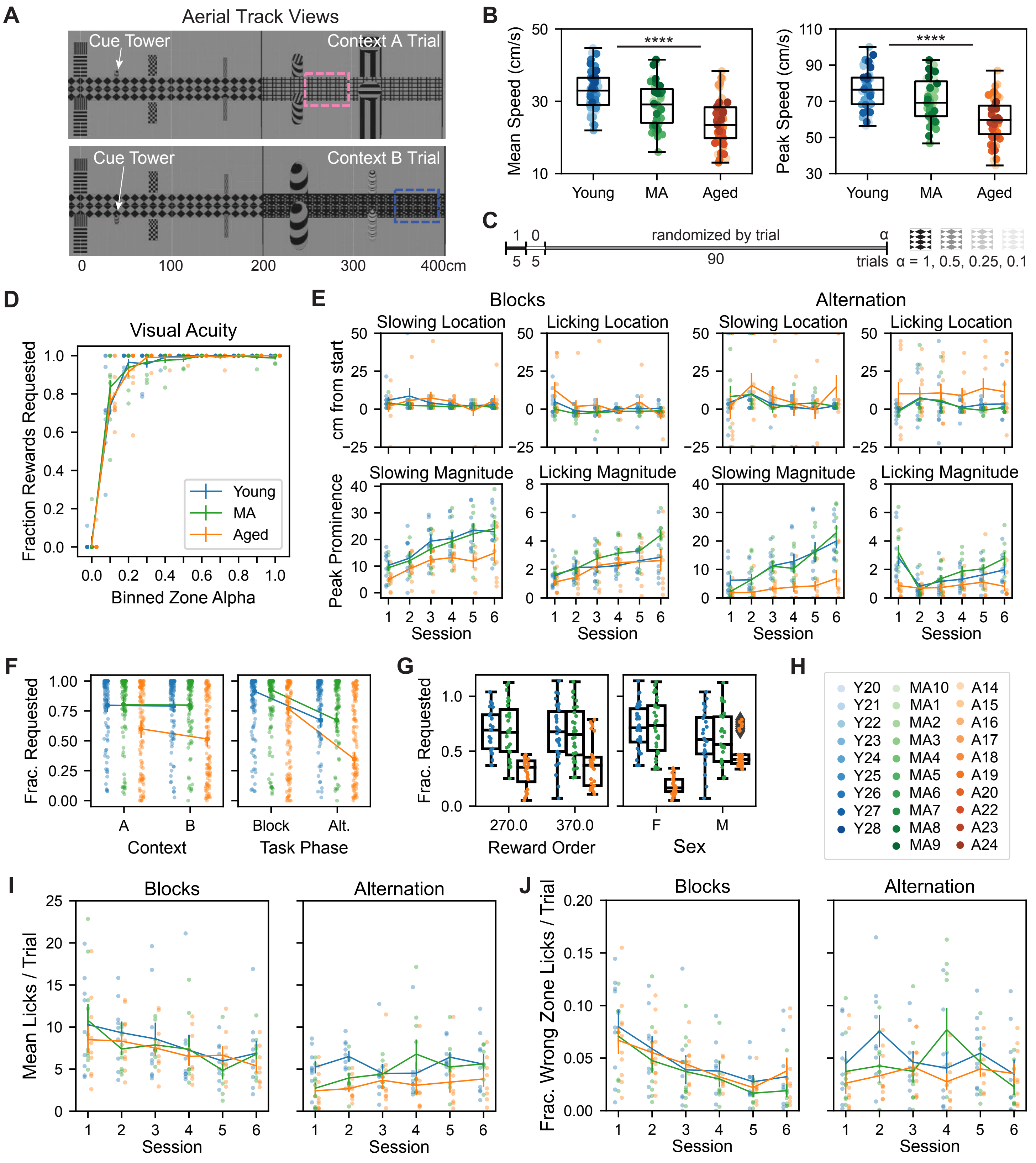
A
Aerial Track Views
Cue Tower
Context A Trial
Cue Tower
Context B Trial
0
100
200
300
400cm
B
Mean Speed (cm/s)
****
Young
MA
Aged
Peak Speed (cm/s)
****
Young
MA
Aged
C
1
0
randomized by trial
α
5
5
90
trials
α = 1, 0.5, 0.25, 0.1
D
Visual Acuity
Fraction Rewards Requested
Binned Zone Alpha
Young
MA
Aged
E
Blocks
Alternation
Slowing Location
Licking Location
Slowing Location
Licking Location
cm from start
Slowing Magnitude
Licking Magnitude
Slowing Magnitude
Licking Magnitude
Peak Prominence
Session
Session
Session
Session
F
Frac. Requested
A
B
Context
Block
Alt.
Task Phase
G
Frac. Requested
270.0
370.0
Reward Order
F
M
Sex
H
Y20 Y21 Y22 Y23 Y24 Y25 Y26 Y27 Y28
MA10 MA1 MA2 MA3 MA4 MA5 MA6 MA7 MA8 MA9
A14 A15 A16 A17 A18 A19 A20 A22 A23 A24
I
Blocks
Alternation
Mean Licks / Trial
Session
Session
J
Blocks
Alternation
Frac. Wrong Zone Licks / Trial
Session
Session

#### Figure S2

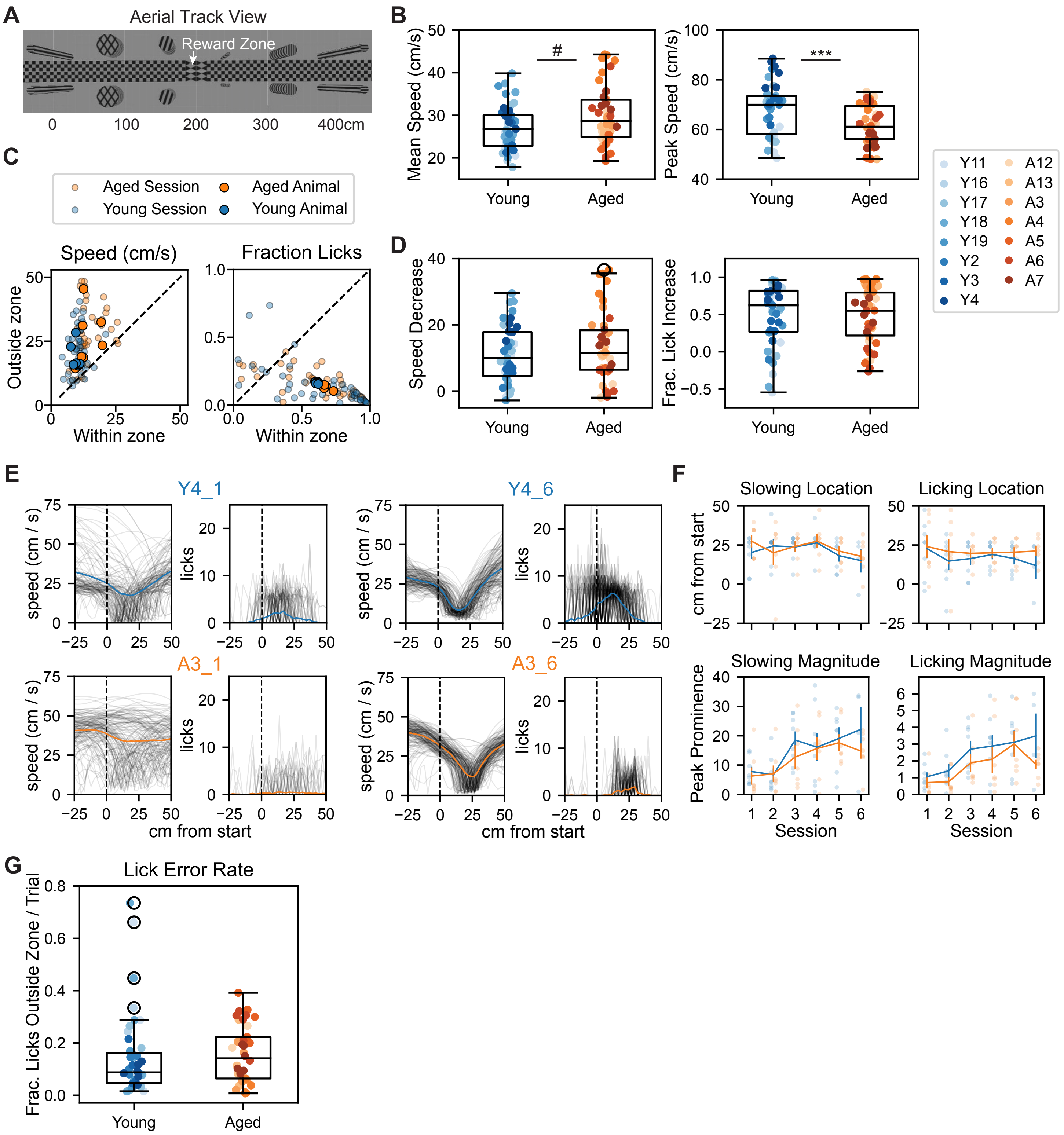
A
Aerial Track View
Reward Zone
0
100
200
300
400cm
B
Mean Speed (cm/s)
#
Young
Aged
Peak Speed (cm/s)
***
Y11
Y16
Y17
Y18
Y19
Y2
Y3
Y4
A12
A13
A3
A4
A5
A6
A7
C
Aged Session
Young Session
Aged Animal
Young Animal
Speed (cm/s)
Fraction Licks
Outside zone
Within zone
D
Speed Decrease
Frac. Lick Increase
E
Y4_1
Y4_6
A3_1
A3_6
speed (cm / s)
licks
cm from start
F
Slowing Location
Licking Location
cm from start
Slowing Magnitude
Licking Magnitude
Peak Prominence
Session
G
Lick Error Rate
Frac. Licks Outside Zone / Trial

#### Figure S3

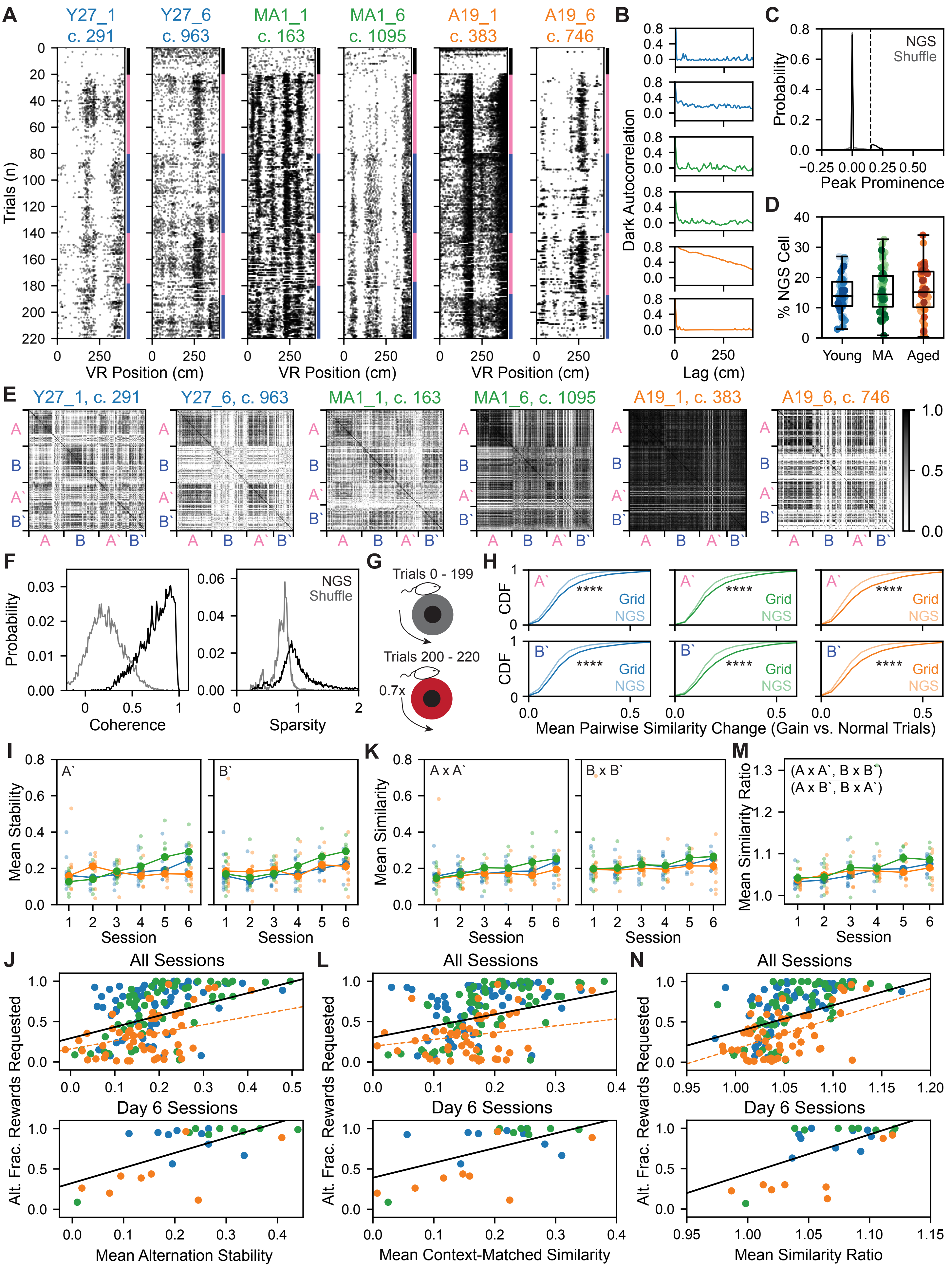

A
Y27_1 c. 291
Y27_6 c. 963
MA1_1 c. 163
MA1_6 c. 1095
A19_1 c. 383
A19_6 c. 746
Trials (n)
VR Position (cm)
B
Dark Autocorrelation
Lag (cm)
C
NGS
Shuffle
Probability
Peak Prominence
D
% NGS Cell
Young
MA
Aged
E
Y27_1, c. 291
Y27_6, c. 963
MA1_1, c. 163
MA1_6, c. 1095
A19_1, c. 383
A19_6, c. 746
F
Probability
Coherence
NGS
Shuffle
Sparsity
G
Trials 0 - 199
Trials 200 - 220
0.7x
H
CDF
Grid
NGS
****
Mean Pairwise Similarity Change (Gain vs. Normal Trials)
I
Mean Stability
Session
K
Mean Similarity
A x A`
B x B`
M
Mean Similarity Ratio
(A x A`, B x B`)
(A x B`, B x A`)
J
L
N
All Sessions
Day 6 Sessions
Alt. Frac. Rewards Requested
Mean Alternation Stability
Mean Context-Matched Similarity
Mean Similarity Ratio

**Figure S4**

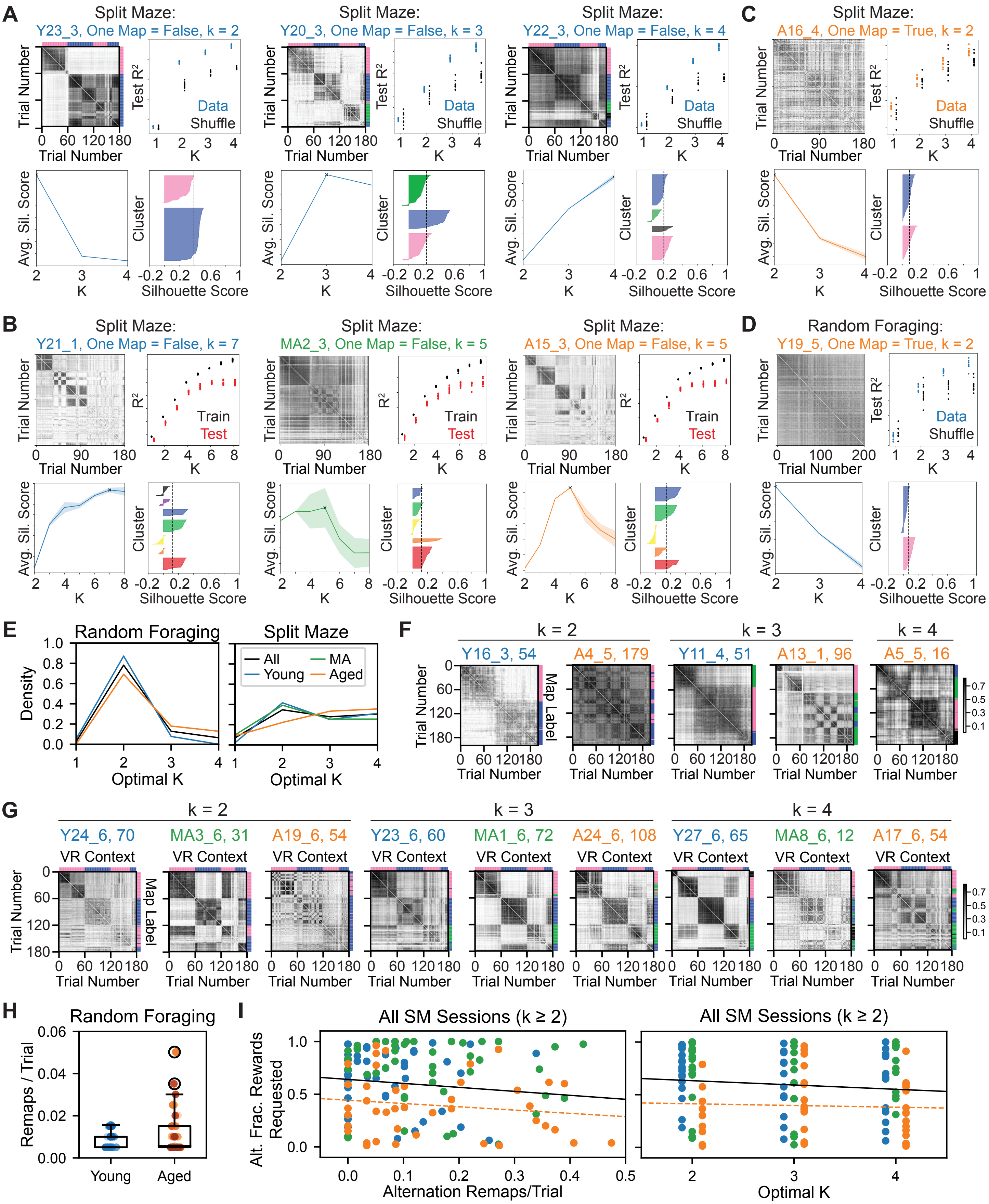
A
Split Maze:
Y23_3, One Map = False, k = 2
Split Maze:
Y20_3, One Map = False, k = 3
Split Maze:
Y22_3, One Map = False, k = 4
C
Split Maze:
A16_4, One Map = True, k = 2
Trial Number
Test R²
Data
Shuffle
K
Avg. Sil. Score
Cluster
Silhouette Score
B
Split Maze:
Y21_1, One Map = False, k = 7
Split Maze:
MA2_3, One Map = False, k = 5
Split Maze:
A15_3, One Map = False, k = 5
D
Random Foraging:
Y19_5, One Map = True, k = 2
R²
Train
Test
E
Random Foraging
Split Maze
Density
Optimal K
All
Young
MA
Aged
F
k = 2
k = 3
k = 4
Y16_3, 54
A4_5, 179
Y11_4, 51
A13_1, 96
A5_5, 16
Map Label
G
Y24_6, 70
MA3_6, 31
A19_6, 54
Y23_6, 60
MA1_6, 72
A24_6, 108
Y27_6, 65
MA8_6, 12
A17_6, 54
VR Context
H
Random Foraging
Remaps / Trial
Young
Aged
I
All SM Sessions (k ≥ 2)
Alt. Frac. Rewards Requested
Alternation Remaps/Trial
Optimal K

**Figure S5**

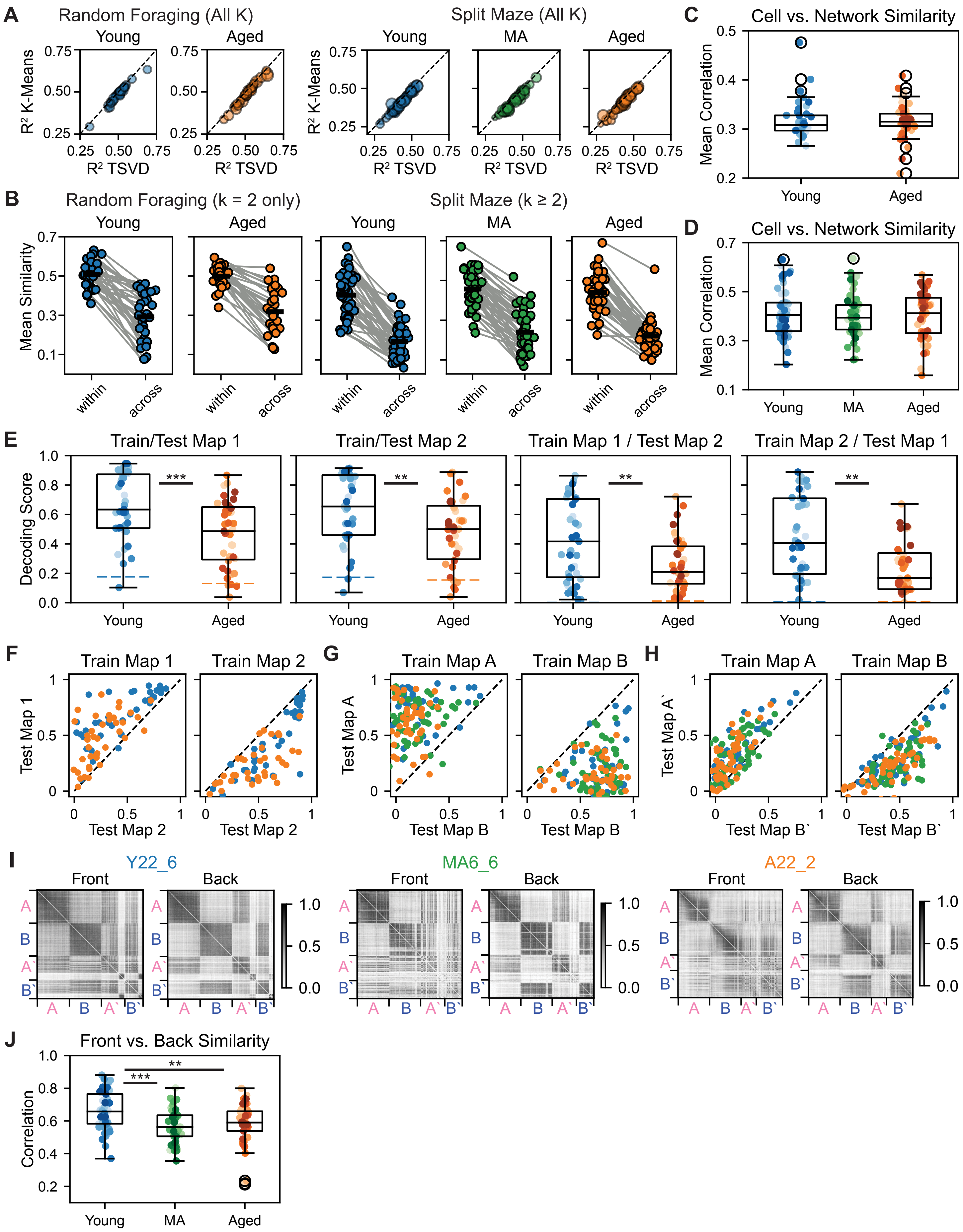
A
Random Foraging (All K)
Young
Aged
Split Maze (All K)
Young
MA
Aged
R² K-Means
R² TSVD
C
Cell vs. Network Similarity
Mean Correlation
Young
Aged
B
Random Foraging (k = 2 only)
Split Maze (k ≥ 2)
Mean Similarity
within
across
D
Cell vs. Network Similarity
Mean Correlation
Young
MA
Aged
E
Train/Test Map 1
Train/Test Map 2
Train Map 1 / Test Map 2
Train Map 2 / Test Map 1
Decoding Score
***
**
F
Train Map 1
Train Map 2
Test Map 1
Test Map 2
G
Train Map A
Train Map B
Test Map A
Test Map B
H
Test Map A`
Test Map B`
I
Y22_6
MA6_6
A22_2
Front
Back
A
B
A`
B`
J
Front vs. Back Similarity
Correlation

**Figure S6**

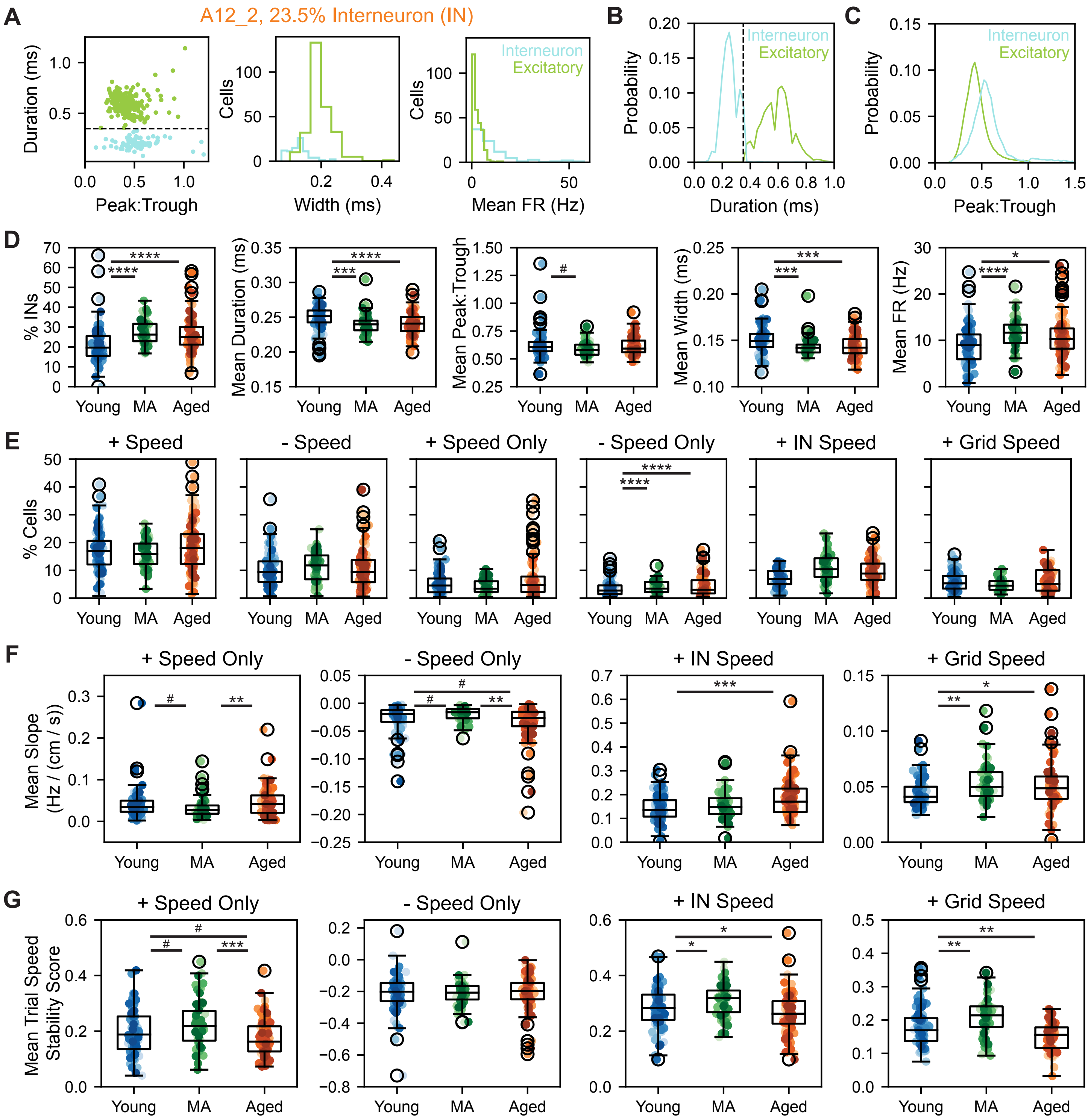

A
A12_2, 23.5% Interneuron (IN)
Duration (ms)
Peak:Trough
Cells
Width (ms)
Mean FR (Hz)
Interneuron
Excitatory
B
Probability
Duration (ms)
C
D
% INs
Mean Duration (ms)
Mean Peak:Trough
Mean Width (ms)
Mean FR (Hz)
Young
MA
Aged
E
+ Speed
- Speed
+ Speed Only
- Speed Only
+ IN Speed
+ Grid Speed
% Cells
F
Mean Slope
(Hz / (cm / s))
G
Mean Trial Speed
Stability Score

### Figure S7

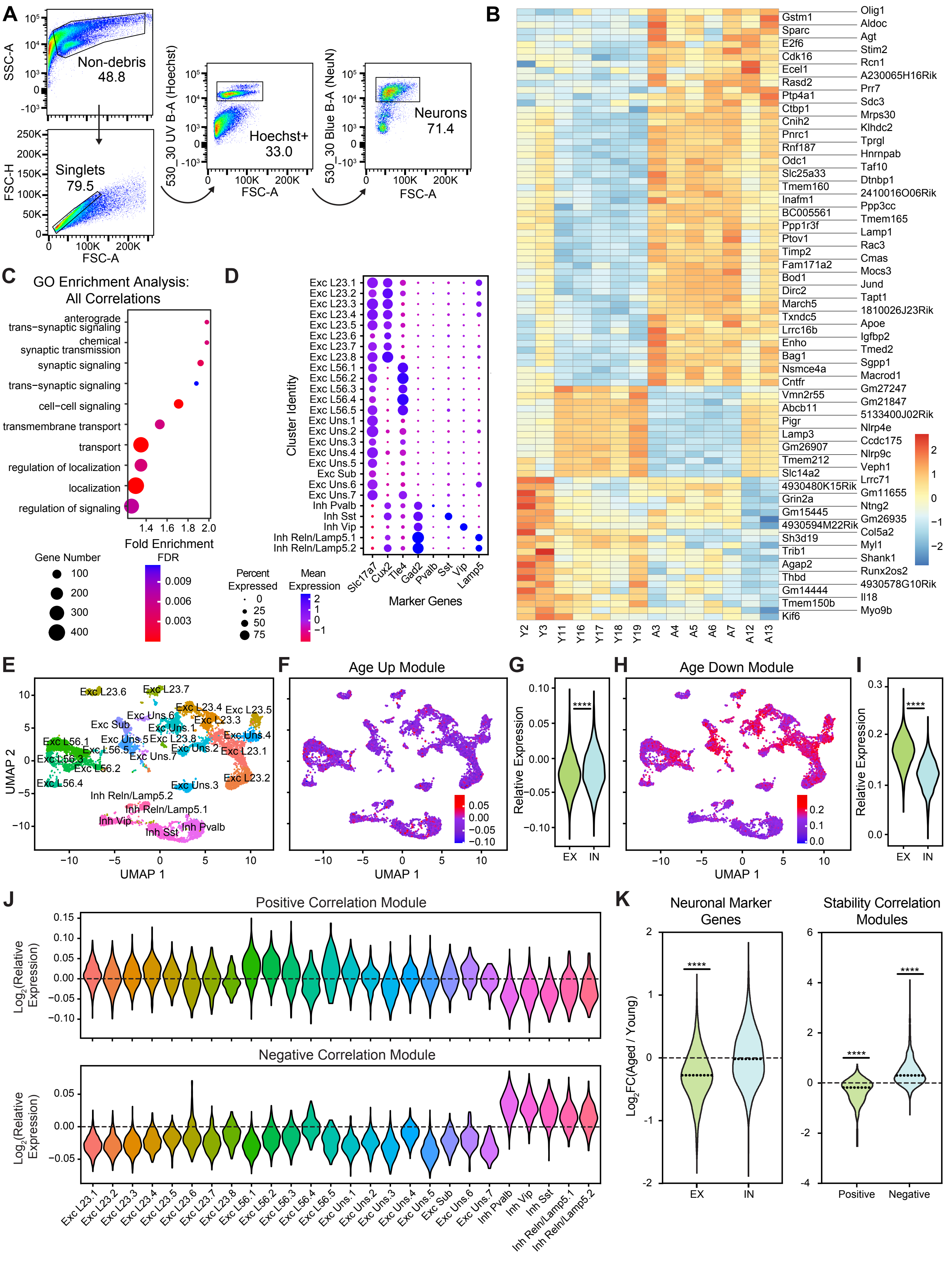

A
Non-debris 48.8
Singlets 79.5
Hoechst+ 33.0
Neurons 71.4
SSC-A
FSC-H
FSC-A
530_30 UV B-A (Hoechst)
530_30 Blue B-A (NeuN)
B
C
GO Enrichment Analysis: All Correlations
anterograde trans−synaptic signaling
chemical synaptic transmission
synaptic signaling
trans−synaptic signaling
cell−cell signaling
transmembrane transport
transport
regulation of localization
localization
regulation of signaling
Fold Enrichment
Gene Number
FDR
D
Cluster Identity
Marker Genes
Percent Expressed
Mean Expression
E
UMAP 1
UMAP 2
F
Age Up Module
G
Relative Expression
EX
IN
H
Age Down Module
I
J
Positive Correlation Module
Negative Correlation Module
Log2(Relative Expression)
K
Neuronal Marker Genes
Stability Correlation Modules
Log2FC(Aged / Young)
Positive
Negative
